# Network-based elucidation of colon cancer drug resistance by phosphoproteomic time-series analysis

**DOI:** 10.1101/2023.02.15.528736

**Authors:** George Rosenberger, Wenxue Li, Mikko Turunen, Jing He, Prem S Subramaniam, Sergey Pampou, Aaron T Griffin, Charles Karan, Patrick Kerwin, Diana Murray, Barry Honig, Yansheng Liu, Andrea Califano

## Abstract

Aberrant signaling pathway activity is a hallmark of tumorigenesis and progression, which has guided targeted inhibitor design for over 30 years. Yet, adaptive resistance mechanisms, induced by rapid, context-specific signaling network rewiring, continue to challenge therapeutic efficacy. By leveraging progress in proteomic technologies and network-based methodologies, over the past decade, we developed VESPA—an algorithm designed to elucidate mechanisms of cell response and adaptation to drug perturbations—and used it to analyze 7-point phosphoproteomic time series from colorectal cancer cells treated with clinically-relevant inhibitors and control media. Interrogation of tumor-specific enzyme/substrate interactions accurately inferred kinase and phosphatase activity, based on their inferred substrate phosphorylation state, effectively accounting for signal cross-talk and sparse phosphoproteome coverage. The analysis elucidated time-dependent signaling pathway response to each drug perturbation and, more importantly, cell adaptive response and rewiring that was experimentally confirmed by CRISPRko assays, suggesting broad applicability to cancer and other diseases.

## Introduction

Cells receive and propagate exogenous signals via receptor-mediated signaling cascades, eventually resulting in the coordinated activation and inactivation of the transcriptional programs necessary to modulate cell state in response to environmental conditions. In multicellular organisms, for instance, this allows individual cells to orchestrate the gene regulatory programs necessary to progress through lineage differentiation trajectories [1] or to respond to changes in nutrient conditions [2]. Signals originating from the interaction of secreted (autocrine), microenvironment (paracrine), and distal (endocrine) ligands, and their cognate receptors, are transmitted via complex signal transduction cascades, whose tissue specificity depends on the availability of individual protein isoforms and on their ability to form functional complexes [3].

Dysregulation of these processes plays a critical role in human disease, especially in cancer, where signaling pathway mutations represent a hallmark of tumor initiation and progression [4]. This is exemplified by colorectal cancer (CRC), where progression from normal cells in the intestinal crypt to adenocarcinoma is determined by progressive accrual of genetic and epigenetic alterations in key signaling pathways, ultimately resulting in transformation [5]. Critically, despite similar histological presentation, we and others have shown that different CRC subtypes exist, due to signaling pathway-mediated integration of heterogeneous mutational landscapes [5], resulting in aberrant activation/inactivation of small Master Regulator protein modules [6]. Yet, the specific signaling mechanisms leading to concerted, aberrant activity of these regulatory modules and causally responsible for their time-dependent response and adaptation to drug perturbations are still largely elusive.

While their elucidation may provide more universal insights into tumor dependencies and response to treatment [6], systematic, proteome-wide elucidation of tissue-specific signaling networks has trailed the study of regulatory interactions and still represents one of the hallmark challenges in systems biology, with potential applications to both basic and translational research.

Signal transduction is mediated by reversible post-translational modifications (PTMs), often responsible for a rapid on/off switch in protein activity or ubiquitin-mediated proteasomal degradation. Among these, phosphorylation represents the most frequently studied event, due to its profound impact on protein conformation and function. In human cells, protein phosphorylation and de-phosphorylation is mediated by *>* 500 kinases [7] and *>* 200 phosphatases [8], respectively (KP-enzymes in the following). Although these enzymes have substrate specificity, determined by low to medium-affinity peptide-binding domains (PBDs), many substrates can be processed by multiple, sometimes closely related enzymes, resulting in considerable cross-talk.

Auto-regulatory feedback loops, sub-cellular localization mechanisms, and context-specific availability of the cognate binding partners necessary for formation of active complexes further increase the complexity of these biological processes.

Enzyme-Substrate (ES) interactions have been broadly studied, including via low-throughput biochemical assays and structure determination [9], as well as by high-throughput methods using array-based [10], affinity purification coupled to mass spectrometry (AP-MS) [11, 12], and computational biology approaches [13, 14]. As a result, established repositories of ES interactions have been assembled, such as PhosphoSitePlus [15] and Pathway Commons [16], among others. However, none of these repositories addresses the context specific nature of ES interactions and only comprise a small fraction of the total number of such molecular interactions.

Furthermore, these have been studied at steady state, thus potentially failing to provide critical insight into the time-dependent signaling processes that underlie cell adaptation to endogenous and exogeneous perturbations.

A handful of reverse engineering methods for the mechanism-based interrogation of signaling pathways have been proposed, such as pARACNe (phospho-ARACNe) [17], KSEA (Kinase Substrate Enrichment Analysis) [18], INKA (Inference of Kinase Activity) [19], or PHONEMeS (PHOsphorylation NEtworks for Mass Spectrometry) [20]. However, in terms of accuracy and sensitivity, they still significantly trail behind equivalent methods for the dissection of regulatory networks [21].

To address these challenges, we developed VESPA (Virtual Enrichment-based Signaling Protein-activity Analysis)—a novel phosphoproteomic-based machine learning methodology for the dissection of ES interactions and for measuring signaling protein activity—and have applied it to study post-translational cell adaptation mechanisms that mediate CRC’s resistance or lack of sensitivity (*i.e.*, *insensitivity*) to clinically-relevant targeted drugs. Our proposed methodology provides four critical elements of novelty, including: (i) the ability to reconstruct and interrogate disease context-specific signaling networks *de novo*, based on phosphoproteomic profiles, such as those made available by the Clinical Proteomic Tumor Analysis Consortium (CPTAC) [22], (ii) the ability to measure the activity of signaling enzymes, including those that are poorly characterized in the phosphoproteomic profiles, based on the phosphorylation state of their substrates, (iii) the ability to deconvolute the time-dependent response of cancer tissues to inhibitors targeting signaling enzymes, a challenging task since the phosphostate of their high affinity targets is not generally affected by treatment, and (iv) the ability to identify potential mechanisms presiding over drug resistance and cell adaptation. Systematic benchmarking, based on ES reference databases, assessing differential KP-enzyme activity of primary drug targets in cell lines with experimentally validated sensitivity to *>* 200 targeted inhibitors, showed that VESPA substantially outperforms established approaches.

In a proof-of-concept application, we designed a large-scale drug perturbation experiment and used VESPA to elucidate the molecular mechanisms of CRC adaptation to drug treatment that mediate resistance or insensitivity in a highly context-specific fashion. For this purpose, we analyzed a recent CPTAC CRC-specific proteogenomic dataset to yield a comprehensive CRC context-specific signaling network for 371 kinases/phosphatases. SigNet interrogation further effectively assessed kinase/phosphatase activity across the CPTAC-CRC cohort, resulting in identification of clinical subtypes associated with aberrant enzyme activity, which allowed us to select a representative cell line panel as experimental model system. Using a highly scalable Data-independent Acquisition (DIA) [23] proteomic workflow [24–26], we then acquired 336 profiles from these six subtype-matched CRC cell lines perturbed with seven clinically-relevant drugs, at seven time points, ranging from 5min to 96h, providing the data foundation for our study.

VESPA analysis provided novel insight into the ability of CRC cell lines to adapt and “rewire” their signaling networks following drug perturbation. Critically, this revealed how specific cells implement similar drug responses yet over highly different timeframes, while others present highly idiosyncratic response mechanisms. Moreover, for drug resistant cells, this allowed us to identify signaling proteins responsible for the progression from initial drug perturbation to development of resistance. To assess its predictive nature, we experimentally validated VESPA’s ability to identify candidate, cell line-specific, resistance mechanisms by performing CRISPR-knock-out experiments, targeting all human phospho-enzymes in drug treated cell lines. Our analysis shows that VESPA’s predictions are indeed enriched in proteins that synergize with drug treatment in resistant cell lines, thus suggesting potential value towards identification of potential combination therapy opportunities.

## Results

### Conceptual Workflow

VESPA comprises two-steps. First, a *dissection* step (dVESPA) reconstructs context-specific signaling networks, *de novo*, from phosphoproteomic and whole-proteome profiles of large-scale tumor cohorts (Fig. 1a). Such datasets—often comprising *>* 100 samples—are now broadly available, having been generated for many cancer subtypes by initiatives such as CPTAC. VESPA-inferred Signal Transduction Networks (SigNets) are unique in that they fully recapitulate the tumor context-specific nature of ES interactions, as well as their directionality and statistical confidence.

**Figure 1.**
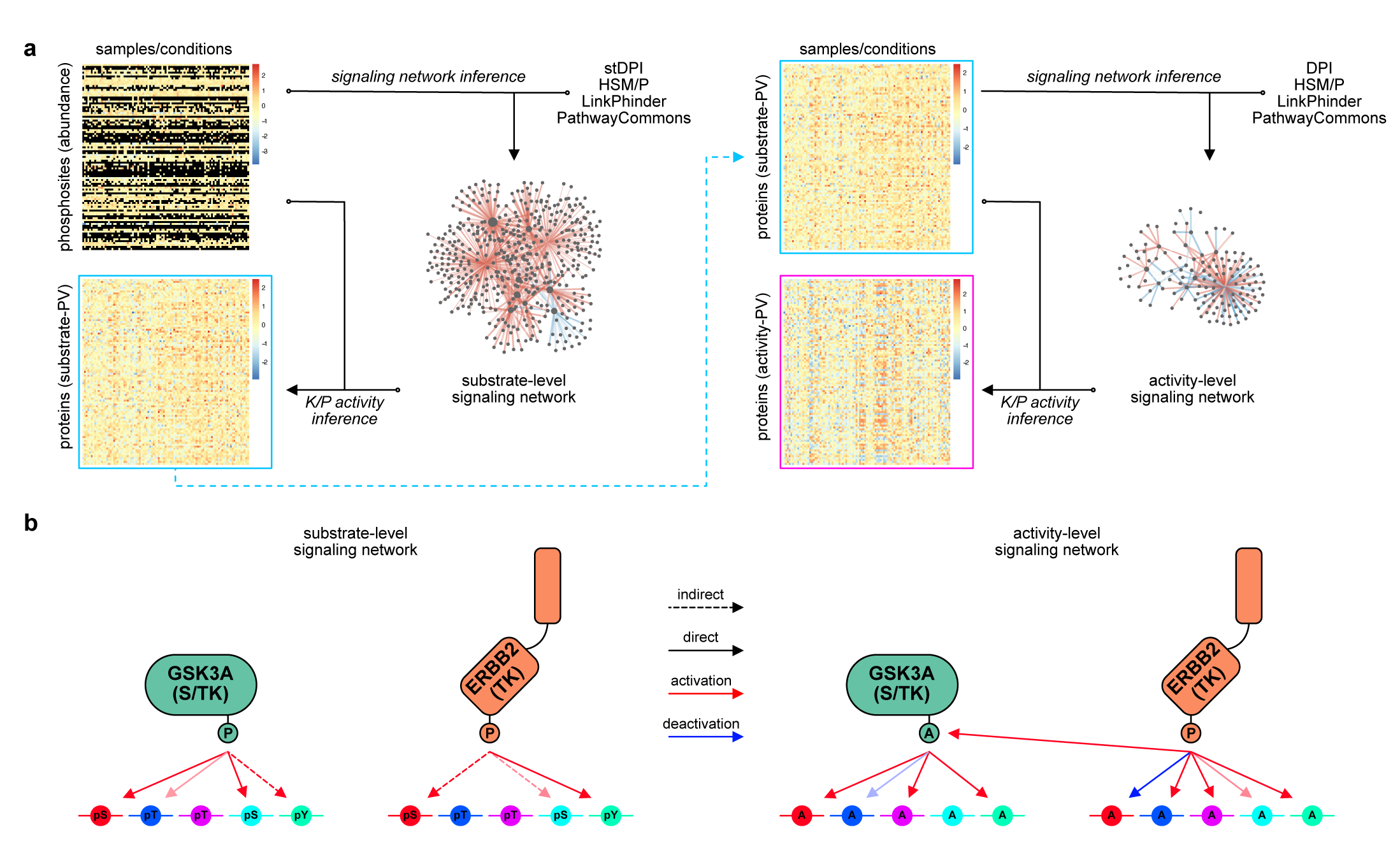
Methodological overview of VESPA. **a)** VESPA infers protein signaling activity for kinases and phosphatases on substrate-level (blue box, left panel) and activity-level (pink box, right panel). A quantitative matrix of phosphopeptide or phosphosite abundance across samples or conditions including missing values (black) represents the main input for VESPA. The signaling network reconstruction module uses this matrix together with optional priors from reference networks to assess regulatory relationships by computing the mutual information between enzymatic regulator and target phosphopeptides or phosphosites and the signal transduction Data Processing Inequality (stDPI) to generate signalons for each regulator consisting of interaction probabilistic weight and mode of regulation (kinase activation: red, phosphatase deactivation: blue) with substrate targets. VESPA then uses the substrate-level signalons to infer substrate-level kinase/phosphatase activity (blue box). This quantitative matrix represents the input for activity-level signaling network reconstruction, which uses the Data Processing Inequality (DPI) to generate more abstract and generalized signalons, which are then in turn used to infer protein signaling activity on activity-level (pink box). **b)** Methodological differences between substrate-and activity-level signaling networks. On substrate-level, ST-Ks (e.g. GSK3A, green) are primarily associated with direct phosphorylation targets, whereas TKs (e.g. ERBB2, orange) can frequently not be directly associated with (unenriched) tyrosine-phosphorylated sites. On activity-level, more abstract “activation/deactivation” events can better associate targets for both ST-Ks and TKs.

In a second step (mVESPA), SigNets are used to *measure* differential KP-enzyme activity in individual samples, based on differential substrate phosphorylation (*signalon*) compared to a reference sample (Fig. 1b). For instance, when assessing the activity of signaling proteins following drug treatment, the reference sample would be the same tissue treated with vehicle control media. To infer enzyme activity, mVESPA leverages a probabilistic framework that integrates the differential phosphorylation state of its substrates, while accounting for potential confounding effects by other enzymes with which it may share some substrates (cross-talk). To improve performance for serine/threonine kinases (ST-Ks)—especially from low phosphoproteomic profile coverage—and to improve substrate coverage of tyrosine kinases (TKs), without requiring immunoprecipitation (IP) based enrichment methods, VESPA leverages a two-step hierarchical approach. Activity profiles are first generated in coarse fashion, based on the phosphorylation state of KP-enzyme’s substrates, and are then used to refine the original activity estimates.

Despite a superficial similarity of these steps to algorithms designed for the study of transcriptional networks, such as ARACNe [27, 28] and VIPER [29], there are critical differences that were necessary to account for the unique structure and sparseness of phosphoproteomic profiles. These are summarized in the following.

#### Substrate inference

To measure the likelihood of an enzyme/substrate interaction, dVESPA extends the ARACNe algorithm [27, 28] to phosphoproteomic data (see Methods). dVESPA is designed to leverage continuous peptide intensities, as obtained from quantitative proteomic workflows [30], using a novel hybrid partitioning approach to compute mutual information (hpMI). This effectively addresses challenges associated with missing values due to censoring [31, 32], a critical issue in bottom-up phosphoproteomic analyses (Supplemental Fig. 1a, Methods). Second, to eliminate indirect ES interactions, dVESPA implements a signal transduction-specific version of the Data Processing Inequality (stDPI) theorem, which accounts for the full topology of three-way signaling interactions, as determined by the enzymatic properties of kinases and phosphatases (Supplemental Fig. 1b, Methods). When phosphoproteomic profile coverage is insufficient to reconstruct KP-enzyme substrate relationships, dVESPA can also incorporate priors from reference databases—such as Pathway Commons [16], LinkPhinder [14] or the Hierarchical Statistical Mechanistic model (HSM) [13]—to generate hybrid signalons that incorporate both context-free and context-specific information. Critically, each inferred interaction has an associated statistical *confidence (p-value)* and *directionality,* as determined by the enzymatic function of the signaling protein, which are also inferred by dVESPA (Methods).

#### Cross-talk correction

mVESPA leverages the Rank-based Enrichment Analysis (aREA)-framework [29]—a probabilistic implementation of gene set enrichment analysis [33]—to address potential confounding effects arising from enzymes sharing common substrates. This allows incorporating the shadow analysis [34] and pleiotropy correction [29] methods that were originally designed to address this issue in gene regulatory networks (see Methods).

#### Site-specific activity inference

Kinase and phosphatase activity can then be measured by assessing the phospho-state of their substrates, either at the *whole-protein* level—i.e., by integrating the state of all phosphosites—or at the *phosphosite-specific* level (Methods). The latter can help elucidate specific phosphosites that critically contribute to a protein’s enzymatic activity. Indeed, proteins generally comprise multiple phosphosites with diverse and potentially opposite contributions to their enzymatic and regulatory activity—ranging from modulating enzyme abundance via ubiquitylation pathways to mediating dimerization or conformational changes necessary to support enzymatic activity—as well as sites providing no measurable contribution. To address this issue, mVESPA infers both phosphosite-specific and whole-protein signalons (Methods).

#### Hierarchical activity and model inference

Some KP-enzymes substrates, including TK substrates, are only sparsely represented within phosphoproteomic profiles. As a result, unless they are enriched—*e.g.*, by pull-down with phosphosite-specific antibodies—they cannot be effectively identified as substrates, resulting in low-quality signalon generation and TK activity prediction. To address this challenge, mVESPA implements a two-step, hierarchical activity inference approach (Fig. 1, Supplemental Fig. 1c-d, Methods). In the first step (*substrate-level SL-analysis*) coarse-level KP-enzyme activities are assessed from phospho-state of dVESPA-inferred substrates (Fig. 1a). Both substrate identification and KP-activity are then refined in a second step (*activity-level AL-analysis*) by leveraging the coarse-level activity of candidate substrates, rather than their phospho-state (Methods). The rationale is that many TK substrates are themselves ST-Ks, whose activity can be inferred quite accurately by substrate-level analysis. Consistently, we show that this two-step approach also improves KP-enzyme signalon and activity reproducibility, when using phosphoproteomic profiles with increasingly lower depth, because lower depth decreases substrate coverage (Fig. 1b). Results from the two-step analysis are then integrated, for both ST-Ks and TKs, using Stouffer’s method, since candidate substrate’s activity and phosphostate are assessed from independent data and are thus statistically independent (Methods).

#### Signalon optimization

If multiple signalons for the same KP-enzyme were generated from independent datasets, mVESPA selects the one that is most informative, based on the overall significance of differentially active KP-enzymes, as implemented by the metaVIPER algorithm [35] (Methods).

### Generating a CRC context-specific SigNet

Kinase inhibitors and other drugs inhibit the activity of their targets without affecting their phospho-state but rather by binding to the protein’s active site. As a result, drug target identification from phosphoproteomic profiles is challenging. However, availability of accurate and comprehensive SigNets allows direct assessment of target activity, independent of their phosphorylation state. To study signal transduction in CRC, we availed ourselves of proteomic and phosphoproteomic profiles from three datasets. The first comprises 97 CRC samples from the Clinical Proteomic Tumor Analysis Consortium (NCI/NIH), here referred to as CPTAC-S045 [36]. As second dataset, we normalized the CPTAC-S045 phosphosite abundances by the corresponding whole protein abundances, which would allow to identify confounded KP→S relationships, as suggested previously [37] (CPTAC-S045N). In addition, we generated a third dataset, comprising phosphoproteomic profiles from six CRC cell lines (HCT-15, HT115, LS1034, MDST8, NCI-H508 and SNU-61) at three time points (1h, 24h, 96h), following perturbation with a collection of seven clinically relevant drugs and vehicle control media (144 samples, here referred to as U54-NET dataset, see below).

We used dVESPA to dissect independent SigNets from these datasets, using the stDPI/DPI method to select the most direct interactions (Supplemental Data 1, Methods). For each kinase and phosphatase, one to three site-specific signalons were generated, depending on whether the phosphosite (or protein) was measured and significant interactions were identified in each dataset. Site-specific signalons were then aggregated for each dataset separately to a protein-level representation, and the most representative of the three possible signalons was selected, generating the best response for a set of target phosphoproteome profiles (Methods). Overall, consistent with the KP-enzyme fraction expected to be expressed in any specific cellular context, signalons comprising ≥ 5 candidate substrates could be reliably inferred for 51.0% of human KP-enzymes, from at least one of the datasets. The first step (SL-analysis) produced an initial SigNet comprising 163,313 interactions, representing the activity of 283 kinases and 88 phosphatases on 7,727 candidate substrates. The second step (AL-analysis) improved the model by identifying 16,309 additional interactions, representing the activity of 187 kinases and 37 phosphatases on 371 candidate substrates. To support more mechanistic studies, we also generated a phosphosite-level network comprising interactions representing 1,649 individual phosphosites, whose differential phosphorylation could be assessed from available datasets. By collapsing phosphosites in the same peptide-binding domain, which are frequently correlated in both phospho-state and functional role—a refined set of 918 non-redundant phosphosites was generated and used for this analysis (Methods). Each KP-enzyme/substrate interaction was characterized in terms of both *mode of regulation* (i.e., substrate activation or deactivation by kinases and phosphatases, respectively) and likelihood.

The CPTAC datasets provided a more comprehensive phosphosite representation than those from cell line perturbations—*i.e.*, 31,339 phosphosites from CPTAC-S045 *vs.* 13,529 from U54-NET, respectively—as expected due to the lower genetic background variability in the handful of selected cell lines and the different analytical depth of the proteomic data acquisition methods (DDA-TMT *vs.* DIA-LFQ). However, U54-NET interactions were often selected as more informative by signalon optimization analysis (Methods). Indeed, at the substrate-level, 47.2%, 43.4% and 9.4% of the optimized signalons were derived from CPTAC-S045, U54-NET and CPTAC-S045N dataset, respectively. Dataset specificity was even more skewed at the activity-level analysis, where U54-NET accounted for 46.4% of the optimized signalons, with CPTAC-S045 and CPTAC-S045N accounting for 38.4% and 15.2% of them, respectively.

A key advantage of the proposed methodology is that, once a SigNet is available, activity can be measured even for KP-enzymes whose phospho-state is undetected in aspecific dataset. Indeed, VESPA could measure enzymatic activity for 158 of 371 (42.6%) of all KP-enzymes in the CRC SigNet that completely lacked phospho-state information thus almost doubling the amount of critical available information.

Furthermore, multiple dataset integration can effectively combine DIA’s high throughput with the more comprehensive nature of the fractionated CPTAC profiling. Overall, despite the well-known sparseness of peptides and phosphopeptides detected by proteomic assays, mVESPA quantitatively assessed the activity of 371 KP-enzymes—*i.e.*, around half of all known human KP-enzymes and, likely, an almost complete subset of the KP-enzymes expressed in CRC cells.

### Benchmarking the MI estimator

With up to 20-80% missing values in typical phosphoproteomic profiles, phosphopeptide-based MI estimation is challenging. We thus benchmarked the performance of the novel hybrid-partitioning mutual information metric (hpMI) using the U54-NET dataset and compared it to either removing proteins with missing data or imputing (random low intensity noise) profiles. Specifically, we generated sparser instances of the quantitative matrix by removing up to 80% of the data, in 20% increments (see Methods) and assessed the impact on MI estimation—including depleted MI (dMI), imputed MI (iMI) and hpMI estimators (Fig. 2a). This was accomplished by computing the recovery of the highest-likelihood interactions, as predicted by the Hierarchical Statistical Mechanics (HSM) [13] algorithm (Methods). As previously described [27], MI estimates were statistically validated using a bootstrapped null model and filtered at 5% FDR. Although for some well-sampled, highly correlated KP→S pairs, both dMI and iMI measured a statistically significant MI, hpMI inferred 102.4% more correct ES interactions than dMI, and 31.3% more correct ES interactions than iMI from sparsely covered interactions (*>*20% complete), particularly in case of lower (*ρ <* 0.5) KP→S correlation (Fig. 2a).

**Figure 2.**
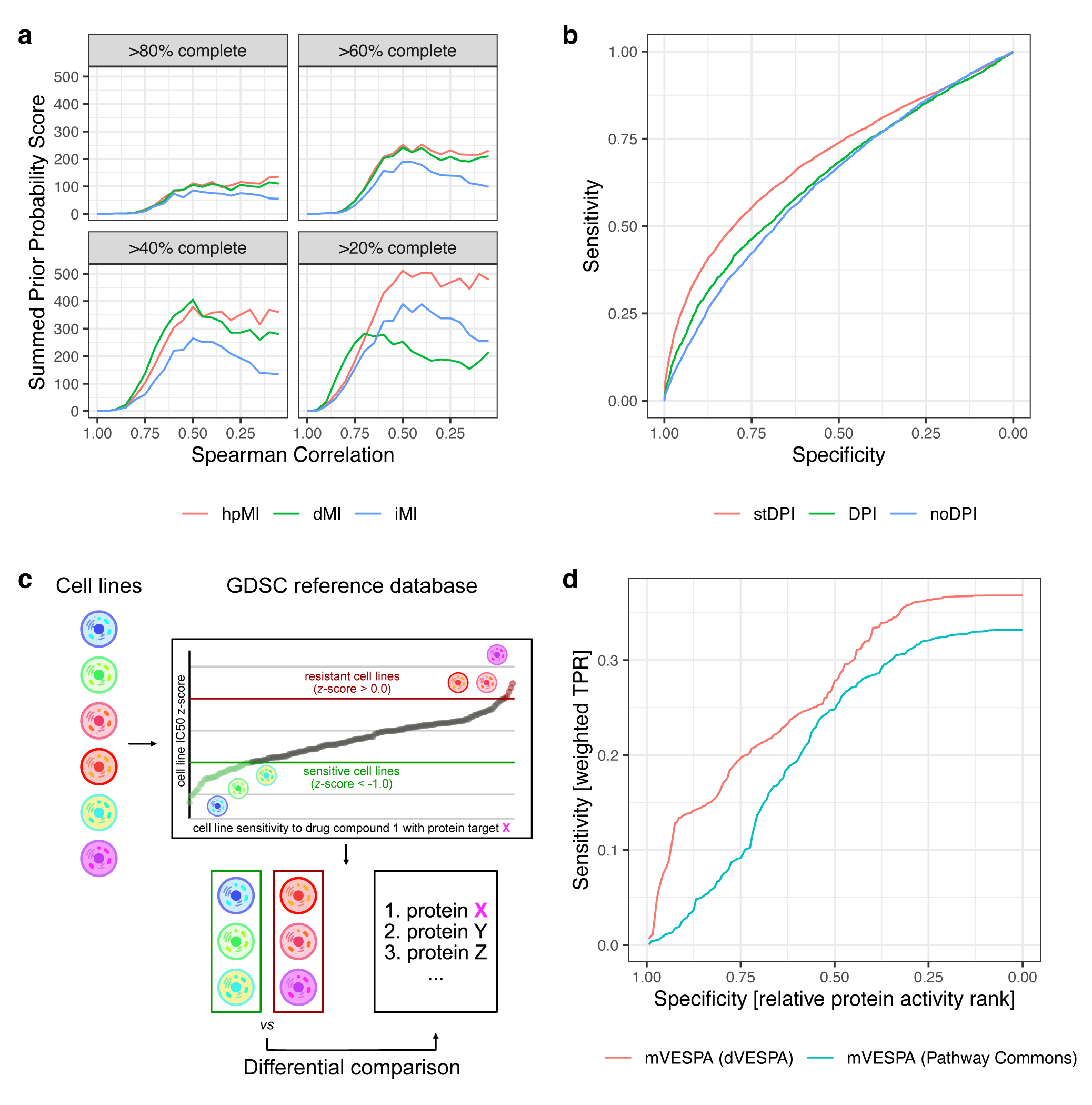
Benchmark and validation of dVESPA and mVESPA. **a)** Comparison of different mutual information (MI) estimation strategies based on imputation (iMI), depletion (dMI) and hybrid partitioning (hpMI) and the MI– Spearman correlation relationship using the CPTAC-S45 dataset. **b)** Receiver operating characteristic of (signal transduction) Data Processing Inequality (stDPI/DPI) and unprocessed (noDPI) signaling dependencies as evaluated using a ground truth dataset. **c)** Baseline profiles of six diverse CRC cell lines were acquired and used with the GDSC reference database to identify sensitive and resistant or insensitive cell lines for each covered drug compound. Using mVESPA and CRC-specific signalons, differential comparisons were conducted for each drug compound to identify the top differentially active regulators. **d)** VESPA (red), consisting of mVESPA and dVESPA, performs substantially better than mVESPA using context unspecific Pathway Commons (blue) SigNets.

### Benchmarking indirect interaction removal

To eliminate indirect interactions (i.e., those due to a cascade of distinct direct interactions, KP→KP’→S), dVESPA uses a *signal transduction-adapted* version of the Data Processing Inequality (stDPI/DPI) originally proposed in [27, 28] (Supplemental Fig. 1b, Methods). The DPI states that, in any system where information is not perfectly transferred (lossy)—thus including virtually all molecular networks—direct information transfer (i.e., KP→S) is always greater than indirect information transfer (KP→KP’→S), thus allowing effective removal of indirect interactions.

To benchmark whether the stDPI improves removal of indirect interactions, including compared to the original DPI formulation, we first generated a gold-standard dataset for ST-K proteins using the Hierarchical Statistical Mechanics (HSM) [13] algorithm (Methods). Specifically, ground truth interactions were selected based on HSM analysis of domains identified as primary determinants of ST-K→phosphopeptide specificity, including PDZ, SH3, WH1, and WW domains. As a negative gold standard, we used HSM predicted TK→S interactions, based on PTB, PTP and SH2 domains, since the dataset used for this benchmark (U54-NET) is not enriched for phosphotyrosine peptides and should thus not support their identification.

We then analyzed the U54-NET dataset using dVESPA to produce an SL-analysis-based SigNet with each of the three DPI options—i.e., no DPI, regular DPI and stDPI—and compared the inferred interactions with the gold standard datasets (Methods). Receiver operating characteristics (ROC) curves show that stDPI significantly outperforms the other two options (stDPI *vs.* no DPI: *p*-value *<* 2.2e-16, stDPI *vs.* DPI: *p*-value *<* 2.2e-16; area under the curve (AUC): noDPI = 0.626 DPI = 0.640, stDPI = 0.693) (Fig. 2b, Methods).

Taken together, these data show that hpMI and stDPI—two novel phosphoproteomic-specific component of dVESPA—significantly improve KP→S inference.

### mVESPA Benchmarking and validation

To benchmark mVESPA predictions, we extended a strategy previously used to infer kinase activity in cell lines [19]. The Genomics of Drug Sensitivity in Cancer project (GDSC) has assessed sensitivity of *>*1,000 human cancer cell lines to hundreds of small molecule compounds, including high-affinity kinase inhibitors [38]. When combined with a curated list [19] of the primary (i.e., high-affinity) targets of each inhibitor, this resource can be used to effectively assess relative kinase activities because—within a specific tumor type and barring adaptive resistance mechanisms—higher activity of the drug target should correlate, on average, with increased sensitivity to its high-affinity inhibitor(s). Thus, the statistical significance of the correlation between predicted differential kinase activity and cognate inhibitor sensitivity, across multiple cell lines, provides an effective benchmark to assess relative predictive power between different protein activity prediction algorithms [19].

For the benchmark, we used the mVESPA-predicted activity of protein kinases representing high-affinity targets of GDSC-tested kinases, as assessed from baseline (*i.e.*, not perturbed) phosphoprofiles of six CRC cell lines, in triplicate (U54-BL, Methods). To support comparative analysis, we modified the benchmark [19] to use pairwise differential rather than absolute target protein activity ranks, in sensitive *vs.* resistant cell lines, and the analysis was performed independently for each cell line (Fig. 2c).

Indeed, for each drug, GDSC previously identified sensitive (low z-score) and resistant (high z-score) cell lines, based on the compound’s log(IC_50_) as measured across 1,000 cell lines. For this benchmark, we thus selected the compounds eliciting the greatest differential sensitivity (sensitive cell line z-score *<* -1.0, resistant cell line z-score *>* 0.0) across the pairwise combinations of the six CRC cell lines for which phosphoproteomic profiles were available (Figure 2d, Methods). For each selected drug, differential target kinase activity was measured by mVESPA in sensitive *vs.* resistant cells, using the CRC SigNets (Fig. 2c, Methods). Finally, we assessed the method’s sensitivity using a score represented by the inferred differential activity of the i-th protein (*DP _i_*) weighted by the cell line sensitivity to the associated inhibitor (*w_i_*) and integrated over the top *n* most differentially active proteins (*S*(*n*) = *w_i_DP _i_*) (Methods).

First, we assessed the differences in mVESPA performance when using signalons from either dVESPA analysis (*i.e.*, context-specific) or from other sources, including generalized and contextualized reference databases, by plotting *S*(*n*) as a function of *n*, normalized to the total number of covered proteins, resulting in pseudo-ROC curves (Methods). For this comparison, either only the overlapping (intersection) or complete (full) sets of kinases were compared (Methods), where the intersections provide a measure of accuracy of predictions and the full sets provide a measure of network coverage. The analysis shows that dVESPA significantly outperformed the generalized reference database Pathway Commons, for both ST-Ks and TKs activity inference (intersection: *p*-value *<* 0.001, full set: *p*-value *<* 0.023) (Fig. 2d, Supplemental Fig. 2a-3a, Supplemental Tables 1-2, Methods). Furthermore, the benchmark shows that stDPI/DPI-based removal of indirect interactions in dVESPA generally provided equal or slightly higher accuracy but better comprehensiveness compared to using contextualized LinkPhinder (LP) [14] (intersection: *p*-value *<* 0.156, full set: *p*-value *<* 1.9e-6) or Hierarchical Statistical Mechanistic model (HSM) [13] (intersection: *p*-value *<* 0.003, full set: *p*-value *<* 4.4e-4) reference networks. Assessing the performance for TK activity inference, stDPI/DPI only improved in terms of comprehensiveness over LP (intersection: *p*-value *<* 0.580, full set: *p*-value *<* 4.7e-4) but not HSM (intersection: *p-* value *<* 0.766, full set: *p*-value *<* 0.947) (Methods, Supplemental. Fig. 2b-3b, Supplemental Tables 1-2).

We then benchmarked performance differences associated with each mVESPA component, including (a) signalon integration and optimization across multiple dataset (Supplemental Fig. 2c-3c, Supplemental Tables 1-2), (b) differences between substrate-level, activity-level and integrated analysis (Supplemental Fig. 2d-3d, Supplemental Tables 1-2), and (c) the effects of cross-talk correction (Supplemental Fig. 2e-3e, Supplemental Tables 1-2).

These analyses show the effect of both each individual improvement of mVESPA as well as their cumulative effect, the latter resulting in the best overall performance and asignificant improvement over the current state-of-the-art (Fig. 2d). Based on these results, we selected the stDPI for SL and regular DPI for AL signalon inference, respectively, followed by integration using Stouffer’s method, for all subsequent studies (Methods).

### CRC cell line selection and representation

To study CRC-specific drug mechanisms using phosphoproteomic profiles, we proceeded to identify cell lines representing high-fidelity models of established CRC subtypes for pharmacological perturbation. Since comprehensive phosphoproteomic data for all CCLE cell lines was not available, CRC subtype selection was based on a recent analysis of the TCGA CRC cohort, stratifying transcriptional state and genetic alterations into 8 distinct clusters [39]. We thus identified cell lines from the Cancer Cell Line Encyclopedia (CCLE) [40] representing high-fidelity surrogates of each subtype based on the overlap of the most differentially active proteins in patient derived samples and cell lines (OncoMatch algorithm [41, 42], *p* ≤ 10^-5^, Methods), as assessed by VIPER analysis. We use the term “high-fidelity” surrogate because we have shown that OncoMatch can identify cancer models, including cell lines and mouse models, that recapitulate patient-relevant drug response [42]. Considering also criteria related to optimal growth in culture and use in a high-throughput microfluidic setting, the analysis identified six high-fidelity models, including HCT-15, HT115, LS1034, MDST8, NCI-H508 and SNU-61, representing 5 out of 8 previously identified CRC clusters with at least one top-5 ranking cell line for each cluster, based on OncoMatch, and all 8 cell lines with at least one top-10 ranking cell line (Supplemental Fig. 4).

Given this initial selection, we then proceeded to assess whether the selected cell lines were also representative of subtypes identified by phosphoproteomic cluster analysis by comparing their KP-enzyme differential activity profiles to those of subtypes identified by cluster analysis of the 97 clinically annotated CRC samples in the CPTAC-S045 cohort [36]. For this purpose, we acquired baseline phosphoproteomic profiles of each cell line in triplicate by label-free DIA. At 1% peptidoform and protein FDR, the analysis identified and quantified the state of 9,813 phosphosites on 18,012 unique peptide precursors from 3,320 proteins (Methods). We refer to this dataset as “U54-BL”. At the peptide precursor-level, the dataset/matrix completeness—i.e., the fraction of runs where peptide precursors were confidently detected and quantified—ranged from 77.3% to 83.1% per cell line, while the average completeness over all cell lines and replicates was 54.2%. Since CPTAC samples are profiled via a tandem mass tag (TMT)-based workflow, they present deeper coverage, with 31,339 phosphosites from 6,383 proteins, and a matrix completeness of 40.2%. However, they appeared to harbor considerable batch effects, due to the employed data-dependent acquisition (DDA) and TMT-labelling. To allow unbiased comparative analysis of the two datasets, we thus identified a shared subset comprising 8,617 phosphosites, with equivalent completeness (Methods). We then used mVESPA to measure protein activities using the dVESPA-inferred SigNet, as described above (Fig. 3, Methods), yielding an activity matrix for 381 KP-enzymes (Supplemental Fig. 5 – 7, Supplemental Table 3).

**Figure 3.**
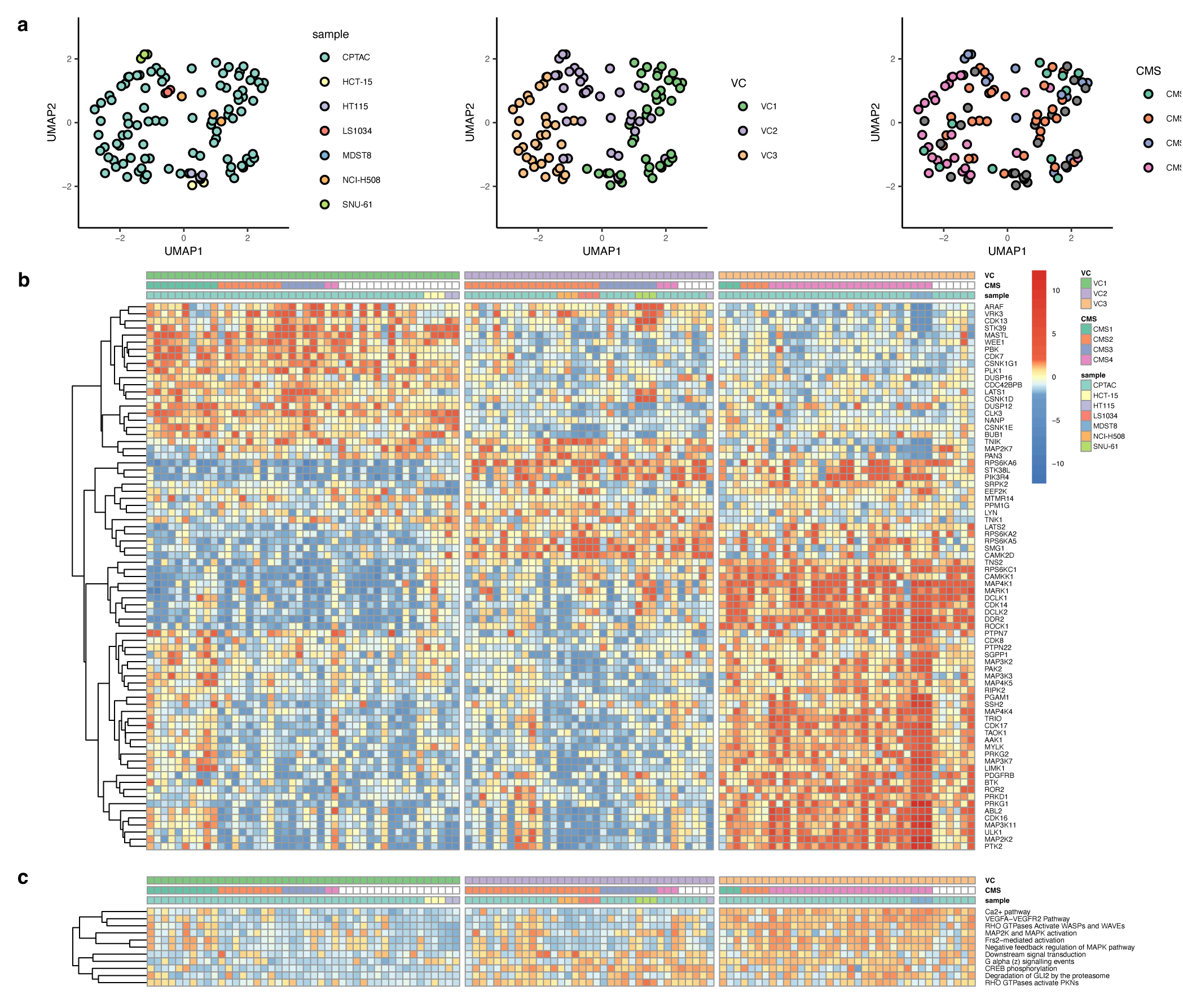
Representation of CRC subtypes by cell line models and activity-level VESPA. **a)** UMAP embedding of KP-enzyme activities and coloration according to different classification systems (phosphoproteome-based VESPA; VC and the CRC Consensus Moleular Signature; CMS). **b)** The most informative proteins and their VESPA inferred normalized enrichment scores (NES) have been selected for visualization (full datasets: Supplemental Fig. 5-7). CPTAC clinical profiles and cell lines were grouped according to the Consensus Molecular Classifier (CMS) and VESPA clusters (VC). The samples are grouped according to VC. **c)** Gene Set Enrichment Analysis (GSEA) using a signaling subset of the Reactome database. Only terms significant in at least one sample (BH-adj. *p*-value *<* 0.05) are shown. The colors represent GSEA NES and are linked to the legend in b).

K-medoids clustering [39] identified three main KP-enzyme activity-based clusters (VC_1_ – VC_3_) in the CPTAC dataset (Methods), while Random Forest recursive feature elimination identified the KP-enzymes with the greatest independent contribution to subtype classification (Fig. 3, Supplemental Table 4-5, Methods). KP-enzyme-based OncoMatch analysis confirmed that most of the cell lines recapitulated differential enzyme activity across the three subtypes. Specifically, HCT-15 and HT115 matched VC_1_, NCI-H508, LS1034 and SNU-61 matched VC_2_ and MDST8 matched VC_3_. Notably, one replicate of HT115 was assigned to VC_2_ instead of VC_1_.

Finally, we assessed the ability of the six cell lines to recapitulate the four subtypes (CMS_1_ – CMS_4_) identified by consensus transcriptomic cluster analysis in the Consensus Molecular Subtype (CMS) classification system [43], by the Colorectal Cancer Subtyping Consortium (CRCSC) (Methods).

Considering samples classified by both VESPA and CMS (CMS probability *>* 0.5), there was broad consistency between CMS and VESPA subtypes (Fig. 3a, colored, non-white labels). Specifically, VC_1_ was significantly enriched in CMS_1_, VC_2_ in CMS_2_ samples, and VC_3_ in CMS_4_ samples. Due to the smaller number of VESPA-subtypes, however, CMS_3_ samples were split between VC_1_ and VC_2_, likely reflecting finer-grain stratification at the transcriptional regulation level, likely due to epigenetics differences across tissues that would not directly affect signaling. Based on CMS classification, OncoMatch analysis identified the NCI-H508 and LS1034 cell lines as high-fidelity models for CMS_2_ samples, SNU-61 for CMS_3_, and MDST8 for CMS_4_, confirming that the selected cell line panel is broadly representative of patient-derived subtypes (Fig. 3a-b). HCT-15, HT115 could not be confidently classified by CMS to one of the subgroups. A recent study [44] observed similar results when matching cancer cell lines to CMS, obtaining good matches for MDST8, NCI-H508, LS1034 and SNU-61, and ambiguous matching for HT115 (HCT-15 was not reported).

Gene set enrichment analysis [33] (GSEA) using the Reactome database [45] supports these assignments since it identified several enriched signaling pathways in the three VESPA clusters (*p <* 0.05, Benjamini-Hochberg (BH)-corrected, see Methods) (Fig. 3c, Supplemental Table 6). For instance, we identified enrichment of VEGFA-VEGFR2 Pathway in VC_3_, a hallmark of the CMS_4_ subtype [43], which was further supported by the activation of RHO GTPases involved in WAVE complex regulation, a key regulator of actin-remodelling, invasiveness and EMT-like processes [46] (Fig. 3c). This was recapitulated by the MDST8 cell line in our panel, representing an established EMT model [47].

In summary, except CMS_1_, for which no representative cell lines could be identified, the six cell lines selected for our study effectively represent the major CRC subtypes inferred by either transcriptional or phosphoproteomic analysis.

### Drug Perturbation Profiles

To assess drug mechanism of action (MoA), CRC cell adaptive mechanisms leading to drug resistance, and potential treatment-mediated rewiring of signaling pathways, we designed a comprehensive longitudinal drug perturbation experiment (Methods). We focused specifically on seven compounds, based on their ability to target a diverse and complementary set of pathways relevant to CRC tumorigenesis. With the exception of WIKI4 (a TNKS & TNKS2 inhibitor), these represent FDA-approved drugs for the treatment of CRC and related cancer types, including alpelisib (PIK3CA), imatinib (ABL1/3 & c-Kit [48]), linsitinib (IGF1R [49]), osimertinib (EGFR-T790M), ralimetinib (p38 MAPK), and trametinib (MEK1 & MEK2). Although some of these drug compounds were originally found to target genes with specific or activating mutations (osimertinib [50] and alpelisib [51], respectively), we set up our experimental design and analysis strategy to disregard genetic dependencies since targeted drug compounds frequently also inhibit wild-type genes [51] or can have off-target effects on related proteins [52]. In the case of alpelisib, the cell line panel represents both mutated (HCT-15, HT115, NCI-H508) and wild-type (LS1034, MDST8, SNU-61) PIK3CA genes, whereas no cell lines of our panel (or any other CRC cell lines covered by CCLE) harbored a primary EGFR-T790M mutation targeted by osimertinib. Indeed, response to osimertinib was also reported for patients that lack the hallmark T790M mutation [53].

Assessing drug MoA requires careful selection of an optimal, physiologically achievable concentration *in vivo*, at which the MoA is manifested with the least contribution by confounding factors. For instance, selecting an exceedingly high concentration may induce confounding effects from both lower-affinity targets (off-target effects) and mechanisms associated with activation of cell stress and death pathways. Consistent with our prior studies [54, 55], we thus selected drug concentrations representing the lowest of the reported C_max_ (maximum tolerated serum concentration in vivo) and the 48h IC_20_ in the most sensitive cell line from our panel, as experimentally determined by dose response curves (Methods). Concentrations were also capped at ≤ 0.5*µ*M, consistent with maximum levels achievable in tissues. Based on this rationale, imatinib, osimertinib, ralimetinib and WIKI4 were titrated at 0.5*µ*M, while alpesilib, linsitinib and trametinib were titrated at 0.12*µ*M (IC_20_), 0.14*µ*M (IC_20_), and 0.036*µ*M (C_Max_), respectively (Methods). Differentiating between sensitive and resistant cell lines is non-trivial [56]. For example, the frequently applied threshold of 1.0*µ*M (IC_50_), would define 23 out of 27 of our investigated cell line and drug perturbation combinations with available GDSC reference data as non-sensitive or resistant (Supplemental Fig. 8a) [56]. For this reason, we transformed log(IC_50_) values to *z*-scores, computing relative metrics for all cell lines, aggregated per drug compound and GDSC dataset. To differentiate between more sensitive and more resistant combinations, we selected sensitive combinations as one datapoint reaching z-score *<*-1.0, with resistant combinations having z-score *>* 1.0 (Supplemental Fig. 8b). This identified trametinib-treated MDST8, LS1034, and NCI-H508, and linsitinib-treated LS1034, NCI-H508, as well as alpelisib-treated HCT-15 cells as most sensitive according to GDSC. Conversely, linsitinib-treated SNU-61, HCT-15, and HT115, as well as trametinib-treated NCI-H508 were identified as most resistant. Notably, trametinib-treated NCI-H508 was identified as both sensitive (GDSC2) and resistant (GDSC1) in different datasets.

We generated phosphoproteomic profiles from each cell line, by DIA-based proteomics (Methods), at seven time points following perturbation with each of the seven inhibitors and vehicle control (DMSO). This allowed assessing quantitative effects of KP-enzyme activity following short (5min, 15min), intermediate (1h, 6h) and long-term (24h, 48h, 96h) treatment. Cumulatively 336 phosphoproteomic profiles were acquired by label-free DIA, for quantification and statistical validation at peptidoform-level [25] (Methods). We refer to this dataset as “U54-DP”. To minimize cross-sample statistical dependencies that would affect the mutual information estimator in dVESPA for SigNet inference, we generated a reduced “U54-NET” dataset comprising only sufficiently spaced time points, i.e., 1h, 24h and 96h, respectively.

In total, 27,813 peptidoform precursors, 14,376 phosphosites and 3,786 phosphoproteins were identified and quantified at 1% global-context peptidoform and protein FDR [57] (Supplemental Data 2). Across all perturbations and time points, our workflow achieved high consistency on peptidoform-precursor level, on a cell line by cell line basis (48.7-55.6%), whereas the global completeness across all 336 runs of 36.6% indicates considerable biological inter-cell-line heterogeneity and different response to drug perturbations.

After data preprocessing—including normalization and missing value imputation (Methods)—we used VESPA to assess KP-enzyme differential activity in each cell line, at each time point following treatment with each drug *vs.* vehicle control, using the integrated substrate and activity level analysis. The resulting matrix represents the differential activity of 381 KP-enzymes across 336 sample conditions, with positive and negative NES values indicating either increased or decreased enzymatic activity (Fig. 4a, Supplemental Fig. 9-12, Supplemental Tables 7-9). Expectedly, this analysis showed that cell line identity dominated unsupervised hierarchical clustering when activity was computed only at the substrate-level (Supplemental Fig. 9-10, Supplemental Table 8), suggesting that drug response depends on cellular state. However, activity level-based analysis grouped cell line perturbations according to different signaling network states (Supplemental Fig. 11-12, Supplemental Table 9), as assessed by Reactome signaling pathways enrichment analysis (Fig. 4b, Supplemental Table 10).

**Figure 4.**
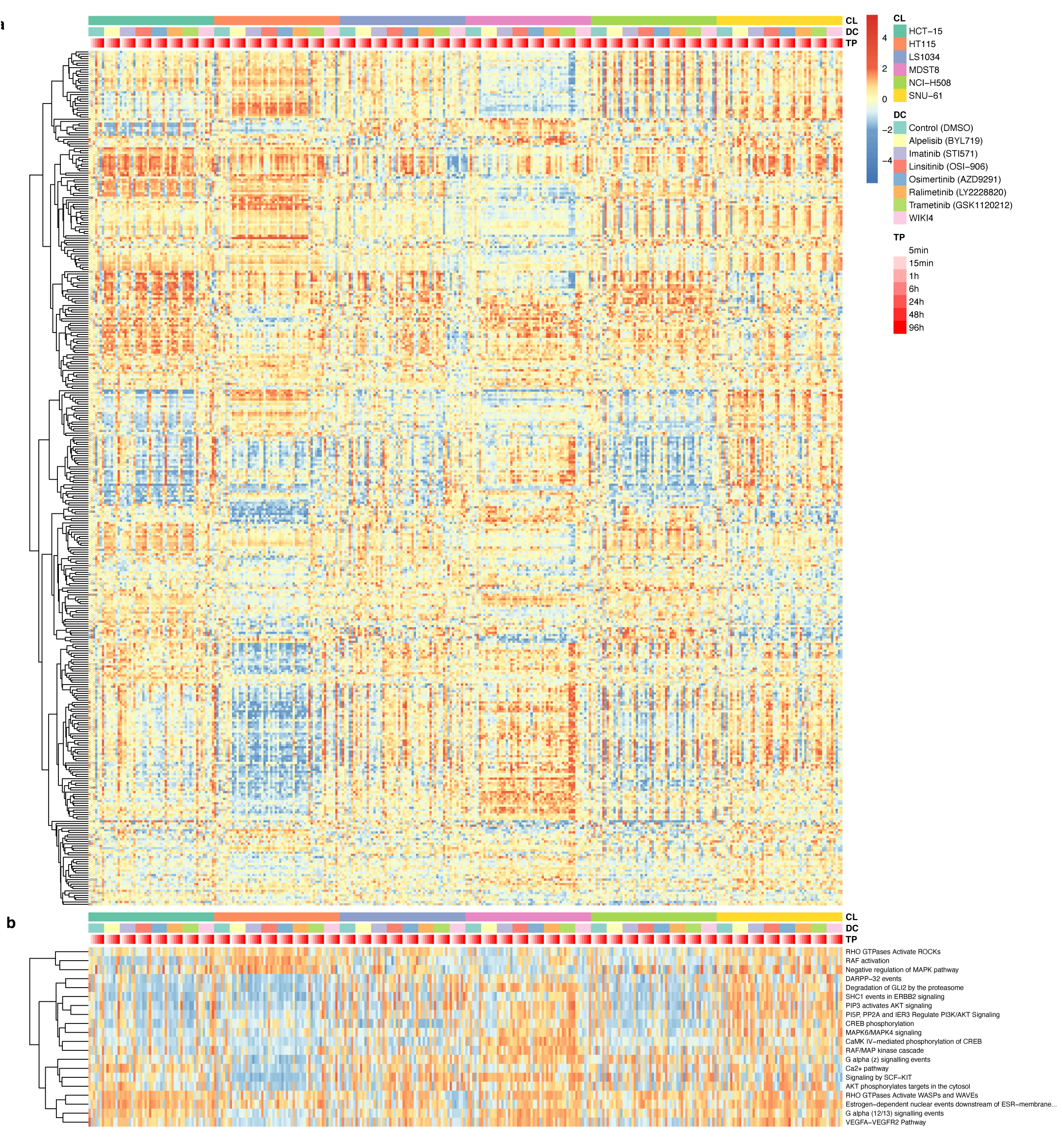
Targeted drug perturbations of CRC cell lines. **a)** A global overview of VESPA inferred normalized enrichment scores (NES) across the full drug perturbation dataset (336 samples), covering six CRC cell lines, 7 drug perturbations and DMSO control across 7 time points. **b)** Gene Set Enrichment Analysis (GSEA) using a signaling subset of the Reactome database. Only terms significant in at least one sample (BH-adj. *p*-value *<* 0.05) are shown. The colors represent GSEA NES and are linked to the legend in a).

As a first high-level validation, a focused analysis of primary (i.e., high affinity-binding) targets could be insightful, even though the applied drug concentrations should only result in perturbation and not full knock-down of their primary targets. We thus investigated the time-dependent effect of each drug on its established primary targets, as reported in DrugBank [58] and ProteomicsDB [59] (Fig. 5, Methods). For drugs with *>* 5 primary targets, we selected the five with highest average inhibition across all cell lines. This analysis shows that primary targets were effectively inhibited in responsive cell lines, yet with highly variable temporal effects, requiring from 5min to 96h before maximum inhibition was achieved. In addition, significant time-dependent variability was observed, likely due to cell homeostatic mechanisms driven by temporal responses to the low perturbational concentrations of the drugs.

**Figure 5.**
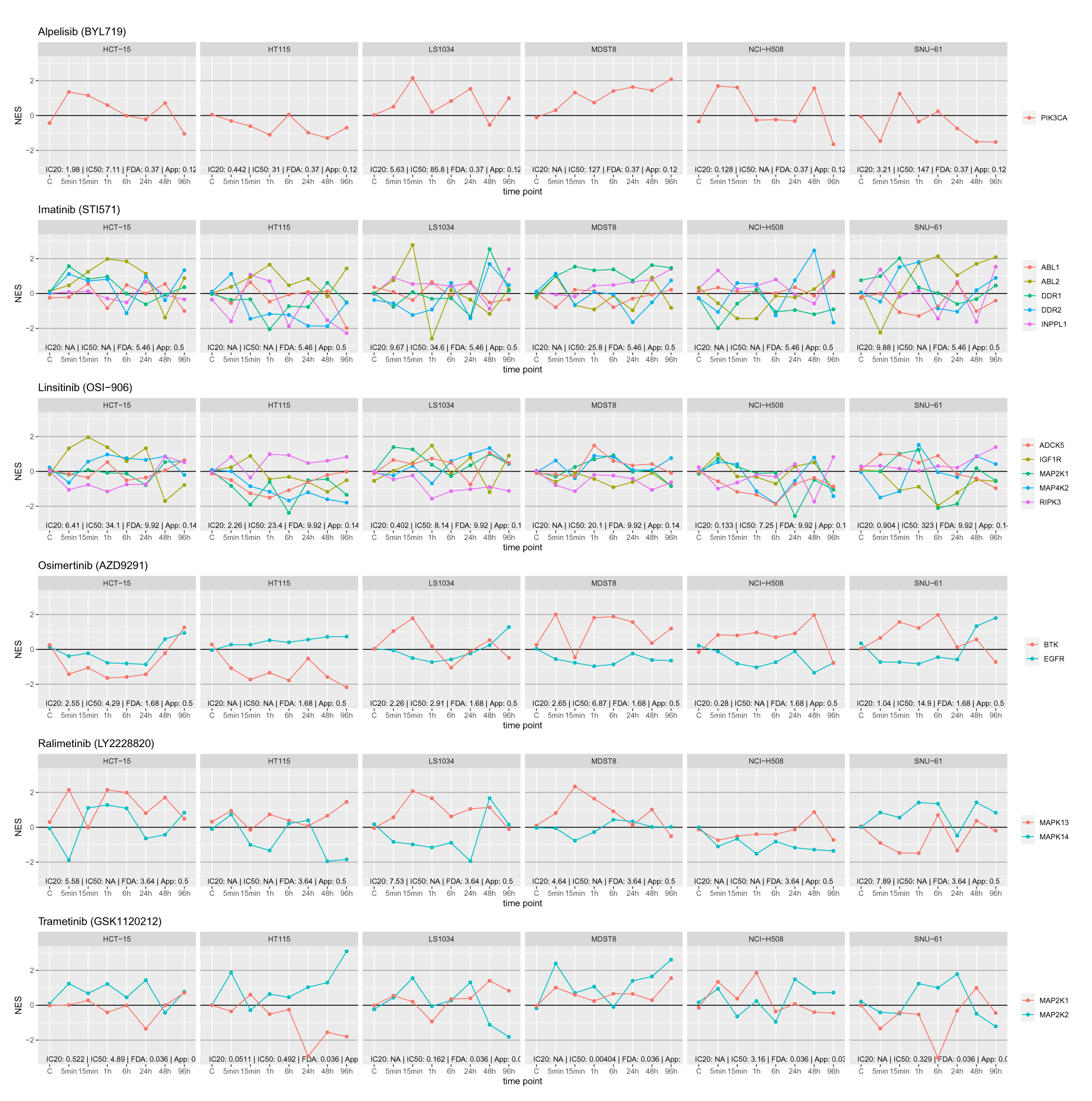
Temporal VESPA perturbation profiles of known primary drug compound targets. The VESPA normalized enrichment scores corresponding to Fig. 4 are extracted and visualized for the top 5 downregulated known primary targets, grouped according to drug perturbations and cell lines.

Further supporting the cell-line-specific effect of each drug, primary target inhibition across cell lines was highly variable even for the same drug. For instance, following ralimetinib treatment, MAPK13 (specifically targeted by ralimetinib) and MAPK14 activity was inversely correlated in LS1034, MDST8 and SNU-61 cells yet positively correlated in other cell lines (Fig. 5). Comparative analysis shows that raw phosphopeptide abundance of primary targets was often less informative than VESPA-measured KP-enzyme activity, frequently because active sites were not directly or only inconsistently measured (Supplemental Fig. 13). In addition, VESPA-based activity provided critical clues for the discovery of the enzymatically active sites using the signalons resolved to individual phosphosites. Since the profiled drug perturbations inhibited enzyme function in a targeted fashion, we found that the VESPA activities were frequently modulated for the active, but not all other phosphosites of the same kinase (Supplemental Fig. 14, Supplemental Table 11). For example, MAP2K2:S222 phosphorylation was previously associated with induced kinase activity [15]. Consistent with the literature, our data shows that trametinib-mediated MAP2K2 inhibition often results in consistently lower S222-specific, time-dependent, VESPA-inferred activity, whereas the time series profile of MAP2K2:S23 correlated only across some of the cell lines (Supplemental Fig. 14). Interestingly activity of MAP2K1:S298—a distinct, previously reported active site [15]—was anti-correlated with that of MAP2K2:S222, following trametinib treatment of HCT-15, HT115 and NCI-H508 cells, suggesting a compensatory mechanism. A similar pattern could also be observed for the correlation between MAPK14:Y182 activity and the activity of both MAPK13:S350 and MAPK13:T265, following ralimetinib treatment of HCT-15, HT115 and LS1034 cells (Supplemental Fig. 14). Additional established active sites targeted by specific drugs include EGFR:S991, EGFR:S1071 and EGFR:Y1092 (osimertinib), MAP2K1:S298 MAP2K2:S222, RIPK3:S227 (linsitinib) and INPPL1:S132 (imatinib) [15]. Taken together, these data show that VESPA analysis of data generated by drug perturbation assays can help elucidate subtype-specific drug MoA and cell adaptation mechanisms.

### Context-specific Signaling Network Adaptation and Rewiring

A primary goal of our experimental design was to study potential context-specific, drug-mediated signaling network buffering/rewiring to elucidate mechanisms of cell adaptive drug response. For this purpose, we analyzed KP-enzyme activity inferred by VESPA analysis, with the DeMAND algorithm [55] to identify sub-networks dysregulated by each drug (Methods). DeMAND assesses dysregulation of individual PPIs using the Kullback-Leibler divergence, by computing changes in mutual information across drug perturbations at different time points or concentrations *vs.* vehicle controls [55]. Enrichment of dysregulated PPIs (edges) emanating from each protein (node) in the network can then be used to identify those most affected by a drug.

The DeMAND analysis integrated two different network levels: First, dysregulation of the activity-level CRC-specific SigNet—comprising 14,390 high-confidence interactions between 329 proteins—was assessed based on time-dependent KP-enzyme activity (Methods). Additionally, we probed 915 high-likelihood (LR ≥ 0.5), non-phosphorylation-related interactions between 198 proteins from the STRING database [60] (Methods). Indeed, since phospho-state may affect protein conformation and thus the ability to form complexes, integration of protein-protein interactions is expected to further improve the DeMAND analysis [61, 62]. Results from the two analyses were then integrated (Methods, Supplemental Tables 12-13).

To assess both global (i.e., most conserved across all cell lines) and cell-line-specific drug MoA, two analyses were performed: For the former, we used data from all cell lines and time points *vs.* vehicle control samples. For the latter, the analysis was performed in cell line-specific fashion. In the global analysis, 62 significantly dysregulated proteins were identified (*p <* 0.05, BH-corrected) with an average of 12 to 21 proteins per drug. Hierarchical clustering of DeMAND-inferred MoA profiles identified cell lines presenting either congruent or divergent MoA for the same drug (Fig. 6a). Interestingly, some proteins were highly dysregulated by virtually all treatments, across most cell lines, including established colorectal cancer risk factors, such as PRKCZ [63], BMP2K [64] and MAPK14 [65]. This suggests that the signaling logic of the cell plays a critical role in canalizing the effect of drug perturbations into common dysregulation patterns.

**Figure 6.**
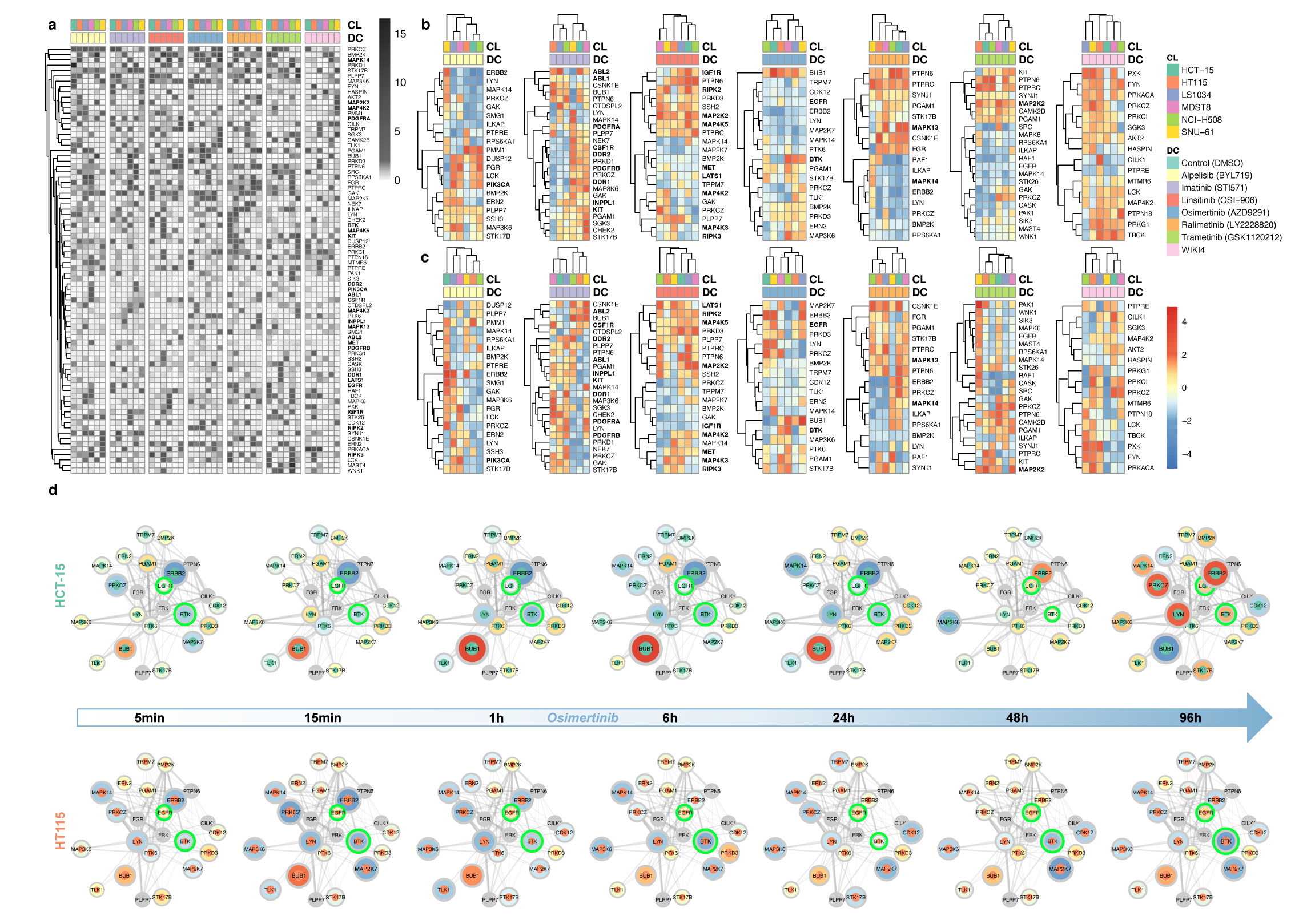
Context-specific wiring of signaling pathways. **a)** Analysis of the VESPA-inferred activities by the DeMAND algorithm identifies regulators with context-specific dysregulated interactions. The heatmap depicts significance of dysregulation (-log10(BH-adj. *p*-value). Only known drug targets (bold) and proteins with significant score (black: BH-adj. *p*-value *<* 0.05) in the unspecific DeMAND analysis are visualized. **b)** Grouping of the dysregulated proteins according to drug perturbations and overlay with VESPA-inferred activities of the aggregated early time points. **c)** Grouping of the dysregulated proteins according to drug perturbations and overlay with VESPA-inferred activities of the aggregated late time points. **d**) Visualization of network dysregulation and drug compound mechanism of action (MoA) for osimertinib. Nodes indicate the most affected regulators with the inner circos colors indicating cell line type and the outer circos color and node size indicating VESPA activity. The edges indicate dysregulated, undirected interactions between the regulators (Methods). Line thickness indicates significance of dysregulation. Proteins highlighted in green indicate known primary and secondary targets.

To assess the early *vs.* late effects of drug perturbations on these proteins, we plotted the VESPA-inferred activity of the proteins identified as most dysregulated by DeMAND for each drug, using either the early (5min, 15min, 1h) (Fig. 6b) or the late (24h, 48h, 96h) (Fig. 6c) time point measurements (Methods). As shown, for each drug, responses clustered into 1 to 3 sub-signatures (with most showing 2) indicating that drug response is mediated by distinct CRC-specific signaling networks. For instance, at the early time points, NCI-H508 and LS1034, both classified as high-fidelity CMS_2_ models, behaved similarly in 3 of the 7 treatments (imatinib, linsitinib, and ralimetinib) but not in the other 4.

As an illustrative example, two main clusters were identified in the early time points for osimertinib, including either NCI-H508, HCT-15, and HT115 (cluster 1) and MDST8, LS1034, and SNU-61 (cluster 2) (Fig. 6b). To illustrate how network wiring affects drug MoA, we thus visualized the propagation of signaling activity dysregulation over time on the most drug-dysregulated sub-networks of HCT-15 and HT115, as representative of the two clusters (Fig. 6d). While the activities of key dysregulated proteins—BUB1, ERBB2, LYN, PRKCZ—are very similar in the early (Fig. 6b), they are clearly different between HCT-15 and HT115 in the late time points (Fig. 6c).

Visualization of the signaling activity time course profiles for the two cell lines shows that activity of the primary drug target (EGFR) was not significantly affected, likely because it is not highly activated at baseline (Fig. 6a). However, for HT115, the known off-target [59] BTK was dysregulated, especially based on its interaction with ERBB2. ERBB2—a lower-affinity target of Osimertinib [52]—is deactivated at early time points for both cell lines, but activated at the 48h and 96h time points for HCT-15. The mitotic checkpoint serine/threonine kinase BUB1—which interacts with EGFR, BTK, ERBB2, LYN and PTK6—was strongly activated in HCT-15 up until the 24h time point, suggesting that tumor cell proliferation in this CRC cell line could be attributed, to some extent, to increased signaling activity of this protein [66]. Together with the late time point activation of LYN and PRKCZ (Fig. 6d), these alterations represent the main differences between the two cell lines. Interestingly, LYN has already been reported as a key mediator of resistance to EGFR inhibitors, due to its involvement in nuclear translocation of EGFR [67], while PRKCZ is mainly associated with the cancer cell response to nutrient deprivation in intestinal tumorigenesis [68], suggesting that HCT-15 underwent metabolic adaptations that mediate insensitivity mechanisms.

Similarly, at the early time points, ralimetinib also shows a comparable response across all cell lines, however at the later time points, a divergent response in two cell line clusters—*i.e.*, NCI-H508, HT115, LS1034 and SNU-61, HCT-15, MDST8, respectively, was found by the analysis. Across these clusters and two representative cell lines, HT115 and SNU-61, the primary ralimetinib targets (MAPK13 and MAPK14) show inverse temporal perturbation profiles (Supplemental Fig. 15). However, while MAPK14 inhibition in HT115 induced consistent inactivation of downstream MAPK targets, MAPK13 inhibition in SNU-61 resulted in alternating activation and inactivation of downstream targets, potentially due to a negative feedback loop.

In summary, DeMAND analysis shows that many of the dysregulated sub-network are subtype-specific and have distinct temporal patterns, which can also be explored in a visual graph representation (Supplemental Data 3). Moreover, the VESPA-measured time-dependent protein activity profiles, following drug perturbation, can be used to investigate differential mechanisms of action and cell adaptation associated with context-specific rewiring of signaling interactions. In addition, these results show that drug MoA is much more complex and cell context-specific than originally thought.

### Cell Adaptation-mediated Drug Resistance

Cancer resistance mechanisms are among the most critical issues preventing long-term efficacy of targeted drug compounds. While so far the focus has been on the discovery of genetic events leading to selection of drug resistant clones, elucidation of dynamic network-based adaptation without clonal selection is increasingly emerging as a promising avenue to improve therapeutic efficacy [69].

Using VESPA and our drug perturbation time series data, we investigated the adaptive response of kinases and phosphatases by comparing the late drug-perturbed time points (24h, 48h, 96h) against the corresponding DMSO controls. Specifically, our goal was to identify the KP-enzymes most likely to have induced the measured phospho-state signature of resistant cells, following drug treatment. For this purpose, we assessed the effect of drug perturbation *vs.* control in late time points for each cell line and drug perturbation separately using a time-point-paired *t*-test for differential testing of the VESPA inferred protein signaling activities (Methods, Supplemental Table 14). The *p*-values of all conditions were then integrated by Stouffer’s method to select significant, increased activity of candidate resistance factors across all conditions (*q*-value *<* 0.05, mean(*t*-statistic) *>* 0) (Fig. 7a, Methods, Supplemental Table 15).

**Figure 7.**
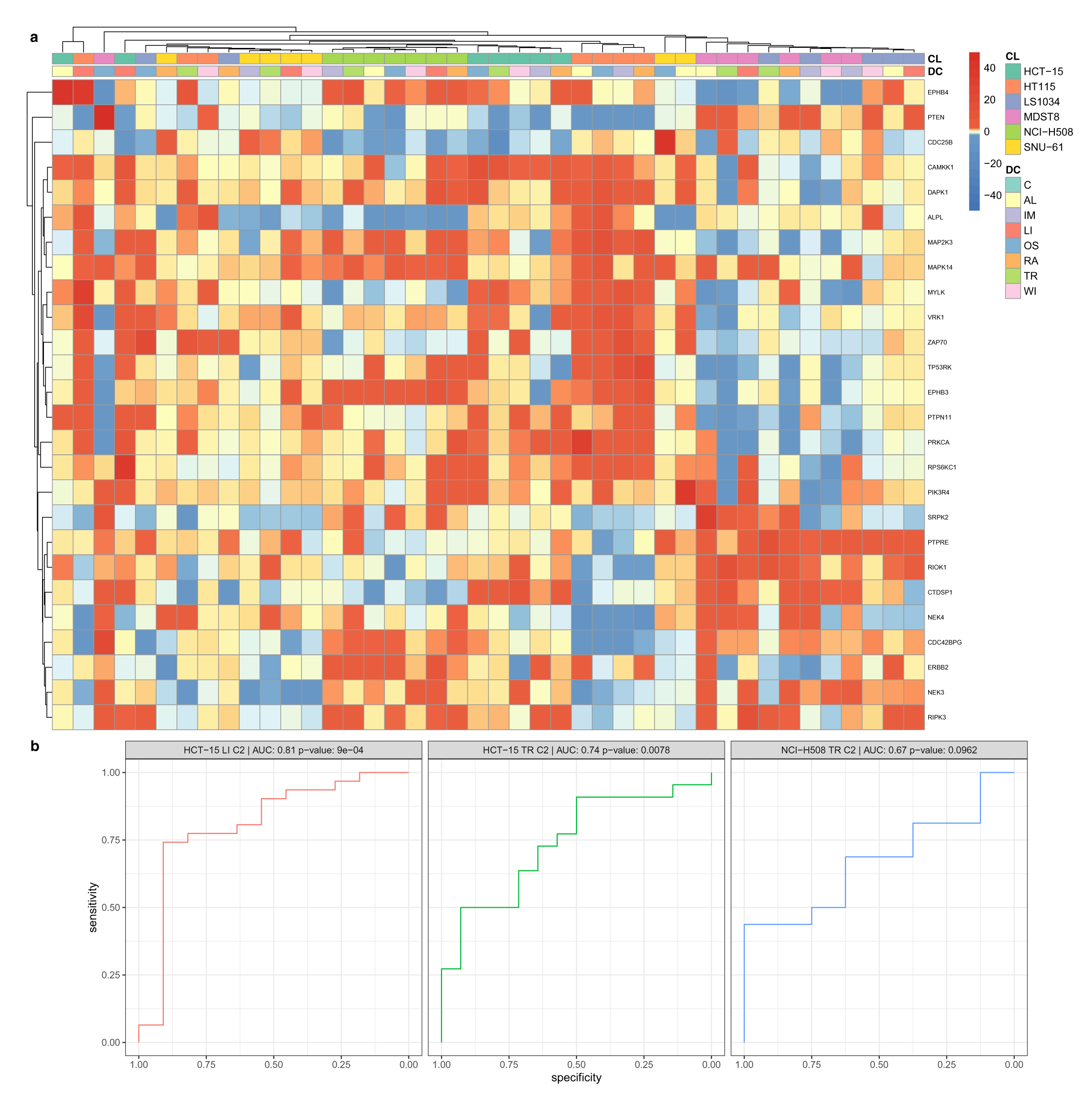
Context-specific adaptive stress resistance mechanisms. **a)** The effect of drug perturbation *vs.* control in late time points is visualized for each cell line and drug perturbation separately using the differential VESPA paired t-test *t*-statistic between late-time-point (24h, 48h, 96h) drug perturbation *vs.* DMSO control samples. The depicted KP-enzymes were selected by selecting those found to be significant (*q*-value *<* 0.05, avg(t-statistic) *>* 0) after integrating all *p*-values across all comparisons by Stouffer’s method (Methods). **b**) Receiver operating characteristics (ROC) with area under the curve (AUC) and statistical significance (Mann-Whitney-U test) are depicted for the CRISPRko validation experiment covering HCT-15 linsitinib (LI; C2: 4.0 *µ*M) *vs.* DMSO and trametinib (TR; C2: 0.7 *µ*M) *vs.* DMSO, as well as NCI-H508 trametinib (TR; C2: 0.01 *µ*M) *vs.* DMSO perturbation.

As in the network dysregulation analysis, the late-time-points-based clustering was dominated by cell line-specific effects. All drug perturbations for MDST8 and LS1034, except osimertinib and ralimetinib treated LS1034, resulted in increased signaling activity of a cluster of KP-enzymes including SRPK2 [70], PTPRE [71], RIOK1 [72], CTDSP1 [73], NEK4 [74], CDC42BPG [75], ERBB2 [76], NEK3 [77] and RIPK3 [78], all of which have been previously been associated with colorectal cancer tumorigenesis or drug resistance and insensitivity. The association of several of these KP-enzymes with the MAPK/ERK or STAT3 signaling pathways and their lower activity in osimertinib (targeting EGFR-T790M) and ralimetinib (targeting p38 MAPK) treated LS1034 cells, suggests that this pathway could be a key mediator of drug insensitivity in these two cell lines.

A similarly distinct cell-line-specific clustering profile could also be observed for HT115 cells perturbed with all drug compounds, except trametinib and WIKI4, resulting in increased signaling activities of KP-enzymes CAMKK1, DAPK1 [79, 80], ALPL [81], MAP2K3 [82], MAPK14 [82], MYLK, VRK1 [83], ZAP70 [84], TP53RK [85], EPHB3 [86], PTPN11 [87], PRKCA [88], and RPS6KC1, of which the majority has previously been associated with resistance or insensitivity mechanisms or tumor suppression in colorectal cancer. Although the tumor suppressor role of DAPK1, ALPL, EPHB3 and PTPN11 seems to conflict with VESPA’s association to drug insensitivity, these highly connected KP-enzymes frequently have very different, sometimes contradicting effects on tumorigenesis depending on cellular context. For example,

DAPK1, as an autophagy inducer, has a tumor suppressor effect at early stages of cancer progression, limiting genomic instability in response to metabolic and oxidative stress [79]. However, at later stages, autophagy contributes to the resistance of tumor cells to chemotherapy treatment by blocking apoptosis [79]. PTPN11, has similarly been associated with tumor suppression in genomic screens [89], however, mechanistic studies have also identified its potential as central target in intrinsic and acquired targeted drug resistance [87].

The other three cell lines HCT-15, NCI-H508 and SNU-61 also primarily exhibited cell-line-specific responses to drug perturbations, albeit with less distinct signatures. To validate the candidate resistance factors identified by VESPA and to systematically assess whether targeting of the predicted resistance factors would increase chemosensitivity of insensitive cells in a cell-line-specific matter, we conducted a large-scale, pooled CRISPR knock-out (CRISPRko) screen experiment, targeting all annotated human kinases, phosphatases and E3 ligases of our cell line and drug perturbation panel with four different guides per target (Methods, Supplemental Table 16). We selected resistant drug perturbation and cell line combinations according to the GDSC-based classification system described above (Supplemental Fig. 8b, Methods): For linsitinib, we selected HCT-15 (z-score = 1.12), but not SNU-61 (z-score = 1.55), due to its relative complex culturing conditions. For trametinib, we selected HCT-15 (z-score = 0.89) and NCI-H508 (z-score = 1.13), even though the combination of NCI-H508 and trametinib resulted in discrepant sensitive (GDSC2) and resistant (GDSC1) responses within the two datasets and HCT-15 did not reach the strict threshold for our classification as resistant.

To validate these predictions, we performed CRISPRko screens in HCT-15 cells treated with linsitinib for 10 population doublings (C1: 1.0 *µ*M, C2: 4.0 *µ*M) and trametinib (C1: 0.1 *µ*M, C2: 0.7 *µ*M), as well as in trametinib treated NCI-H508 cells (C1: 0.005 *µ*M, C2: 0.01 *µ*M). DMSO was used as vehicle control. The initial (drug / DMSO-free) time point-samples (T0) for these screens were collected approximately 5-7 days after the sgRNA lentiviral transductions and puromycin selection. To pick the correct drug concentrations for the pooled CRISPRko screens, we performed a long term (10 population doublings) growth test for each cell line and their corresponding drug(s) with multiple different drug concentrations (Methods). For the CRISPRko screens, we picked two drug concentrations for each cell line, which appeared to only have a perturbation, but not a full inhibition effect, analogously to the phosphoproteomic perturbations (Methods). The only exception was the cell line NCI-H508, where we had to use a lower drug concentration for the long term pooled CRISPRko screening, due to drug toxicity manifesting after 96h time point (last time point of the short-term assay). Differential sgRNA abundance analysis was performed using DESeq2 (Methods, Supplemental Table 17). Sequencing quality was found to be excellent, with an average alignment ratio of 90.98% (Supplemental Fig. 16).

Differential expression analysis of DMSO *vs*. T0 samples identified known essential genes for CRC with an area-under-the-curve (AUC) of 0.96 for both NCI-H508 and HCT-15 (Supplemental Fig. 17, Methods).

For tumor suppressors that can also act as resistance or insensitivity factors, such as DAPK1 or PTPN11, the nature of perturbation or knock-out will substantially bias their activity and function [90]. It was recently suggested that tumor suppressor genes, or genes whose knock-out imparts a growth advantage on cells, could cause recurrent drug suppressor hits in drug-gene interaction CRISPRko screens, and thus a source of a systematic bias and false positives in drug-perturbed CRISPRko screens [90]. There is thus a potential discrepancy in the experimental design of the VESPA predictions and the CRSIPRko experiment, where VESPA predicts KP-enzyme late-timepoint activity and potential involvement in resistance or insensitivity mechanisms, whereas the CRISPRko experiment assesses their gene essentiality starting from timepoint 0 in combination with drug perturbations for altogether 10 population doublings. For this reason, we excluded knock-outs of known tumor suppressors [91] from the analysis (Supplemental Fig. 18-19, Methods).

To compare candidate resistance factors predicted by VESPA with the ground truth from CRISPRko assays, we conducted separate analyses for each cell line and drug perturbation to assess receiver operating characteristics (ROC) (Methods). Gene essentiality (log-fold-change perturbation *vs.* control, including negative (*i.e.*, essential) and positive (*i.e.*, non-essential) values), is expected to be inversely correlated to VESPA-assessed activity (*t*-statistic perturbation *vs.* control; positive: increased activity, negative: decreased activity).

The analysis strongly supports the relevance of VESPA’s predicted resistance factors in combination with the drug perturbations (Fig. 7b). ROC was found to be particularly significant for HCT-15 perturbed by linsitinib and trametinib (AUC = 0.81, *p*-value = 9e-04; AUC = 0.74, *p*-value = 0.0078, respectively), with only slightly lower performance for NCI-H508 perturbed by trametinib (AUC = 0.67; *p*-value = 0.0962). Correlation analysis further shows that VESPA can identify high numbers of true positive candidates with only few false positives (Supplemental Fig. 19), an essential requirement for diverse applications.

In summary, VESPA in combination with the time-series drug perturbation phosphoproteomic profiles allows to identify candidate resistance factors that could be exploited for combination therapies. The CRISPRko validation experiment demonstrated VESPA’s excellent predictive performance to identify candidates for linsitinib and trametinib treated cells.

## Discussion

The central importance of kinases and phosphatases as key regulators of signaling pathways has driven the development of targeted drug compounds, particularly to treat diseases where sustained proliferative signaling is a hallmark [4]. Although the phosphoproteome has been identified and quantified by mass spectrometry-based bottom proteomics since decades [92], enormous efforts are still required to systematically characterize diseases with comprehensive coverage of the phosphoproteome while simultaneously representing the diversity of observed clinical subtypes. For different cancer types, including CRC [36] and more than ten others, CPTAC initiatives have achieved or are working towards comprehensive proteogenomic characterization of dozens to hundreds of tumors, representing the disease landscape [22]. These datasets, whose acquisition frequently require the collaborative efforts of several research groups, provide invaluable resources for integrative or follow-up studies.

In this study, our goal was to identify context-specific wiring of signaling pathways and adaptive stress resistance mechanisms of CRC subtypes in response to targeted drug perturbations. For this purpose, we designed a comprehensive experiment, measuring the phosphoproteomic profiles of six representative CRC cell lines, perturbed by seven targeted drug compounds and controls across seven time points. Including the baseline profiles, we acquired 354 phosphoproteomic profiles, which, to our knowledge, resulted in one of the largest context-specific targeted drug perturbation phosphoproteomic datasets. The measurement of this large sample cohort required a flexible and scalable approach. The recent development of new data-independent acquisition (DIA) strategies [23, 24] and corresponding computational analysis Methods [25, 93, 94], provided an opportunity for the comprehensive and consistent quantification of the phosphoproteomic profiles within less than 3 weeks of instrument time for the full dataset. Although our unfractionated, label-free approach provided substantially lower coverage of the phosphoproteomes than the fractionated, label-based CPTAC studies, we speculated that quantitative consistency within the sample cohort might be more important than the depth of proteome coverage for our research questions. Further, we hoped that an ideal analysis strategy could allow to transfer knowledge and hypotheses from the comprehensive CPTAC to our focused drug perturbation profiles. For this purpose, we developed VESPA.

Identically to previous approaches [18, 19, 61, 62, 95, 96], VESPA is based on the concept that the activity of kinases and phosphatases can be inferred from their substrate abundances. However, instead of relying on generalized KP→S interaction databases, a key feature of VESPA is the generation of context-specific signaling networks from comprehensive phosphoproteomic profiles, such as the CPTAC studies. This has several advantages, but most importantly, the extensive space of chemically possible KP→S interactions is reduced to those are active within a specific disease context, dramatically improving the specificity of the inferred protein signaling activities. Further, in comparison to pathway databases, our approach provides many more KP→S interactions per regulator (Pathway Commons: ^∼^70; VESPA: ^∼^500, with mode of regulation and probabilistic weight) and covers more kinases and phosphatases (Pathway Commons: 211; VESPA: 371) when considering the coverage of the measured substrate proteins. In addition to the improved sensitivity of VESPA, the inferred protein activities can optionally be assessed in a site-specific manner and corrected for signaling cross-talk. Cross-talk represents a critical property of cellular signaling, which can only be addressed with the context-specific, comprehensive signaling networks generated by VESPA and its analytical framework based on the original VIPER algorithm [29]. Finally, VESPA introduces a hierarchical approach to activity inference: Although affinity chromatography enrichment methods are robust and generic, they introduce different biases within the measured phosphoproteomes. First, the current methods are limited in substantially enriching phosphorylated tyrosine residues of the frequently disease-relevant tyrosine kinases. Second, the coverage of enriched serine and threonine phosphopeptides varies substantially between different subtypes or cell lines, particularly with shallow profiling methods, thus limiting inter-cell-line comparability. To some degree, the hierarchical approach employed by VESPA can account for these issues by further abstraction of the signaling network and inference of tyrosine kinase activity based on substrate activity instead of substrate phosphosites directly. In a future extension of VESPA, less scalable TK-enriched phosphoproteomic profiles measured for only few conditions might help to more accurately estimate activity-level kinase signaling activities based on more specific TK-signalons that can also be applied to affinity-chromatography-enriched datasets. Using extensive benchmarks, we validated the improvements of each component of VESPA and demonstrated substantial improvements over the state-of-the-art. However, due to the intrinsic limitations of the benchmark, specifically the bias towards well studied kinases, and the requirement that primary target activities need to correspond with drug sensitivities of the specific cell lines, we believe that our benchmark represents a lower estimate of the potential improvements of VESPA.

We supported the use of our CRC cell line panel by comparing the transcriptional and signaling network states with the CPTAC tumor profiles (Fig. 3a-b). Both within the established transcriptional Consensus Molecular Classifier (CMS) of the Colorectal Cancer Subtyping Consortium (CRCSC), and the VESPA clusters, the cell lines represent the tumor samples.

The assessment of our drug perturbation study from three different perspectives, 1) temporal activity profiles of known primary drug targets, 2) the systematic assessment of context-specific wiring of signaling pathways, and 3) the identification context-specific adaptive stress resistance or insensitivity mechanisms, illustrates the diverse possibilities for data interpretation.

The temporal activity profiles showed that phosphosites of primary targets very often are difficult to measure consistently or do not show a direct response, especially in contrast to VESPA-inferred signaling activities, which in many cases could even resolve response profiles to the level of individual phosphosite activities (Supplemental Fig. 14). We believe VESPA’s unique feature to generate site-specific, data-driven signaling networks will be useful to support more mechanistic investigations of signaling networks in future studies.

Drug perturbation-dependent rewiring of signaling networks is a critical component of adaptive response. Our network dysregulation analysis based on DeMAND demonstrated the value of context-specific interactions for this purpose. In contrast to the original implementation for transcriptomic data [55], we believe that our application to phosphoproteomic data could provide a more direct measure of protein activity, and thus more mechanistic insights, as demonstrated by the propagation of adaptive response through neighboring KP-enzymes (Fig. 6d, Supplemental Fig. 15).

Differential analysis of late *vs.* early time point VESPA activities predicted a set of candidate vulnerabilities for each drug perturbation and cell line combination (Fig. 7a). While a considerable fraction of those has been validated as resistance factors in CRC, and some targets even being assessed as complementary target to reduce resistance or insensitivity, we wanted to experimentally validate the most resistant combinations.

The CRISPRko experiment targeting all kinases, phosphatases and E3 ligases provided the ideal high-throughput, orthogonal dataset for this purpose. Although both VESPA and the CRISPRko experiment have intrinsic biases and limitations, it allowed us to confidently support VESPA’s predictions with high selectivity and sensitivity and to position VESPA as a useful tool to predict potential targets for combination therapies. In future applications, the cell line subtypes could potentially be mapped back to clinical subtypes for follow-up translational studies, either on signaling network or transcriptional level, to assess the suitability of combination therapies for personalized medicine.

Although our study focuses on the phosphoproteomic profiles, the signaling activities inferred by VESPA are ideally suited and directly compatible with upcoming methods for causal integration of multi-omic profiles, e.g. via TieDIE [61] or COSMOS [62].

VESPA is directly compatible with popular upstream bottom-up proteomic workflows and can be easily adapted for various experimental designs. The algorithmic components are available as platform-independent open-source software under a non-commercial usage license. We hope that VESPA can become a useful tool to assess upcoming medium to large-scale model system and clinical phosphoproteomic profiles in a context-specific fashion.

## Acknowledgments

This study was supported by a supplemental grant to NCI U54 CA209997 (Cancer Systems Biology Consortium). G.R. was supported by grants P2EZP3 175127 and P400PB 183933 from the Swiss National Science Foundation. Y.L. was supported by the National Institute of General Medical Sciences (NIGMS) through grant R01GM137031. A.C. was supported by NCI U54 CA209997 (Cancer Systems Biology Consortium). G.R.’s computational work was supported by NIH Shared Instrument grants S10 OD012351 and S10OD021764.

## Author contributions

- G.R.: Conceptualization, Methodology (VESPA, Benchmarking, Analysis, Integration), Software (VESPA, Benchmarking, Analysis, Integration), Validation, Writing – Original Draft, Visualization, Funding Acquisition
- W.L.: Methodology (Phosphoproteomics), Writing – Review & Editing
- M.T.: Methodology (CRISPRko), Validation (CRISPRko), Writing – Review & Editing
- J.H.: Methodology (hpMI, stDPI), Software (vespa.aracne), Writing – Review & Editing
- P.S.S.: Methodology (Cell culture, Drug sensitivity assays, Drug perturbation assays), Writing – Review & Editing
- S.P.: Methodology (Cell culture, Drug sensitivity assays, Drug perturbation assays), Writing – Review & Editing
- A.T.G.: Methodology (CRISPRko), Software (DESeq2-based analysis), Writing – Review & Editing
- C.K.: Methodology (Cell culture, Drug sensitivity assays, Drug perturbation assays), Writing – Review & Editing
- P.K.: Methodology (CRISPRko), Validation (CRISPRko), Writing – Review & Editing
- D.M.: Conceptualization, Writing – Review & Editing, Project Administration, Funding Acquisition
- B.H.: Conceptualization, Writing – Review & Editing, Funding Acquisition
- Y.L.: Conceptualization, Methodology (Phosphoproteomics), Writing – Review & Editing, Supervision, Funding Acquisition
- A.C.: Conceptualization, Writing – Original Draft, Supervision, Funding Acquisition

## Declaration of Interests

A.C. is founder, equity holder, and consultant of DarwinHealth Inc, a company that has licensed some of the algorithms used in this manuscript from Columbia University. Columbia University is also an equity holder in DarwinHealth Inc and assignee of patent US10,790,040, which covers some components of the algorithms used in this manuscript. The other authors declare no competing interests.

## Methods

### VESPA

#### Data preprocessing

The primary input data for VESPA are quantitative, proteotypic/unique peptide-level phosphoproteomic profiles from bottom-up mass spectrometry experiments. The data format is a long list (hereafter referred to as VESPA input list (PVL)) consisting of the columns “gene id” (UniProtKB entry name without species, e.g. “EGFR”), “protein id” (UniProtKB entry identifier, e.g. “P00533”), “peptide id” (free text unique peptide identifier from upstream software), “site id” (unambiguous combination of gene id, protein id and phosphosite, separated by “:”, e.g. “EGFR:P00533:S229”), “modified peptide sequence” (free text modified peptide sequence from upstream software), “peptide sequence” (free text unmodified peptide sequence from upstream software), “phosphosite” (unambiguous phosphosite identifier, e.g. “S229”), “run id” (free text sample or MS run identifier), “peptide intensity” (float log2-transformed peptide intensity from upstream software). To avoid any ambiguities and to allow for data transferability, all peptide sequences, phosphosites and protein names and identifiers are expected to be mapped to UniProtKB. If a phosphosite is covered by multiple peptide precursors, the most consistently detected peptide precursor is used to represent the phosphosite. If a peptide precursor contains multiple phosphorylated sites, redundant entries for each phosphosite are added. Each dataset (e.g. CPTAC sample cohort or study) should be stored in a separate PVL to ensure that differences in experimental design or batch effects can be accounted for in downstream steps.

Optionally, protein-level abundances can be used for the inference of signaling networks (see below). The file format is a similar PVL, however columns “peptide id” (free text unique protein identifier), “site id” (unambiguous combination of gene id, protein id and ”PA” (protein abundance), separated by “:”, e.g. “EGFR:P00533:PA”), “modified peptide sequence” (free text unique protein identifier), “peptide sequence” (free text unique protein identifier), “phosphosite” (”PA” (protein abundance)), and “peptide intensity” (float log2-transformed protein intensity from upstream software) are slightly altered.

The “vespa” R-package provides fully automated import functionality for the OpenSWATH [26], IonQuant [97], MaxQuant [98] and the CPTAC [99] file formats. Support for other file formats can be easily enabled by adapting the reference implementation. During data import, peptide sequences are first mapped to a user provided UniProtKB/SwissProt FASTA database to ensure consistent mapping of phosphosites and identifiers. Peptide intensities are (optionally batch-wise) quantile normalized and centered.

#### Inference of signaling networks

Bottom-up proteomic experiments spanning dozen to hundreds of samples are affected by both biological and technical variability. For phosphoproteomic experiments, accounting for the technical effects can be particularly challenging, because different sample preparation workflows, phosphopeptide enrichment strategies, labelled or label-free quantification, biochemical peptide fractionation, data acquisition techniques and signal processing can have a dramatic effect on phosphoproteome coverage, depth and consistency, introducing missing values that can originate from various causes. To ensure the integrity of the algorithmic assumptions, for each dataset or PVL, a separate signaling network is generated by a fully integrated Snakemake [100] workflow (“vespa.net”) consisting of the “vespa” and “vespa.db” R-packages and the “vespa.aracne” algorithm:

##### Data preprocessing

The PVL is transformed to a peptide-level quantitative matrix with missing values designated as NA. The matrix is peptide-wise rank-transformed over all samples with missing values being retained. To restrict the query space of regulator-target interactions, several options are available: a) all combinations of peptides (not recommended), b) non-directional regulation, where a regulator list is provided to restrict the query space, c) directional regulation, where activating and repressing regulator lists are provided to restrict the query space based on positive (kinases) or negative (phosphatases) correlation with substrates, and d) reference network, where a list of regulators and targets with optional priors is supplied. For options b) and c), a list of targets can be supplied if not all peptides of the matrix should be queried as substrates.

##### Hybrid adaptive partitioning to estimate mutual information

Peptides measured by bottom-up proteomics have individual limits of detection (LOD) and limits of quantification (LOQ), resulting in censored values in most proteome-wide studies. Over the full distribution of peptide intensities of a proteome, the absence of peptides in some samples not due to technical effects (e.g. stochastic data-dependent acquisition or batch effects) might thus contain information about those peptides not reaching LOD/LOQ abundance levels. To make use of this information and to estimate mutual information (MI) between the two sparse abundance rank vectors of regulators (R) and targets (T), a novel hybrid adaptive partitioning algorithm implemented in “vespa.aracne” is used: The two vectors are split in four quadrants: 1) data points without missing values in R and T, 2) data points with missing values in both R and T, 3) data points with missing values in R, and 4) data points with missing values in T. For quadrant 1, the MI is estimated by the algorithm originally implemented in ARACNe-AP [27]. For quadrants 2-4, the MI is estimated by assessing the numbers of data points in each quadrant in comparison to the other quadrants. The MI of all quadrants are then combined and normalized, providing a more robust metric to assess the relationship between regulators and targets.

##### Estimation of mutual information threshold

To estimate a lower MI threshold for random interactions, the rank-transformed quantitative peptide matrix including missing values is permutated while maintaining the number of missing values. Based on the query space restrictions, the MI distribution of all interactions is computed, a null distribution is fitted and the MI threshold fulfilling a user-defined family-wise error rate (default: FWER=0.05) is estimated.

##### Bootstrapped network reconstruction

Using the estimated mutual information threshold, and the hybrid adaptive partitioning mutual information algorithm, the bootstrapping and network reconstruction algorithm from ARACNe-AP [27] is applied, randomly sampling N (default: N=200) samples of the quantitative matrix, inferring mutual information and removing interactions not fulfilling the MI threshold. To remove putative indirect interactions, the Data Processing Inequality (DPI) can be applied, which resolves regulator-regulator-substrate dependencies [28]. If signal transduction regulation query restrictions were imposed, the novel stDPI function of “vespa.aracne” is used, which ensures that only valid regulator-regulator-substrate triangles are considered, either kinase-kinase-substrate (i) or phosphatase-kinase-substrate (iv) relationships, but not phosphatase-phosphatase-substrate (ii) or kinase-phosphatase-substrate (iii) relationships to ensure valid application of the DPI (Supplemental Fig. 1b).

#### Consensus network generation

In the final step, the consensus network generation approach of ARACNe-AP is used, where the statistical significance of edge detection is estimated based on a Poisson distribution and the network is filtered to significant interactions only (default: BH-adjusted *p*-value*<*0.05). At this step, two networks are generated, one on phosphosite-and the other on whole protein level, where all phosphosites are combined.

#### Signalon generation

Based on the consensus network, signalons, i.e. the set of substrate peptides regulated by the same kinase or phosphatase, are generated for use with the activity inference module of VESPA by adapting the approach originally developed for VIPER [29]: Peptide identifiers are mapped back to site identifiers to ensure transferability between different datasets. For each interaction, the probabilistic weight is computed by normalizing the interaction MI by the maximum MI across the network. For each interaction, the optional prior from the reference networks is normalized by the maximum prior specific to each regulator. The mode of regulation is then determined as described previously [29] by fitting a three-Gaussian mixture model, representing clearly repressed, clearly activated and non-monotonically regulated targets. However, Spearman’s correlation coefficient is computed by using the pairwise complete datapoints only. Finally, the signalons are restricted to the adaptive top N (default: *N* = 500) substrates, selected by decreasing probabilistic weight until threshold T 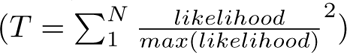 is reached, optionally weighted by the reference network priors. Signalons need to consist of at least M (default: *M* = 5) substrates to be considered.

#### Activity-level network reconstruction and signalon generation

The exact same steps as described above are applied to generate activity-level networks and signalons, however, instead of using a peptide-level PVL as input, a substrate-level PVL aggregated to the protein identifiers is used as described below. Further, non-signal transduction and standard DPI will be used in ”vespa.aracne”, allowing to abstract the signaling network to a functional instead enzymatic representation of the system.

#### Protein abundance-level regulation of substrates

In some cases, the signaling activity of kinases and phosphatases does not only correlate with their phosphorylation state, but also with their protein abundance, e.g. EGFR in cancer cells [101]. To assess tyrosine kinases, where the active sites involve phosphorylated tyrosine residues that are difficult to measure by affinity chromatography enrichment methods, using protein abundance as proxy for activity might be useful. For network reconstruction, a run identifier matched PVL of protein abundances, as described above, can be supplied that is used to assess the interactions of regulators represented by their protein abundances with downstream substrates.

### Inference of kinase/phosphatase activities

#### Inference of signaling activity

To infer kinase and phosphatase activities on substrate-level, the signalons from the steps above as well as the PVL used for their generation, or from an independent but biologically related experiment, represent the main input to the “viper” [29] R-package. First, the PVL is transformed to a quantitative matrix, with missing values imputed by the row-wise minimum, adding numerical jitter to break ties. The parameters for the “viper” activity inference function can be tailored for different applications and support the same experimental designs as the original implementation. Most importantly, a bootstrapped null model can be employed using the “viperSignature” function to assess differential protein signaling activity in comparison to a reference dataset. To infer activity-level kinase and phosphatase activity, a substrate-level activity matrix is first generated as described above and used as input for the “viper” function in combination with activity-level signalons. By default, signalons need to consist of at least M (default: *M* = 10) substrates to be considered. Substrate-and activity-level VESPA activities are then integrated using Stouffer’s method.

#### Cross-talk correction

Within the CRC-specific signaling network presented here, VESPA signalons have a median number of 547 or 43 substrates on substrate-and activity-levels, respectively. This results in considerable cross-talk between signalons and signaling pathways that should ideally be corrected for. For this purpose, VESPA is fully compatible with the shadow analysis [34] and pleiotropy correction [29] methods implemented in the “viper” function [29]: All signalon pairs affected by cross-talk are generated that fulfill two conditions: First, signalons A and B need to be both significantly enriched (default: *p*-value *<* 0.05) in the peptide abundance signature.

Second, they need to coregulate at least five substrates. To assess whether the enrichment of either signalon is driven by the coregulated substrates, the enrichment of the coregulated substrates within two subsets consisting of the signalon A and B substrates is computed, resulting in the estimated enrichment *p*-values *pA* and *pB*. Identically to “viper” [29], the cross-talk differential score is computed as *CDE* = log_10_(*pB*) − log_10_(*pA*). If *pA < pB*, the coregulated peptides of signalon A are penalized by *CDE^CI/NT^*, where the cross-talk index (CI) is a constant (default: *CI* = 20) and NT is the number of test pairs where signalon A is involved and vice versa.

### Signaling network optimization

VESPA signalons are typically generated for multiple dependent or independent phosphosites, one or several independent datasets and are potentially generated by using different priors from reference databases or predictive algorithms. To select the best signalon for each phosphosite and/or protein, the metaVIPER [35] approach is employed by VESPA. Briefly, among the total set of available signalons for a specific phosphosite or protein, the one resulting in the highest NES according to the rank-normalized [39] signature is selected. In absence of suitable context-specific phosphoproteomic datasets, this strategy can also be used to generate an optimized set of signalons for any target phosphoproteomic profiles.

Because phosphopeptides frequently carry several phosphosites, site-specific regulons are redundant. To also provide a non-redundant set, VESPA identifies and removes orthogonal regulon using the “findCorrelation” function from the R-package “caret” with a specified correlation cutoff (default: *C* = 0.5).

### Integrated generation of signalons on substrate-and activity-level

VESPA integrates all steps described above to generate and optimize substrate-and activity-level signalons by the “vespa.net” Snakemake workflow in a fully automated fashion. As primary input, one or several context-specific PVL of phosphoproteomic (and optionally related protein abundance) profiles are supplied. In addition, a PVL representing the reference phosphoproteomic dataset for which the signalons should be optimized are required. “vespa.net” then by default generates directional regulation / stDPI substrate-level signalons, as well as separate signalons constrained by and initialized using the priors from Pathway Commons, HSM/P and LinkPhinder. For each phosphosite and protein, optimized “meta” signalons are then generated by the optimization function described above and the reference phosphoproteomic dataset.

This network of optimized protein/substrate-level signalons is then used to infer cross-talk-corrected substrate-level protein signaling activities for each separate PVL. This new quantitative matrix in turn is then used to repeat the full network reconstruction process and to infer activity-level signalons. However, instead of directional regulation, undirectional regulation / DPI is used.

### Application to target datasets

After running “vespa.net” and generating substrate-and activity-level signalons, “vespa” and the “viper” function is applied to the phosphoproteomic profiles used previously as reference network. At this step, the experimental design, e.g. control runs within a drug perturbation experiment, can be used as null model as described above. The VIPER and VESPA frameworks provide a flexible toolkit suitable for several applications. The tutorial dataset (“vespa.tutorial”) illustrates the use cases of this study and describes the required parameters.

## Cell culture

The six CRC cell lines used in this study (HCT-15, HT115, LS1034, MDST8, NCI-H508, SNU-61) were previously selected to ideally represent the clinical phenotypes covered by TCGA as assessed by their transcriptional state inferred by VIPER, while fulfilling practical considerations [39]. The cell lines were obtained from ATCC and cultured using prescribed conditions to the amounts as described below.

## IC_20_ determination

To determine the 48h IC_20_ of each drug, cell lines were plated into 96well tissue culture plates, in 100 *µ*L total volume, and incubated at 37°C. After 16 hours the plates were removed from the incubator and compounds were transferred into assay wells (1 *µ*L) in triplicate. Plates were then returned to the incubator. After 48 hours the assay plates were removed from the incubator and allowed to cool to room temperature prior to the addition of 100 *µ*L of CellTiter-Glo (Promega Inc.) per well. The plates were then mechanically shaken for 5 minutes prior to readout on the EnVision Multi-Label Reader (Perkin Elmer Inc.) using the enhanced luminescence module. Relative cell viability was computed using matched Thimerosal control wells as reference. IC_20_ was estimated by fitting a four parameter sigmoid model to the titration results.

## Drug perturbation of cell lines

All cell lines were perturbed with the seven drug compounds and DMSO control for the corresponding time points, 5min, 15min, 1h, 6h, 24h, 48h and 96h. Treatment concentrations for each targeted drug compound were selected to be at maximum 0.5*µ*M and lower than the FDA-approved C_max_, as well as lower than the IC_20_ value of the most sensitive cell line of our panel as determined above: Alpelisib (BYL719): 0.12*µ*M, Imatinib (STI571): 0.5*µ*M, Linsitinib (OSI-906): 0.14*µ*M, Osimertinib (AZD9291): 0.5*µ*M, Ralimetinib (LY2228820): 0.5*µ*M, Trametinib (GSK1120212): 0.036*µ*M, WIKI4: 0.5*µ*M and DMSO: 0.5%. Each cell line was plated in 6-well plates in numbers that would approach confluency by 96h for the fastest growing cell line. After overnight attachment, cells were treated with the drugs at concentrations described and for the time points needed, and then lysed and processed as described below. Each sample was run in triplicate. For baseline phosphoproteomic profiling of the cell lines, cell lines were grown in 150 mm × 25 mm dishes to about 80% confluency and split into 3 batches. At time of harvest, cells were washed 3x with PBS, pelleted, snap-frozen by liquid nitrogen and stored at -80°C.

## Proteomic sample preparation

Cells were lysed and digested, mainly as previously described [102, 103]. For frozen cell pellets, cells were lysed on ice by adding 10 M urea containing complete protease inhibitor cocktail (Roche) and Halt™ Phosphatase Inhibitor (Thermo) to the pellets, pellets well resuspended, and processed for tryptic digestion. For cells in 6-well plates, plates were washed 3x with pre-cooled PBS and cells in wells lysed on ice immediately in 10 M urea containing complete protease inhibitor cocktail (Roche) and Halt™ Phosphatase Inhibitor (Thermo) and lysates stored at -80°C until for further analysis. Lysates were processed for tryptic digestion as follows. The cell pellets/lysates were further subjected to sonication at 4°C for 2 min using a VialTweeter device (Hielscher-Ultrasound Technology) and then centrifuged at 18,000 × g for 1 h to remove the insoluble material. A total of 300-500 *µ*g supernatant proteins (determined by BioRad Bradford assay) were transferred to clean Eppendorf tubes. The supernatant protein mixtures were reduced by 10 mM tris-(2-carboxyethyl)-phosphine (TCEP) for 1 h at 37°C and 20 mM iodoacetamide (IAA) in the dark for 45 min at room temperature. Then five volumes of precooled precipitation solution containing 50% acetone, 50% ethanol, and 0.1% acetic acid were added to the protein mixture and kept at -20 °C overnight. The mixture was centrifuged at 18,000 × g for 40 min. The precipitated proteins were washed with 100% acetone and 70% ethanol with centrifugation at 18,000 × g, 4°C for 40 min, respectively. The protein pellets were dried in SpeedVac for 5 min. 300 *µ*L of 100 mM NH_4_HCO_3_ was added to all samples, which were digested with sequencing grade porcine trypsin (Promega) at a ratio of 1:20 overnight at 37°C. After digestion, the peptide mixture was acidified with formic acid and then desalted with a C18 column (MarocoSpin Columns, NEST Group INC). The amount of the final peptides was determined by Nanodrop (Thermo Scientific). About 5% of the total peptide digests were kept for total proteomic analysis of the cell line baseline profiles.

## Phosphoproteomic sample preparation

From the same peptide digest above, ^∼^95% of peptides per sample was used for phosphoproteomic analysis. The phosphopeptide enrichment was performed using the High-Select™ Fe-NTA kit (Thermo Scientific, A32992) according to the kit instruction, as described previously [104]. Briefly, the resins of one spin column in the kit were divided into five equal aliquots, each used for one sample. The peptide-resin mixture was incubated for 30 min at room temperature and then transferred into the filter tip (TF-20-L-R-S, Axygen). The supernatant was removed after centrifugation. Then the resins adsorbed with phosphopeptides were washed sequentially with 200 *µ*L× 3 washing buffer (80% ACN, 0.1% TFA) and 200 *µ*L×3 H2O to remove nonspecifically adsorbed

peptides. The phosphopeptides were eluted off the resins by 100 *µ*L×2 elution buffer (50% ACN, 5% NH3 H2O). All centrifugation steps above were conducted at 500 g, 30 sec. The eluates were collected for speed-vac and dried for mass spectrometry analysis.

## Mass spectrometry data acquisition

For each proteomic and phosphoproteomic sample generated above, DIA-MS analysis was performed on 1 *µ*g of peptides, as described previously [103, 105].

Briefly, LC separation was performed on EASY-nLC 1200 systems (Thermo Scientific, San Jose, CA) using a self-packed analytical PicoFrit column (New Objective, Woburn, MA, USA) (75 *µ*m × 50 cm length) using C18 material of ReproSil-Pur 120A C18-Q 1.9 *µ*m (Dr. Maisch GmbH, Ammerbuch, Germany). A high-throughput 75-min measurement with buffer B (80% acetonitrile containing 0.1% formic acid) from 6% to 37% and corresponding buffer A (0.1% formic acid in H2O) during the gradient was used to elute peptides from the LC. The flow rate was kept at 300 nL/min with the temperature-controlled at 60°C using a column oven (PRSO-V1, Sonation GmbH, Biberach, Germany).

The Orbitrap Fusion Lumos Tribrid mass spectrometer (Thermo Scientific) instrument coupled to a nanoelectrospray ion source (NanoFlex, Thermo Scientific) was calibrated using Tune (version 3.0) instrument control software. Spray voltage was set to 2,000 V and heating capillary temperature at 275°C. All the DIA-MS methods consisted of one MS1 scan and 40 MS2 scans of variable isolated windows [105], with1 m/z overlapping between windows. The MS1 scan range is 350 – 1650 m/z, and the MS1 resolution is 120,000 at m/z 200. The MS1 full scan AGC target value was set to be 2.0E5, and the maximum injection time was 100 ms. The MS2 resolution was set to 15,000 at m/z 200 with the MS2 scan range 200–1800 m/z, and the normalized HCD collision energy was 28%. The MS2 AGC was set to be 5.0E5, and the maximum injection time was 50 ms. The default peptide charge state was set to 2. Both MS1 and MS2 spectra were recorded in profile mode. Detailed MS settings can be inspected through raw files provided via ProteomeXchange.

## Mass spectrometry data analysis

All raw data files were processed and converted to mzXML by ProteoWizard [106] (version 3.0), enabling centroiding (using the vendor-provided algorithm) on MS1 and MS2 levels. For peptide identification and quantification, an integrated Snakemake workflow consisting of DIA-Umpire [107, 108] (version 2.1.6), MSFragger [109] (version 2.3.0), the Trans-Proteomic Pipeline (PeptideProphet [110, 111], PTMProphet [112], iProphet [113], version 5.2.0), EasyPQP (version 0.1.6), OpenSWATH [26] (OpenMS [114], version 2.5.0), PyProphet [57, 115] (version 2.1.4) and TRIC [116] (msproteomicstools, version 0.11.0) was used.

A UniProtKB/Swiss-Prot protein sequence database was used for MSFragger. The spectral library was controlled to 1% PSM-, peptide-and protein-level FDR in global context and the best site-localization per phosphosite was selected. EasyPQP exported a global library, as well as a sample-specific library for each run.

OpenSWATH was run using the sample-specific high confidence library for mass calibration and non-linear retention time alignment with enabled IPF [25] module for peptidoform-level confidence estimation. PyProphet with enabled IPF module and using the XGBoost classifier [117] was used to for statistical validation. Peptides and proteins were filtered to 1% FDR in global context. TRIC was used for feature alignment using the IPF peptidoform-level scores in run-specific context, aligning detected peptides by lowess with a seed FDR of 1% to a maximum of 5%.

For quantitative protein abundance inference, the R-package “iq” [118] (version 1.9), implementing the MaxLFQ algorithm [98] for DIA-based datasets, was used with default parameters.

The full workflow, all used parameters and software distributed as Docker containers that enable accurate reproduction of the analysis are provided with the dataset via ProteomeXchange.

## CRISPRko validation experiment

### Cell culturing

- HCT-15 - RPMI 10%FBS + pen/strep
- NCI-H508 - RPMI 10%FBS + pen/strep
- 293T – DMEM 10%FBS + pen/strep

All cell lines were routinely tested for mycoplasma contamination. Cell lines were kept in a 37 °C humidity-controlled incubator with 5.0% CO2.

### Optimizing drug concentrations for pooled CRISPRko screens

Drug concentrations were optimized for each cell line to ensure ideal long-term CRISPRko screen readouts. The time to reach 10-population doublings depended primarily on characteristics of each cell line and could take between 25-40 days. Trametinib and linsitinib perturbations were tested with 5 concentrations (10*µ*M, 1*µ*M, 0.1*µ*M, 0.01*µ*M and DMSO only) and the cellular growth effect was assessed for each of those concentrations for each of the cell lines in a long-term growth assay. The DMSO concentration was optimized for 0.15%.

We let the cells grow in the presence of these drug perturbations in 15cm plate format, splitting the cells whenever they became approx. 80-90% confluent. When the DMSO-plate reached 10-population doublings, the total number of cell divisions were counted for each of the above-mentioned drug treatment plates. Final concentrations for the pooled CRISPRko-screens were selected to represent drug concentrations which had only a modest effect on cell division rate (approx. 10-20% slower cell divisions compared to DMSO), similarly as previously suggested [119].

### CRISPRko library design

For CRISPRko screening we designed the target gene list to include all human kinases (obtained from UniProt: pkinfam.txt), phosphatases (obtained from reference [8]) and E3-ligases (obtained from reference [120]), altogether 1101 genes. All these genes were targeted with 4 sgRNAs / gene. For guide designs we used CRISPick [121, 122].

### CRISPRko oligo synthesis and library cloning

Oligo libraries (4404 oligos) were ordered from Twist-biosciences in following format: cttgtggaaaggacgaaacaccgNNNNNNNNNNNNNNNNNNNN-gtttAagagctagaaatagcaagttTaaataaGgct

### Twist oligo pool amplifcation

- 1*µ*l Twist oligo library (1ng/ul)
- 10*µ*l 5x KAPA HIFI buffer
- 1 *µ*l dNTPs
- 1 *µ*l KAPA
- 2*µ*l sgRNA insert dd F (10*µ*M)
- 2*µ*l sgRNA insert dd R (10*µ*M)
- 2.5*µ*l 20xSYBR
- 30.5*µ*l H2O
- 95 °C 3 min
- 98 °C 20 sec (done with qPCR, stopped before saturation)
- 56 °C 15 sec (done with qPCR, stopped before saturation)
- 72 °C 20 sec (done with qPCR, stopped before saturation)
- 72 °C 5 min
- 4 °C *∞*
- sgRNA insert dd F: CTTGTGGAAAGGACGAAACACCG
- sgRNA insert dd R: AGCCTTATTTAAACTTGCTATTTCTAGCTCTTAAAC

After PCR, the insert was gel purified (GeneJet) and Golden-gate cloned into BsmBI-digested pLenti-guide-Puro (addgene #52963).

Golden-gate cloned insert + vector was Isopropanol precipitated and large-scale electroporated into Lucigen Enduro competent cells. The bacterial colonies were scraped from 10 x 24,5cm x 24,5cm agar plates, so that the estimated library complexity was *>* 1000 colonies / sgRNA.

### CRISPRko library viral packaging

13 million 293T cells were seeded for each 15cm dish previous night of the transfections. The following morning the viral transfections were conducted the following way: • 22.1*µ*g sgRNA-library containing pLenti-guide-Puro or pLenti-Cas9-Blast

- 16.6*µ*g PsPAX2 (Addgene 12260)
- 5.5*µ*g PMD2G (Addgene 8454)
- 1660*µ*l of sterile H_2_O

After mixing the plasmids 110,6*µ*l of Fugene HD (Promega) was added to the mix.

The transfection mixture was briefly vortexed and incubated 10 minutes in room temperature before adding dropwise to 293T cells. Altogether 3 x 15cm plates were transfected for sgRNA-library containing pLenti-guide-Puro and 1 x 15cm plates were transfected for pLenti-Cas9-Blast (addgene #52962).

The transfection mixture was removed the following day and virus was collected at 48h and 72h after initial transfections. To remove cellular debris, the virus containing supernatant was centrifuged 500 x g for 5min and filtered by using 0.45*µ*m PES filters (Millipore). The lentivirus was concentrated by using Lenti-X concentrator (Clontec), aliquoted and stored at -80 °C.

### Generation of Cas9 expressing CRC cell lines

Cas9 expressing cell lines were generated as follows: Concentrated pLenti-Cas9-Blast-lentivirus was transduced to CRC cell lines (in presence of 8*µ*g/ml polybrene) with estimated MOI 0.3. The virus was removed the following day and 4*µ*g/ml Blasticidin was added to the cells. Blasticidin selection was continued as long as the control cells (non-transduced) were viable.

### CRISPRko screening

sgRNA containing lentiviruses were transduced into Cas9 expressing CRC cell lines (in 15cm plate-format) in quadruplicates (in presence of 8*µ*g/ml polybrene), at an estimated MOI = 0.2. After 24h, the lentivirus containing media was removed, cells were washed with PBS, and puromycin-containing media (3*µ*g/ml) was added to the cells for 48-96h until all control cells (not virus-infected) were dead. After this the cells were cultured for extra couple of days so that the plates reached approx. 80% confluency. At this point the cells were divided into 3 parts; 1/3 going into -80 °C as time point 1 to assess sgRNA representation baseline, 1/3 to continue to culture with DMSO and 1/3 to continue to culture with either with Linsitinib or Trametinib. Cells were always maintained at *>*1,500 cells per guide throughout the screens and finally harvested after 10 population doublings to assess gene essentiality. The exact time (in days) for this varied for DMSO / Linsitinib / Trametinib with different cell lines. After the screen, the genomic DNA from the first and the last timepoints (DMSO & Drug perturbed) were extracted by using Blood and Cell culture DNA Maxi kits (Qiagen).

### Preparation of the sequencing library from genomic DNA

NGS library preparations were done the following way: Briefly, 40 *µ*g of gDNA, theoretically corresponding to 6 million diploid cells, was used as PCR template in 4 parallel NGS PCR1 reactions (10 *µ*g template DNA per reaction) using ExTaq DNA polymerase (Takara bio). After 18 cycles, the 4 replicate reactions were pooled together. 2 *µ*l of pooled NGS PCR1 product was used as template for NGS PCR2 which was run with qPCR with index primers and stopped before the amplification started to saturate. The resulting products of approx. 360 bp were gel purified (GeneJet), pooled together and Next generation sequenced.

### NGS PCR1 master mix

- 10*µ*g gDNA
- 0.75*µ*l ExTaq
- 10*µ*l 10 x ExTaq Buf
- 8*µ*l dNTPs
- 0.5*µ*l CRISPRko PCR 1R (pool of 5 (100*µ*M))
- 0.5*µ*l CRISPRko PCR 1F (100*µ*M)
- to 100*µ*l H_2_O

### PCR1 protocol

- 98 °C 1 min
- 98 °C 10 sec (18 cycles)
- 58 °C 30 sec (18 cycles)
- 72 °C 30 sec (18 cycles)
- 72 °C 10 min
- 4 °C *∞*

### NGS PCR2 master mix

- 2*µ*l DNA (from 1PCR)
- 0.375*µ*l ExTaq
- 5*µ*l 10 x ExTaq Buf
- 4*µ*l dNTPs
- 0.5*µ*l CRISPRko PCR 2F (100*µ*M)
- 0.5*µ*l CRISPRko PCR 2R(index) (100*µ*M)
- 1.25*µ*l 20xSYBR
- 36.4*µ*l H_2_O

### PCR2 protocol

- 98 °C 1 min
- 98 °C 10 sec (done with qPCR, stopped before saturation)
- 60 °C 30 sec (done with qPCR, stopped before saturation)
- 72 °C 30 sec (done with qPCR, stopped before saturation)
- 72 °C 10 min
- 4 °C *∞*

### CRISPRko Oligos used for NGS library preparation

- CRISPRko PCR 1F: TGGAGTTCAGACGTGTGCTCTTCCGATCTTCTAC-TATTCTTTCCCCTGCACTGT
- CRISPRko PCR 1R: CTTTCCCTACACGACGCTCTTCCGATCT(1-5nt stagger)TGTGGAAAGGACGAAACACCG
- CRISPRko PCR 2F: AATGATACGGCGACCACCGAGATCTACACTCTTTCC-CTACACGACGCTCTTCCGATCT
- CRISPRko PCR 2R(index): CAAGCAGAAGACGGCATACGA-GATNNNNNNNNGTGACTGGAGTTCAGACGTGTGCTCTTCCGATCT

## Data processing & Statistical analysis

For all data analysis steps, “viper” (version 1.22.0), “vespa” (version 1.0.2), “vespa.db” (version 1.0.2) and “vespa.aracne” (version 2.2) were used. “vespa.net” (version 1.0.2) was executed using the corresponding Docker images of the algorithms converted to Singularity images. All software tools are available from the corresponding repositories as referred below.

### Inference of a CRC-specific signaling network

To generate a CRC-specific signaling network, we obtained the processed phosphoproteomic and total proteomic profiles from the CPTAC study S045 [36] (referred to as “CPTAC-S045”). To account for potential confounding factors originating from protein abundance levels, we further generated a derived dataset, referred to as “CPTAC-S045N”, normalizing phosphopeptide abundance by the corresponding protein-level intensity values. The datasets were imported from CCT and CPTAC formats and converted to PVL by the corresponding “vespa” functions without further processing except mapping of identifiers. Only tumor samples were used across all analyses.

The phosphoproteomic dataset generated in this study (referred to as “U54”) was imported from the OpenSWATH file format and converted to PVL by the corresponding “vespa” function, with quantile normalization grouped by cell line and centering enabled. The baseline profiles of the six cell lines measured in triplicates (“U54-BL”), as well as drug perturbations across three distinct time points (1h, 24h, 96h; “U54-NET”) and the full time series (”U54-DP”) were exported as separate PVL files.

These three PVL sets (CPTAC-S045, CPTAC-S045N and U54-NET) were used as input to the “vespa.net” workflow. By default, separate signalons were generated for the stDPI/DPI, LP [14] (published dataset), HSM/P [13] (published dataset) and PC [16] (version 12) methods. For all analyses, the PVL of U54 BL was used to

generate optimized signalons. For all analysis, except the benchmark and the DeMAND analysis, the stDPI/DPI signalons were used. For the DeMAND analysis, the HSM/P signalons were used to allow restrictions on the prior confidence.

### Benchmark and validation of dVESPA

#### Benchmark signaling network generation

Signaling networks based on different data completeness thresholds of the U54-NET datasets were generated as described above.

#### Comparison of MI methods

To compare the MI estimation methods, the HSM priors were used as ground truth, as provided by the vespa.db R-package. Based on the U54-NET datasets, subsets were generated covering *>*20%, *>*40%, *>*60%, *>*80% and 100% data completeness. To compute hpMI and dMI, the sparse input matrices were directly used, to compute iMI, missing values were imputed row-wise, as described above. To compute dMI, “vespa.aracne” was extended to support depletion (Git branch “depletion support”; revision 470944f). “vespa.aracne” was run as described above, but without stDPI/DPI and using 100 bootstraps. Only significant interactions (*<*5% FDR) were considered. The overlap of these interactions with HSM was used to compute the summed score.

#### Comparison of DPI methods

stDPI, DPI and noDPI signaling networks were generated from U54-NET as described. Further, ground truth interactions were selected if they were identified as ST-K→S pairs by PDZ, SH3, WH1, and WW domain HSM analysis, since these represent the primary determinants of specific serine and threonine phosphopeptide interactions with ST-Ks. As a negative gold standard, we used candidate TK→S interactions with an inferred PTB, PTP and SH2 domain interaction with phosphotyrosine-containing peptide, since the used dataset for this benchmark (U54-NET), which is not enriched for phosphotyrosine peptides, should not be able to identify these interactions. This resulted in a context-specific reference dataset that separates between very likely direct and indirect interactions suitable for methodological comparisons. Receiver-Operating-Characteristics (ROC) curves were generated using the pROC R-package (version 1.17.0.1) and default parameters for each signaling network separately. P-values for ROC curve comparisons were also computed using pROC by DeLong’s test and using default parameters.

### Benchmark and validation of mVESPA

To benchmark mVESPA, we used the phosphoproteomic cell line baseline profiles (U54-BL) dataset as described above and obtained the curated GDSC [38] drug sensitivity and the primary target list from the original INKA publication [19] (Dataset EV6.xlsx).

Substrate-and activity-level signalons were generated as described above, however reference databases, datasets and parameters were disabled in a combinatorial fashion for the benchmark. Only those regulators present in all substrate-or activity-level comparisons were considered. For the differential comparison, “viperSignature” of the “viper” R-package, comparing sensitive *vs.* resistant or insensitive cell lines was used with default parameters. To assess the sensitivity of the recovered primary targets, the protein activity cumulative probability was weighted by the cell sensitivity cumulative probability: *S* = *pnorm* (*Z_V ESP A_*) * −*pnorm*(*Z_GDSC_*). A metric representing “specificity” was derived by transforming kinase activity ranks to relative activity ranks. The mean area-under-the-curve (AUC) metrics for each benchmark were computed as proposed previously, averaging the results of all differential comparisons [55]. Statistical comparison of the differential comparison AUC metrics were conducted using an unpaired, right tailed Wilcox’ tests (R-package “stats”, version 4.2.1).

### Representation of CRC subtypes by cell line models

#### Cell line selection

Cell lines were selected according to previously developed methods [39, 41]. The significance threshold for matching cell lines and patient samples was set to *p*-value *<* 10^-5^. To rank matching cell lines for each cluster, score 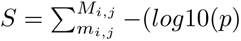, where *M_i,j_* represents all significant matches of cell line *j* to cluster *i*, was computed. This score was used to select six CRC cell lines, representing 5out of 8 (top 5) and all (top 10) clinical subtype clusters with at least one cell line.

#### CMS transcriptome-level classification

Preprocessed RNA-seq profiles for CPTAC S045 [36] were obtained from the original publication. Preprocessed RNA-seq profiles for the six CRC cell lines were obtained from CCLE [40]. Counts were normalized to TPM for both datasets and identifiers were mapped to be compatible with CMS. Only transcripts measured in both datasets were used for downstream analysis. The CMS classifier [43] was then applied using the RandomForest predictor and default parameters.

#### VESPA

Protein signaling activities were inferred by VESPA using the CRC-specific signalons described above. The phosphoproteomic profiles of CPTAC-S045 and U54-BL were first randomly subsampled on phosphosite-level to ensure that the detectability of phosphosites has the same frequency within both datasets. The two datasets were then concatenated and rank-normalized, first column-wise, then row-wise, as described previously [39]. The “viper” function was applied on substrate-and activity-levels including cross-talk correction, and statistically integrated, as described above.

#### Clustering

The substrate-and activity-level VESPA matrices were clustered by the k-medoids approach, prioritizing cluster robustness, as described previously [39].

#### Gene set enrichment analysis

GSEA was conducted by the R-package “fgsea” (version 1.14.0), using the Reactome pathway database (version 75) reduced to signaling-only gene sets (downstream pathway “R-HSA-162582”). Only significant results (adj. *p*-value *<* 0.05), belonging to primary pathways in at least one sample were reported.

#### Feature selection

Feature importance was assessed by applying the Random Forest recursive feature elimination method from the R-package “caret” (version 6.0-86), selecting the top 50 most important features for classification into the specific groups. For simplicity, Fig. 3b only shows the cumulative most important features of the CMS and pVC classification systems, grouped according to pVC. Supplemental Fig. 5-6 show the full results, whereas Supplemental Fig. 7 depicts the data underlying Fig. 3b, grouped according to CMS.

#### Visualization

Heatmaps were generated using the “*pheatmap*” (version 1.0.12) R-package. Hierarchical clustering on row-level was conducted using the default R “hclust” function with default parameters.

### Targeted drug perturbations of CRC cell lines

#### VESPA

The 336 perturbed U54-DP phosphoproteomic profiles were preprocessed by imputing missing values by using the row-wise minimum across all samples including numerical jitter to break ties as described above. To normalize for non-perturbation time-series and batch effects, peptide abundances of all samples were normalized by the corresponding DMSO controls for each cell line separately. Each time point was normalized by a sliding window of the average between the preceding and following time point, if available. E.g. 15min time points were normalized by the 5min, 15min and 1h time points of the corresponding DMSO control runs. The log2 fold changes were then used as input for all downstream steps. Protein signaling activities were inferred by VESPA using the stDPI/DPI CRC signalons as described above. The “viper” function was applied on substrate-and activity-levels using a bootstrapped “viperSignature” null model based on the DMSO controls, with 1000 permutations including cross-talk correction as described above. To infer activity-level signaling activities, the substrate-level activities were inferred without a null model but using the absolute log2 fold change values. Substrate-and activity-level VESPA results were integrated as described above.

#### Drug compound – cell line sensitivity analysis

Drug sensitivity data from GDSC [38] was obtained and *z* -score transformed per drug compound and GDSC dataset over all covered cell lines. Sensitive combinations were defined as z-score *<* -1.0, whereas insensitive combinations were defined as z-score *>* 1.0. Violin plots were generated using the “geom violin” function with default parameters of the R-package “ggplot2” (version 3.4.0).

#### Visualization

Heatmaps were generated using the “*pheatmap*” (version 1.0.12) R-package. Hierarchical clustering on row and column-level was conducted using the default R “hclust” function with default parameters.

#### Gene set enrichment analysis

GSEA was conducted by the R-package “fgsea” (version 1.14.0), using the Reactome pathway database (version 75) reduced to signaling-only gene sets (downstream pathway “R-HSA-162582”). Only significant results (adj. *p*-value *<* 0.05), belonging to primary pathways in at least one sample were reported.

*Temporal VESPA activity-level perturbation profiles of known primary drug compound targets:* Known primary targets for the drug compounds were obtained from DrugBank [58] and ProteomicsDB [59]. Only the top five most downregulated target proteins per drug compound were visualized.

### Context-specific wiring of signaling pathways

#### VESPA

The 336 perturbed U54-DP phosphoproteomic profiles were preprocessed as described above. The “viper” function was applied separately for each cell line on substrate-and activity-levels using a rank-normalized matrix, as described previously [39] and including cross-talk correction. Substrate-and activity-level VESPA results were integrated as described above.

#### DeMAND

The DeMAND [55] (version 1.18.0) algorithm was used to assess context-specific wiring of signaling pathways. Using substrate-level VESPA activities, the activity-level signalons were used. The STRING PPI DB (version 11) was used as reference interaction database on activity-level VESPA activities, considering all interactions with probability *>* 0.5. To generate subtype-unspecific DeMAND MoA profiles, for each drug perturbation, the temporal profiles of all cell lines were compared against the DMSO controls. To generate subtype-specific DeMAND MoA profiles, for each cell line and drug perturbation, the temporal profile was used as target and all DMSO controls were used as null distribution. Edge and node p-values were integrated using Fisher’s method and BH-adjusted for multiple testing.

#### Visualization

Heatmaps were generated using the “*pheatmap*” (version 1.0.12) R-package. Hierarchical clustering on row and column-level was conducted using the default R “hclust” function with default parameters.

#### Cytoscape

To visualize the interaction networks, Cytoscape (version 3.8.2) was used. Nodes indicate the most affected regulators with the inner circos colors indicating cell line type and the outer circos color and node size indicating VESPA activity. The edges indicate dysregulated, undirected interactions between the regulators. Line thickness indicates significance of dysregulation. Dysregulated nodes (BH-adjusted *p*-value *<* 0.05) and known primary targets are colored. Grey nodes indicate connecting dysregulated nodes (BH-adjusted *p*-value *<* 0.1).

### Context-specific adaptive stress resistance mechanisms

#### VESPA differential testing

Using the time series component of our experimental design, which covers drug perturbation time points from 5min to 96h, we investigated the adaptive response of kinases and phosphatases by comparing the late (24h, 48h, 96h) drug perturbed against DMSO time points using a paired, one-tailed t-test (R version 4.2.1). To select candidates for visualization, *p*-values were integrated by Stouffer’s method across all conditions and corrected for multiple testing (*q* -value *<* 0.05).

#### Identification of essential genes using DESeq2

Alignment of NGS with sgRNA guides was conducted using the “ShortRead” R-package (1.54.0). Essential genes for the DMSO *vs*. T0 comparison were obtained from a generalized resource [123], as well as aCRC-specific subset of DepMap [124], filtered to the 10% quantile of the gene effect. Differential expression analysis was conducted separately for each guide with the “DESeq2” R-package (1.36.0). *P* -values were integrated using Stouffer’s method and corrected for multiple-testing by the Benjamini-Hochberg FDR approach [125].

#### Receiver Operating Characteristics

ROC curves and statistics were generated using the R-package “pROC” (version 1.18.0). Significant (FDR *<* 0.01) CRISPRko results were used as ground truth values (negative beta: true; positive beta: false) and the VESPA scores (t-statistic) were used as predictors. ROC *p*-values were computed using the function “roc.area” from the R-package “validation” (version: 1.42).

#### Correlation analysis

Correlation analysis was conducted by comparing the t-statistic of the differential VESPA analysis with the significant (FDR *<* 0.01) log-fold-changes reported by DESeq2. Correlation statistics were computed using aone-tailed Spearman correlation test (R version 4.2.1).

#### Exclusion of tumor suppressor genes

For the analyses excluding tumor suppressor genes, all genes present in TSGene 2.0 database [91] were excluded.

#### Visualization

Heatmaps were generated using the “*pheatmap*” (version 1.0.12) R-package. The t-statistic values of the described above are visualized. Hierarchical clustering on row-and column-level was conducted using the default R “hclust” function with default parameters.

## Data availability

The CRC mass spectrometry proteomics data generated as part of this study have been deposited to the ProteomeXchange Consortium via the MassIVE partner repository (MassIVE) with the data set identifiers MSV000091204 / PXD039859.

The CRISPRko RNA-seq data discussed in this publication have been deposited in NCBI’s Gene Expression Omnibus [126] and are accessible through GEO Series accession number GSE224396 (GEO).

## Code availability

VESPA is available as modular platform-independent open-source software under a non-commercial usage license. VESPA consists out of five different modules, which are provided as versioned source code, binaries or docker containers.

- The “vespa” R-package for signaling protein activity inference is available from GitHub (https://github.com/califano-lab/vespa).
- The “vespa.db” R-package providing preprocessed reference networks is available from GitHub (https://github.com/califano-lab/vespa.db).
- The “vespa.aracne” algorithm with is available from GitHub (https://github.com/califano-lab/vespa.aracne).
- The “vespa.net” Snakemake workflow to generate context-specific signalons from one or multiple datasets is available from GitHub (https://github.com/califano-lab/vespa.net).
- A tutorial describing the full analysis workflow with example data is available from GitHub (https://github.com/califano-lab/vespa.tutorial).

## Supplementary Figures

**Supplementary Figure 1.**
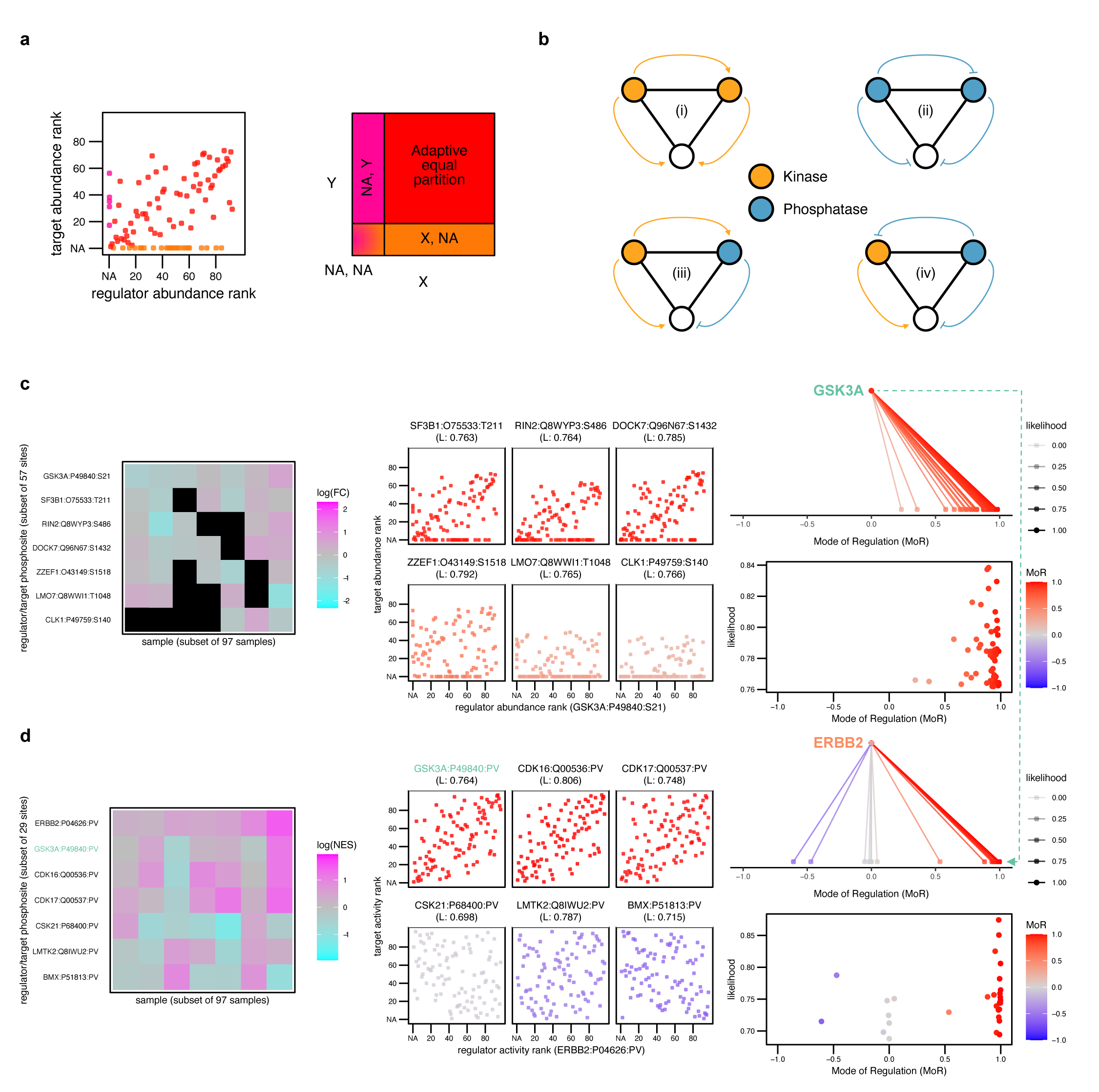
Walkthrough example of VESPA. **a)** Hybrid Partitioning computes Mutual Information (MI) by splitting data points into four quadrants, accounting for different levels of missing data (Methods). **b)** The signal transduction Data Processing Inequality (stDPI) accounts for the limited set of enzymatic actions inherent to kinases and phosphatases, restricting the valid triangular interactions between two enzymes and a substrate (Methods): Only valid regulator-regulator-substrate triangles are considered, either kinase-kinase-substrate (i) or phosphatase-kinase-substrate (iv) relationships, but not phosphatase-phosphatase-substrate (ii) or kinase-phosphatase-substrate (iii) relationships. **c)** An example on substrate-level illustrates the individual steps of VESPA, starting from the raw data (left), over mutual information and probabilistic weight estimation using regulator (GSK3A) and target (SF3B1, RIN2, DOCK7, ZZEF1, LMO7, CLK1) abundance ranks, accounting for missing values (middle), and visualization of the signalons including Mode of Regulation (MoR) and probabilistic weight. **d)** An example on activity-level illustrates how the results from VESPA on substrate-level are used to infer more abstract and generalized protein signaling activities, e.g. ERBB2, where the Data Processing Inequality allows for both activating and deactivating interactions of a kinase or phosphatase.

**Supplementary Figure 2.**
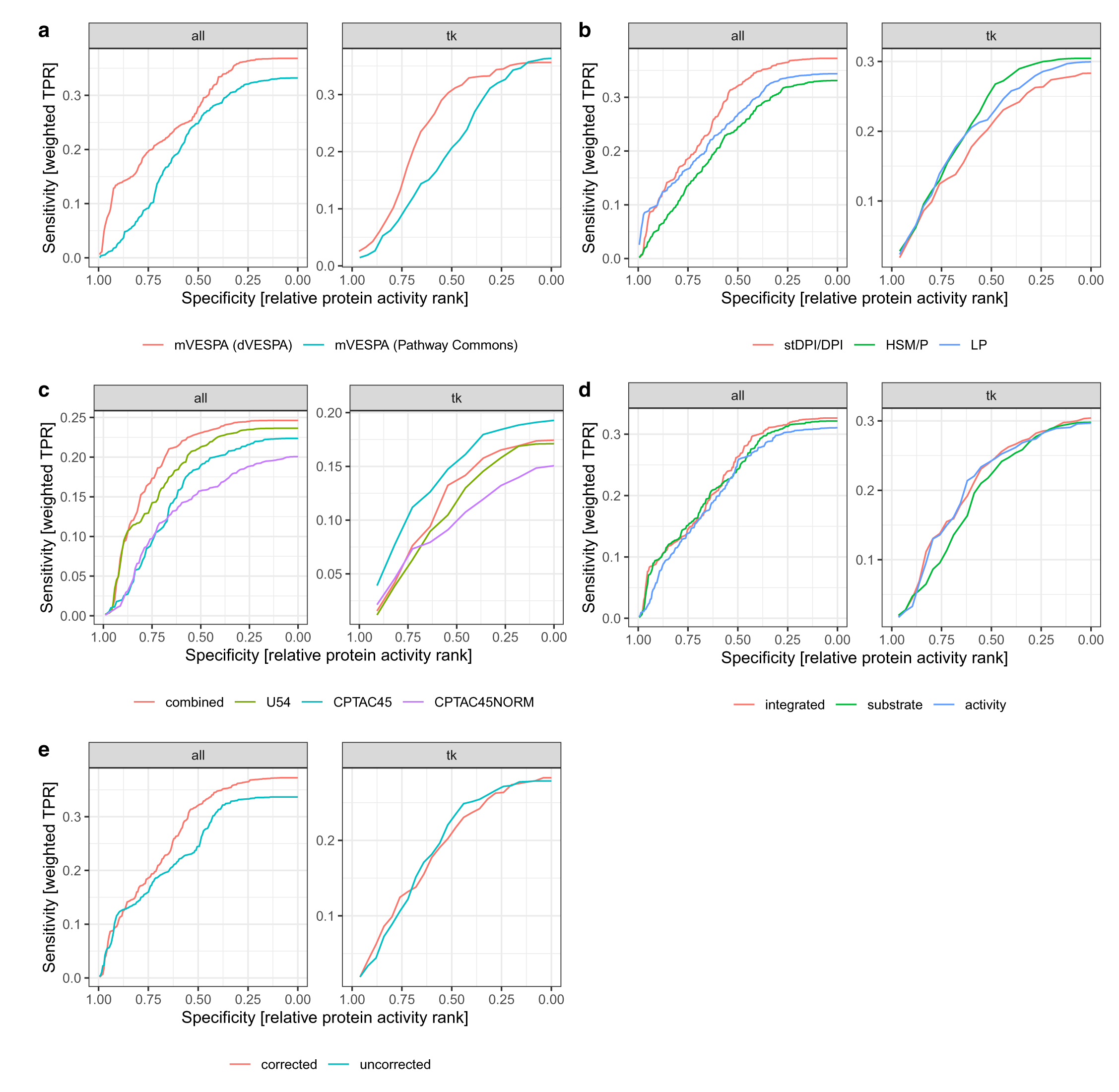
Benchmark and validation of mVESPA (intersection of signalons). Using VESPA and CRC-specific signalons, differential comparisons were conducted for each drug compound to identify the top differentially active regulators. Together with a list of the primary targets of all drug compounds, the sensitivity was computed based on the protein activity cumulative probability, weighted by the cell sensitivity cumulative probability, in dependency of the relative top ranking differential active proteins (selectivity). Computed area-under-the-curve (AUC) and *p*-values are listed in Supplemental Tables 1 and 2, respectively. **a)** Context-specific *vs.* reference-based signalons, **b)** Use of dVESPA-inferred signalons *vs.* reference constrained signalons, **c)** Signalon integration and optimization across multiple datasets, **d)** Hierarchical integration by mVESPA and **e)** mVESPA cross-talk correction.

**Supplementary Figure 3.**
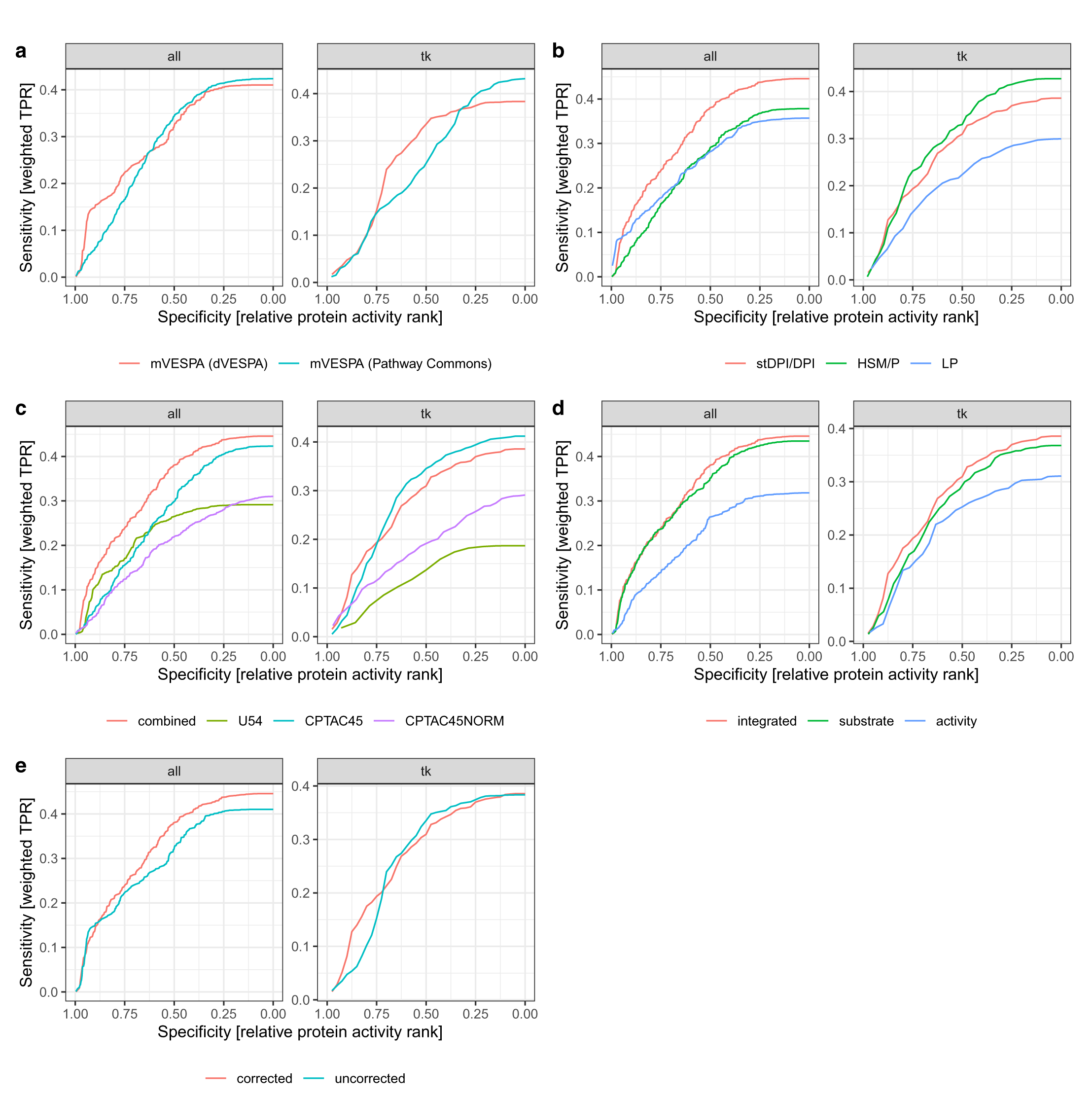
Benchmark and validation of mVESPA (full signalons). Using VESPA and CRC-specific signalons, differential comparisons were conducted for each drug compound to identify the top differentially active regulators. Together with a list of the primary targets of all drug compounds, the sensitivity was computed based on the protein activity cumulative probability, weighted by the cell sensitivity cumulative probability, in dependency of the relative top ranking differential active proteins (selectivity). Computed area-under-the-curve (AUC) and *p*-values are listed in Supplemental Tables 1 and 2, respectively. **a)** Context-specific *vs.* reference-based signalons, **b)** Use of dVESPA-inferred signalons *vs.* reference constrained signalons, **c)** Signalon integration and optimization across multiple datasets, **d)** Hierarchical integration by mVESPA and **e)** mVESPA cross-talk correction.

**Supplementary Figure 4.**
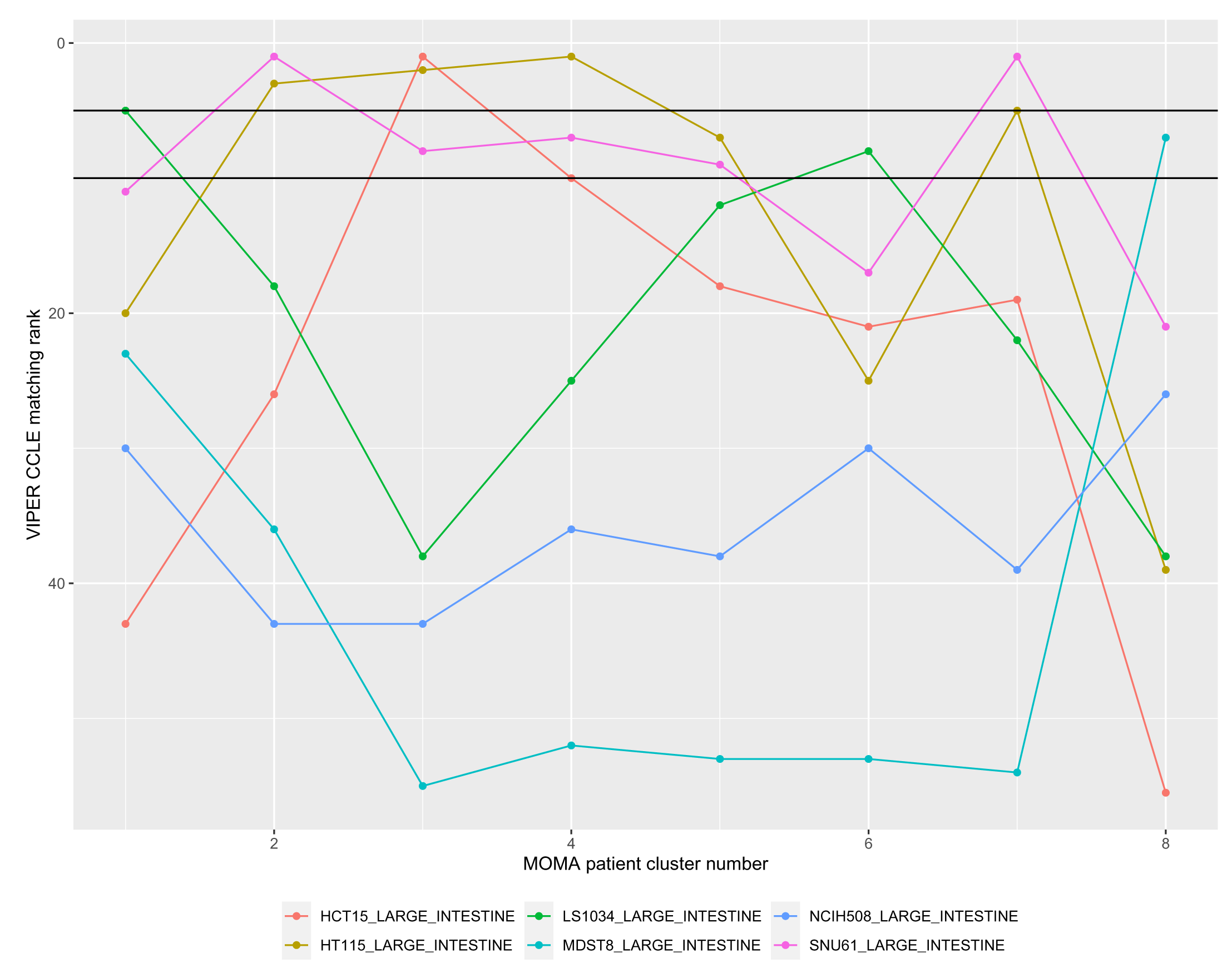
Representative CRC cell line selection. For each MOMA patient cluster, CRC CCLE cell lines were ranked according to their VIPER matching rank (Methods).

**Supplementary Figure 5.**
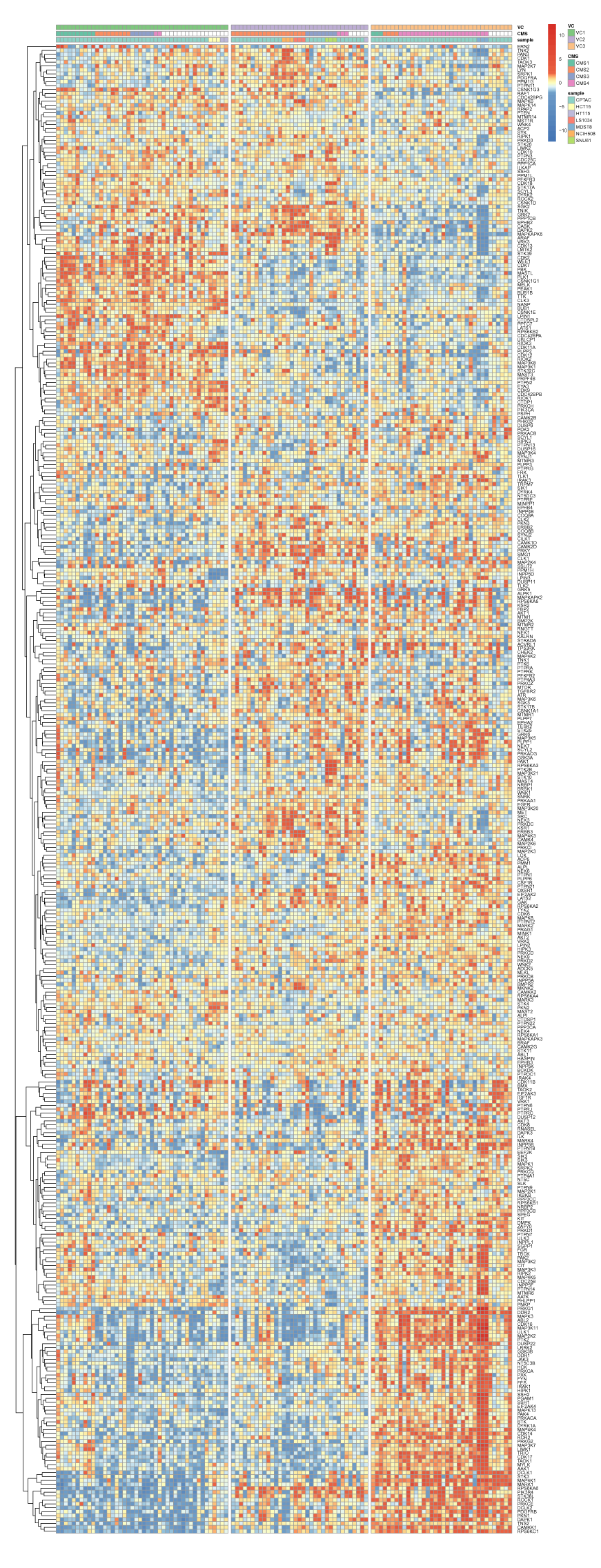
Representation of CRC subtypes by cell line models and the full VESPA matrix grouped according to VC. All regulators and their VESPA inferred normalized enrichment scores (NES) have been used for visualization. CPTAC clinical profiles and cell lines were grouped according to the Consensus Molecular Classifier (CMS) and VESPA clusters (VC). The samples are grouped according to VC.

**Supplementary Figure 6.**
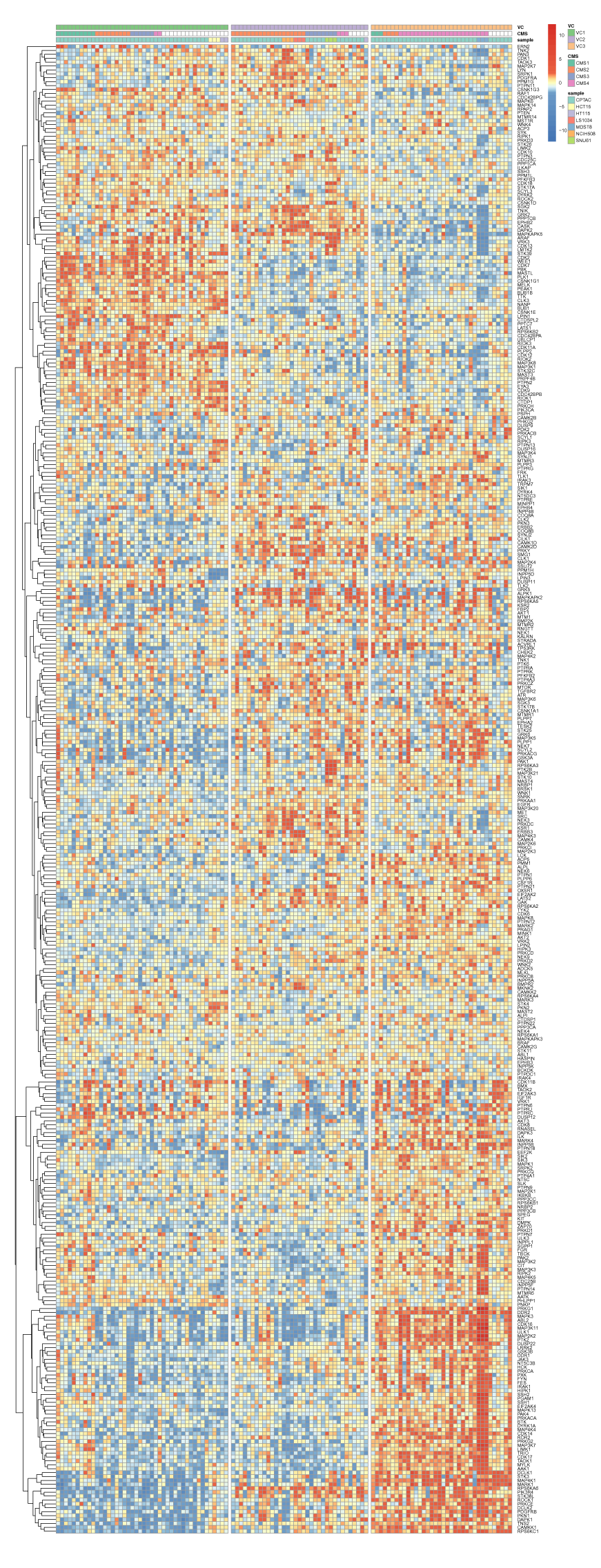
Representation of CRC subtypes by cell line models and the full VESPA matrix grouped according to CMS. All regulators and their VESPA inferred normalized enrichment scores (NES) have been used for visualization. CPTAC clinical profiles and cell lines were grouped according to the Consensus Molecular Classifier (CMS) and VESPA clusters (VC). The samples are grouped according to CMS.

**Supplementary Figure 7.**
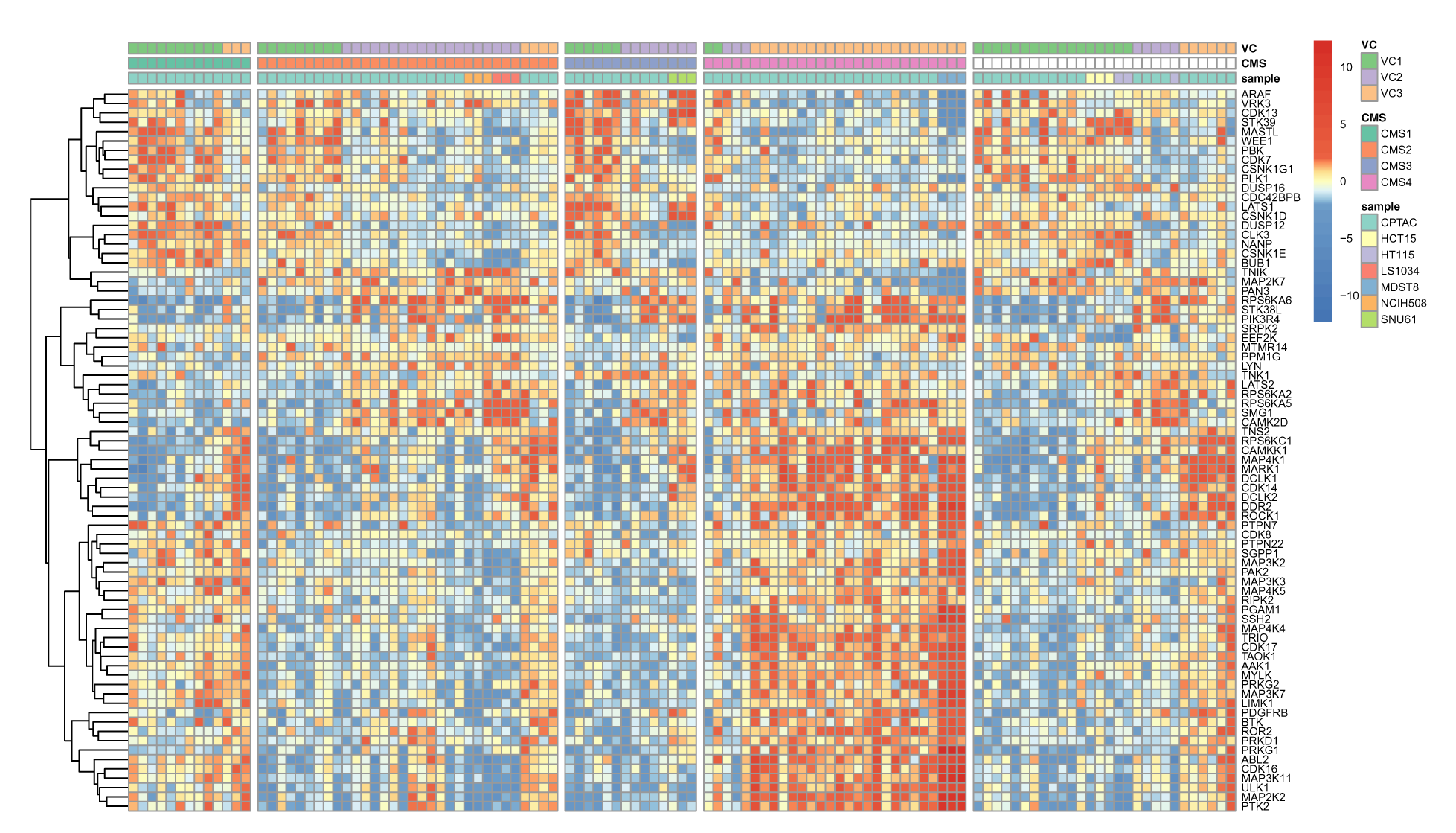
Representation of CRC subtypes by cell line models and VESPA grouped by CMS. The most informative proteins and their VESPA activity-level inferred normalized enrichment scores (NES) have been selected for visualization. CPTAC clinical profiles and cell lines were grouped according to the Consensus Molecular Classifier (CMS) and VESPA clusters (VC). The samples are grouped according to CMS.

**Supplementary Figure 8.**
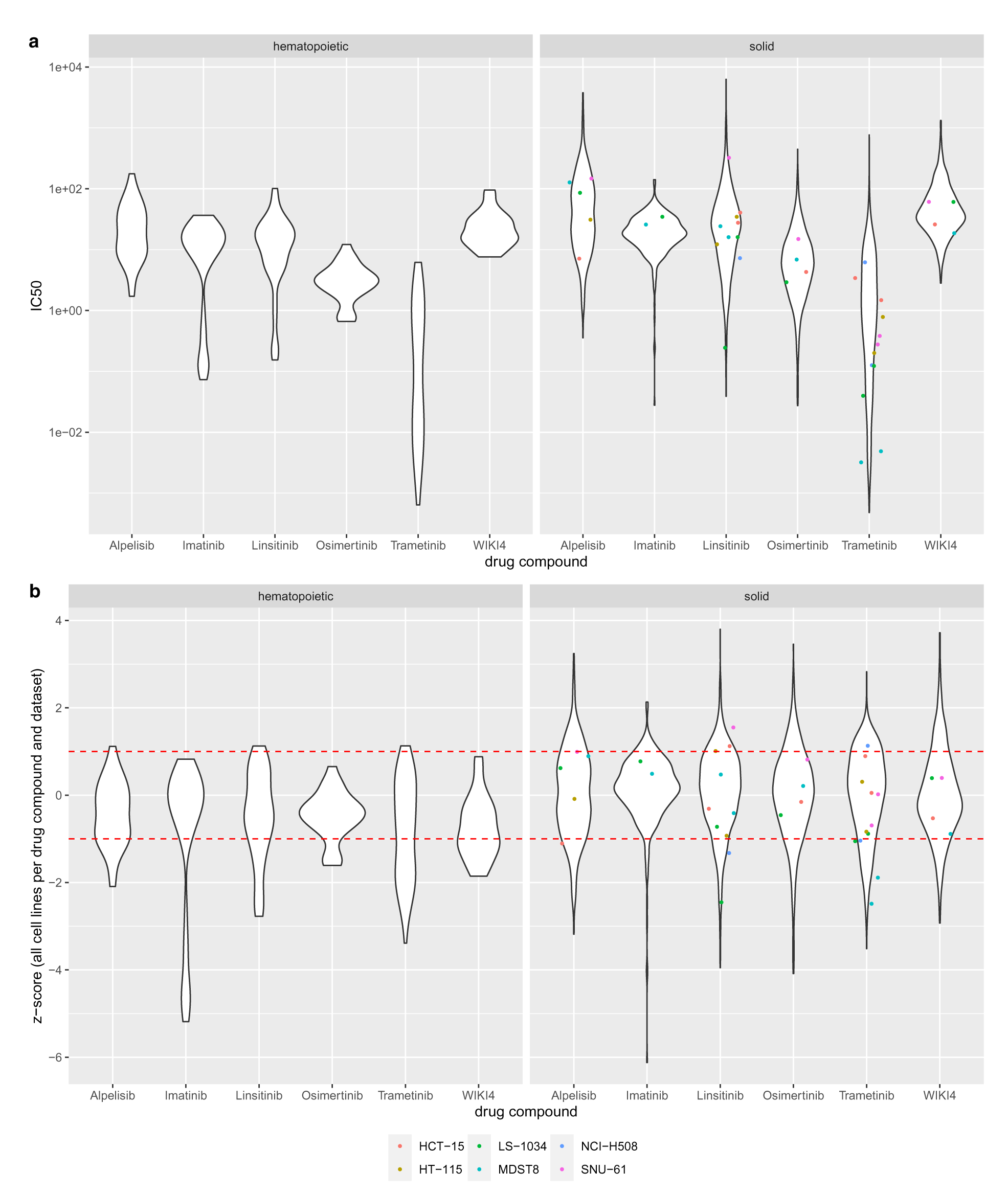
Targeted drug compound sensitivity of selected cell lines in context of GDSC. **a)** The distribution of GDSC 1&2 IC50 values is depicted as violinplots (Methods). **b)** The distribution of GDSC 1&2 *z* -score values, computed per drug compound over all cell lines per GDSC dataset, is depicted as violinplots (Methods). For both subfigures 33, 39, 69, 27, 64, and 27 datapoints were used to draw the distribution of hematopoietic tumors; 676, 325, 1417, 636, 1412, and 635 datapoints were used to draw the distribution of solid tumors. Red lines indicate thresholds for sensitive cell lines (z-score *<* -1.0) and resistant cell lines (z-score *>* 1.0)

**Supplementary Figure 9.**
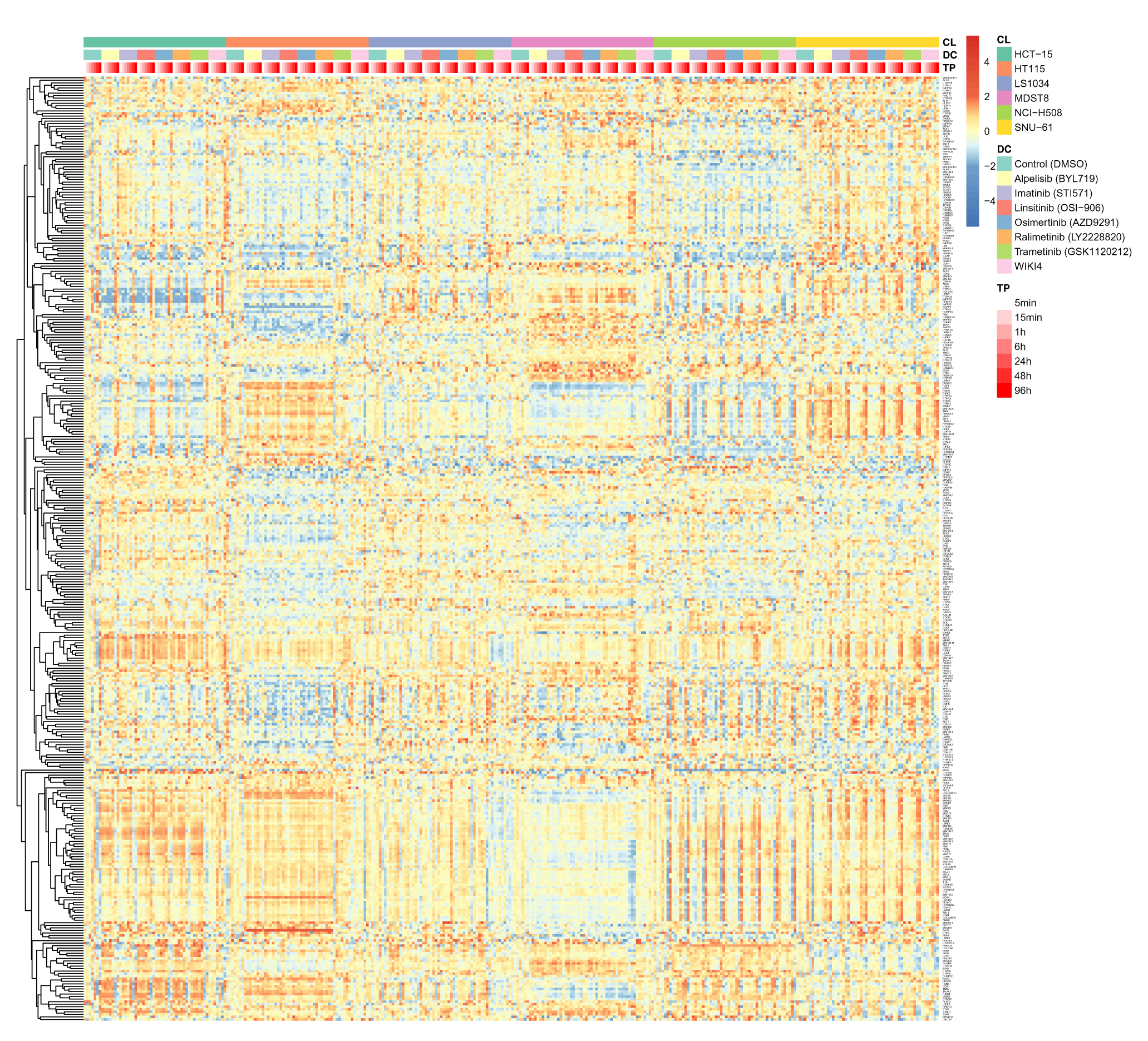
Targeted drug perturbations of CRC cell lines on substrate-level (sorted). A global overview of VESPA substrate-level inferred normalized enrichment scores (NES) across the full sorted drug perturbation dataset (336 samples), covering six CRC cell lines, 7 drug perturbations and DMSO control across 7 time points.

**Supplementary Figure 10.**
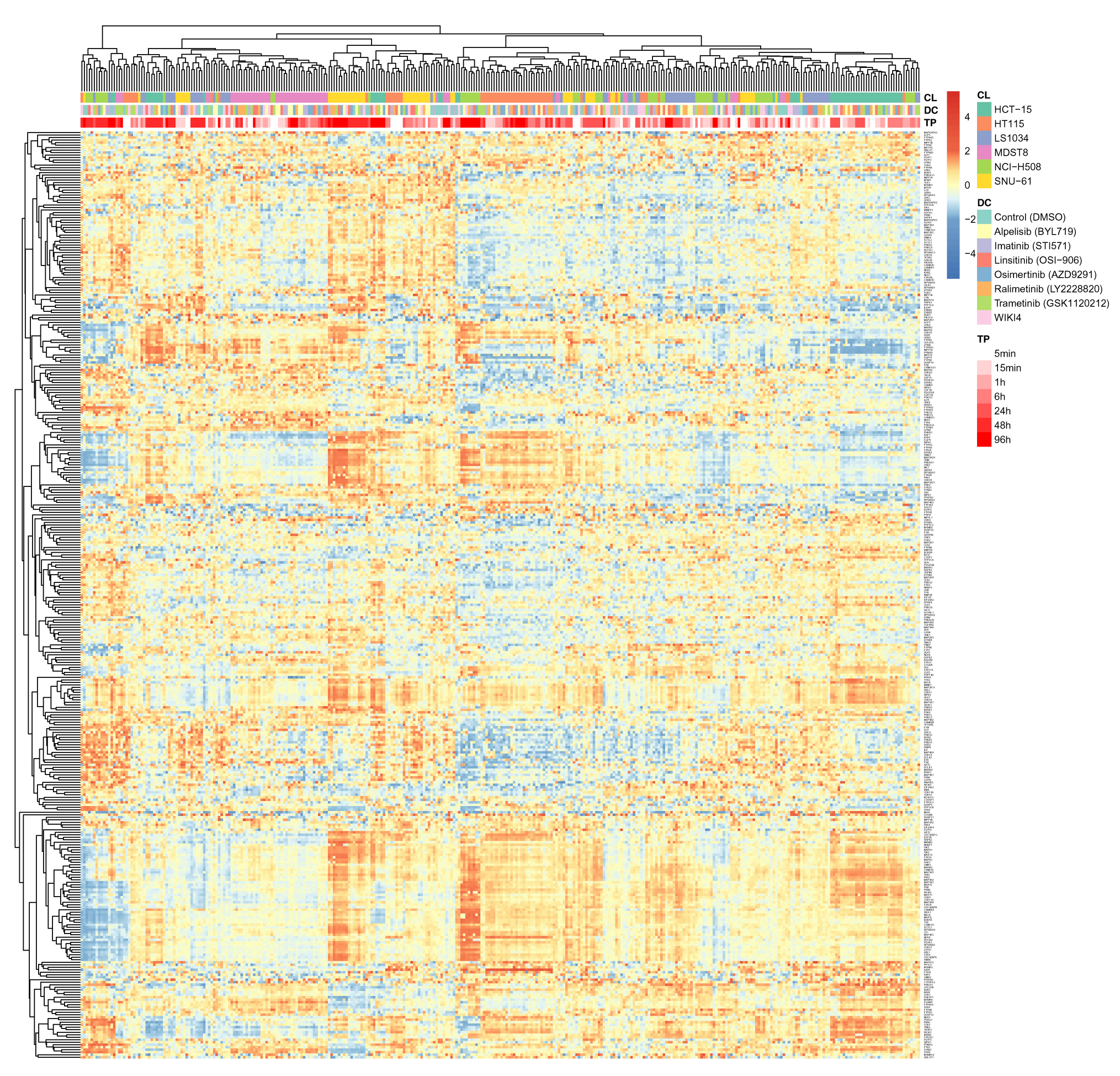
Targeted drug perturbations of CRC cell lines on substrate-level (grouped). A global overview of VESPA substrate-level inferred normalized enrichment scores (NES) across the full grouped drug perturbation dataset (336 samples), covering six CRC cell lines, 7 drug perturbations and DMSO control across 7 time points.

**Supplementary Figure 11.**
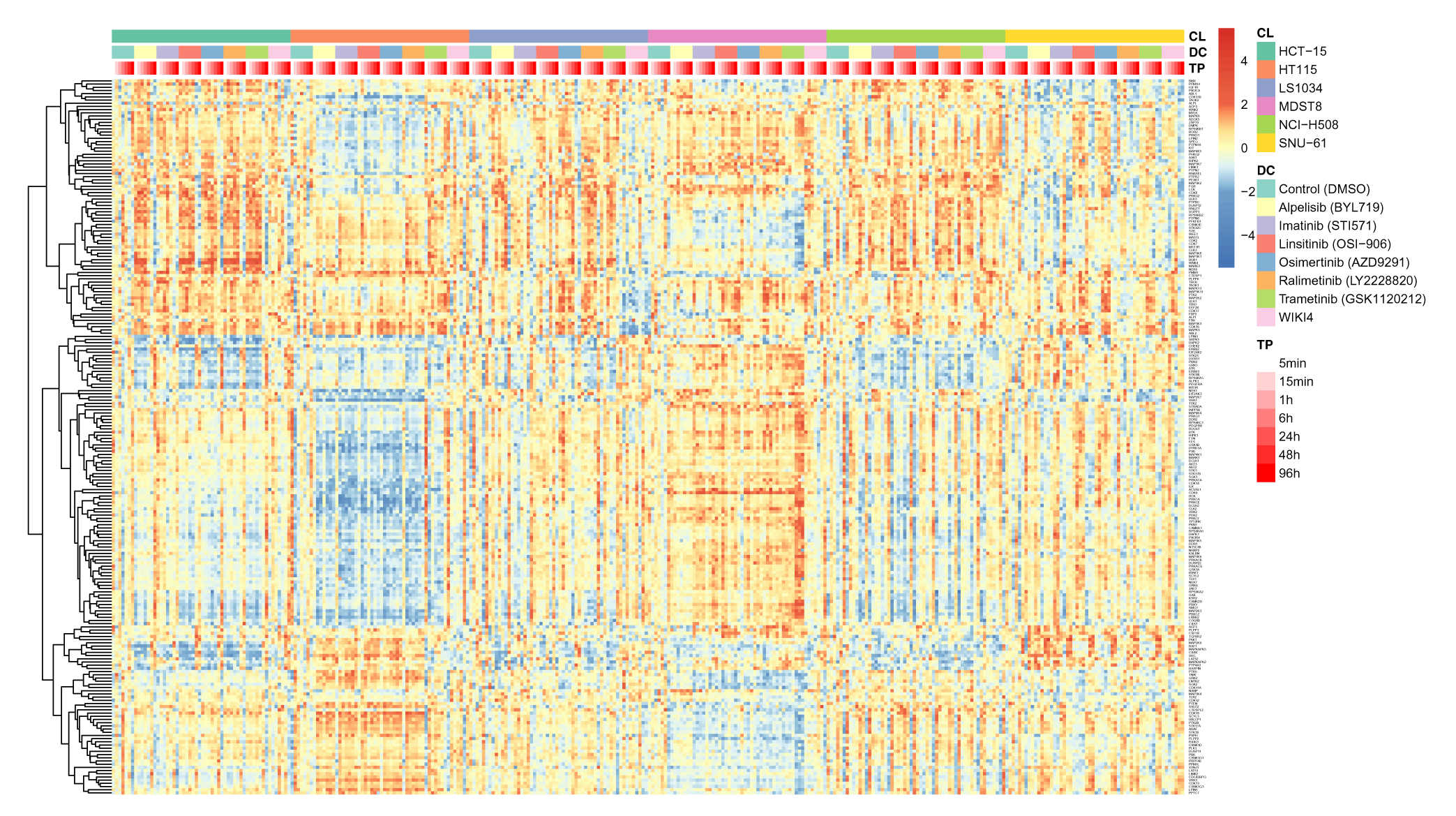
Targeted drug perturbations of CRC cell lines on activity-level (sorted). A global overview of VESPA activity-level inferred normalized enrichment scores (NES) across the full sorted drug perturbation dataset (336 samples), covering six CRC cell lines, 7 drug perturbations and DMSO control across 7 time points.

**Supplementary Figure 12.**
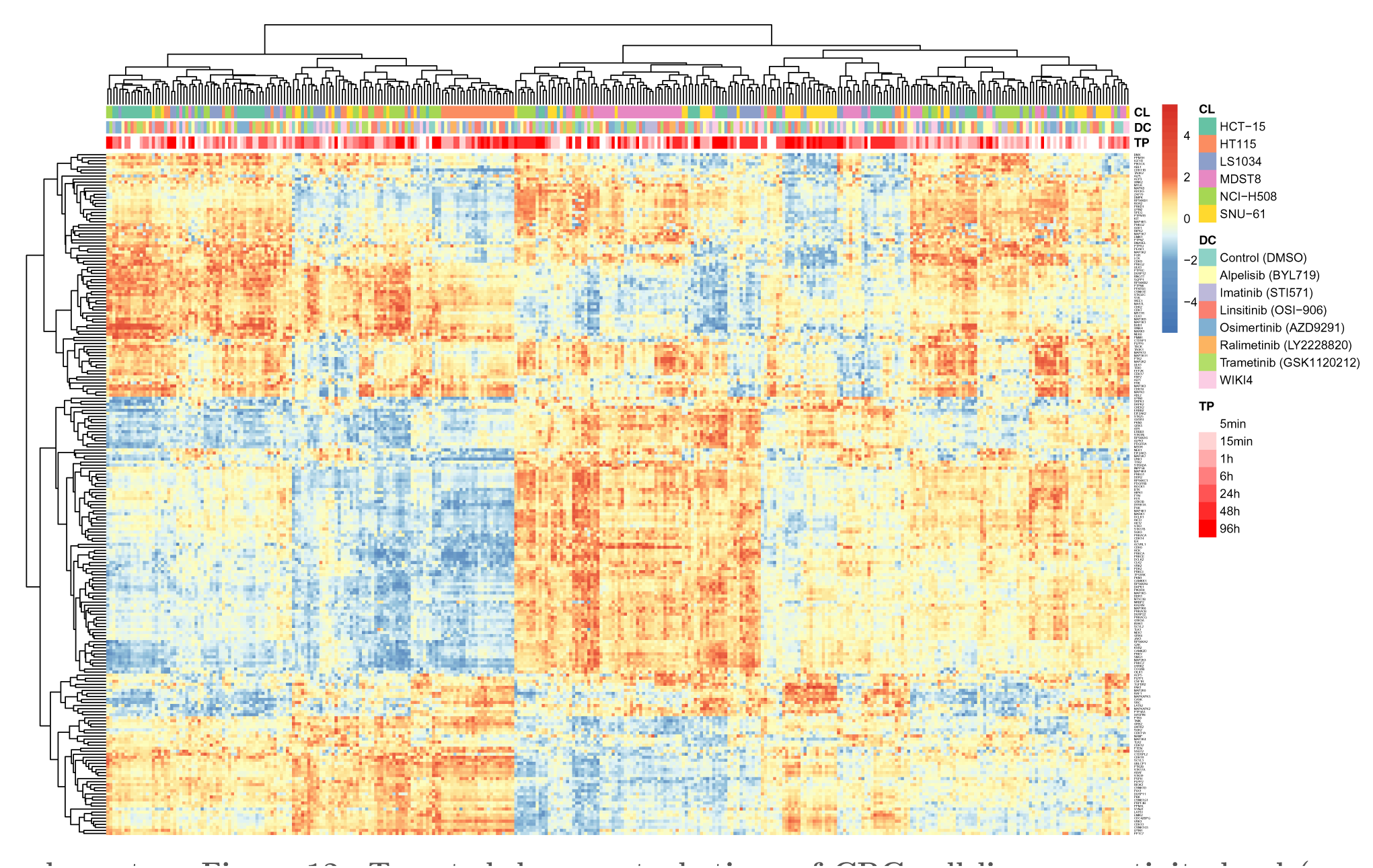
Targeted drug perturbations of CRC cell lines on activity-level (grouped) A global overview of VESPA activity-level inferred normalized enrichment scores (NES) across the full grouped drug perturbation dataset (336 samples), covering six CRC cell lines, 7 drug perturbations and DMSO control across 7 time points.

**Supplementary Figure 13.**
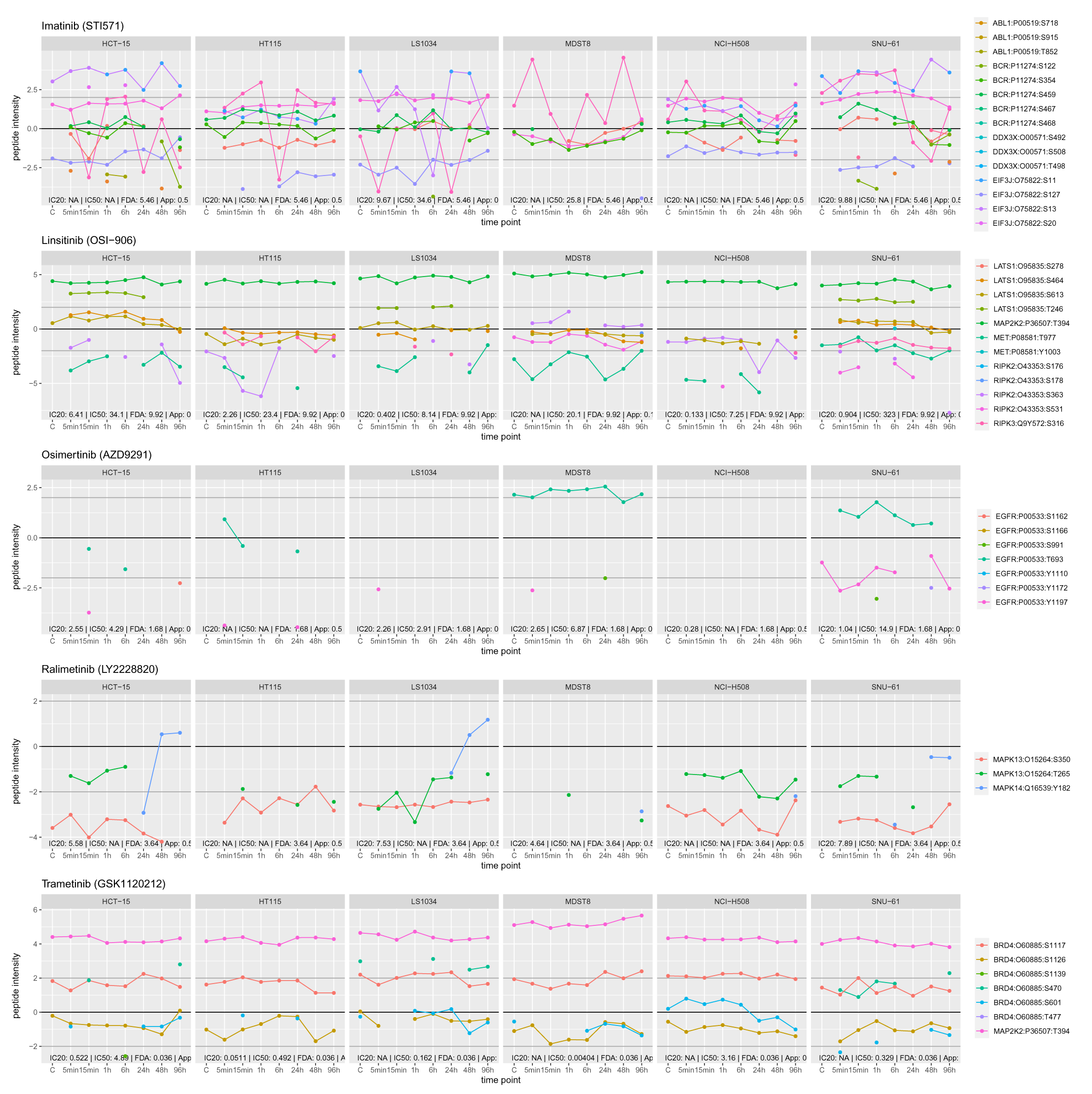
Temporal phosphosite abundance-level perturbation profiles of known primary drug compound targets. The phosphosite abundances corresponding to the quantitative matrix used as input for VESPA are extracted and visualized for the top 5 downregulated known primary targets, grouped according to drug perturbations and cell lines.

**Supplementary Figure 14.**
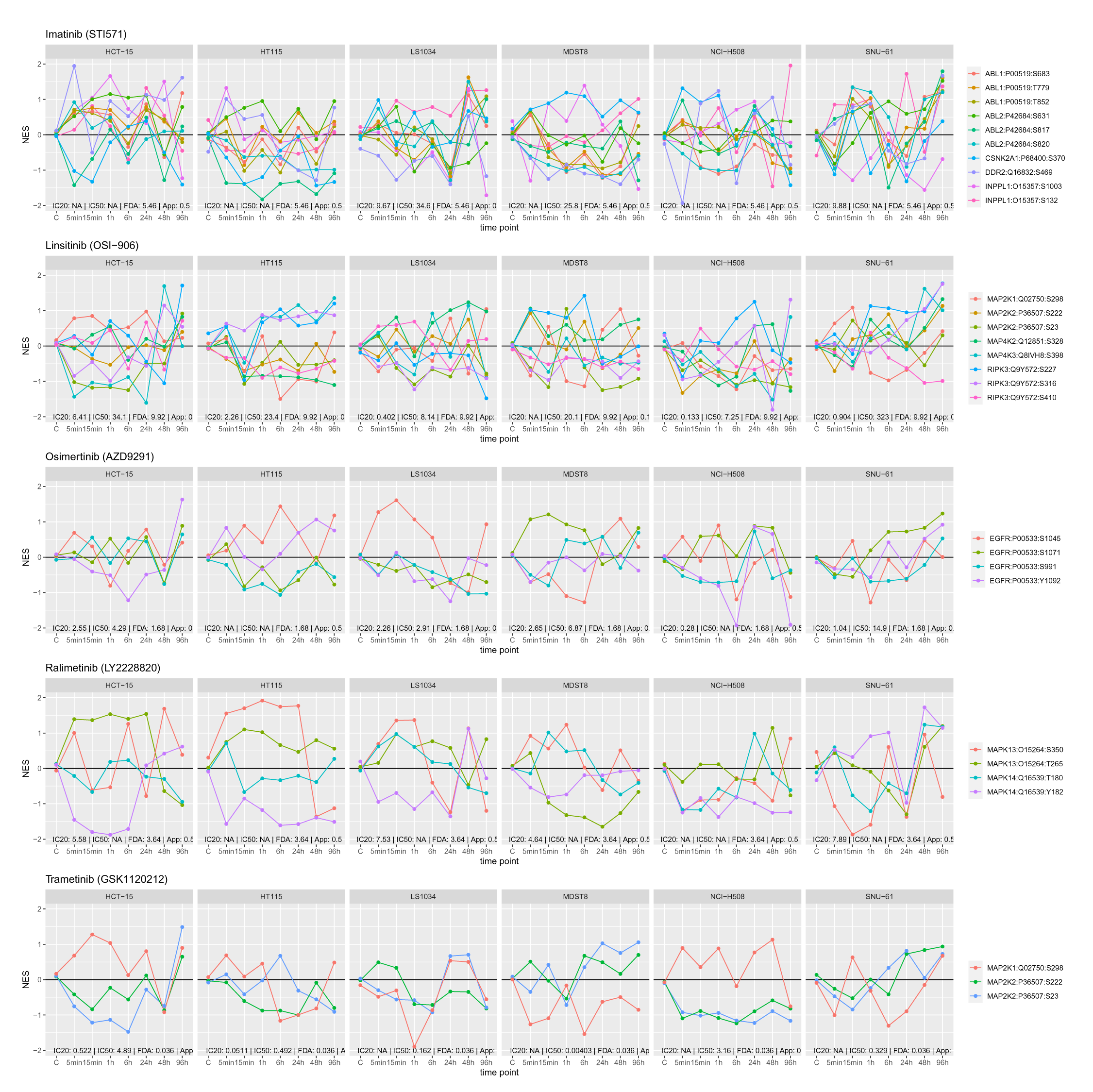
Temporal phosphosite-level VESPA perturbation profiles of known primary drug compound targets. The phosphosite-level VESPA activities are visualized for the top 5 downregulated known primary targets, grouped according to drug perturbations and cell lines.

**Supplementary Figure 15.**
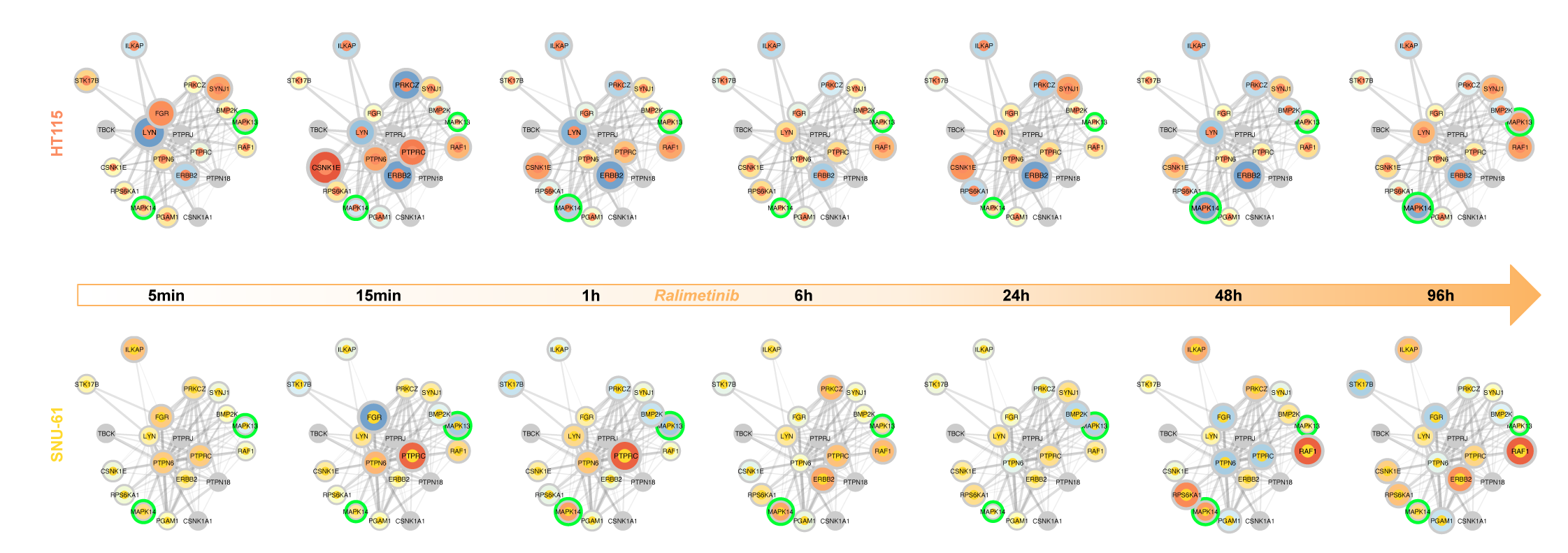
Context-specific wiring of signaling pathways. Visualization of network dysregulation and drug compound mechanism of action (MoA) for ralimetinib. Nodes indicate the most affected regulators with the inner circos colors indicating cell line type and the outer circos color and node size indicating VESPA activity. The edges indicate dysregulated, undirected interactions between the regulators (Methods). Line thickness indicates significance of dysregulation. Proteins highlighted in green indicate known primary and secondary targets.

**Supplementary Figure 16.**
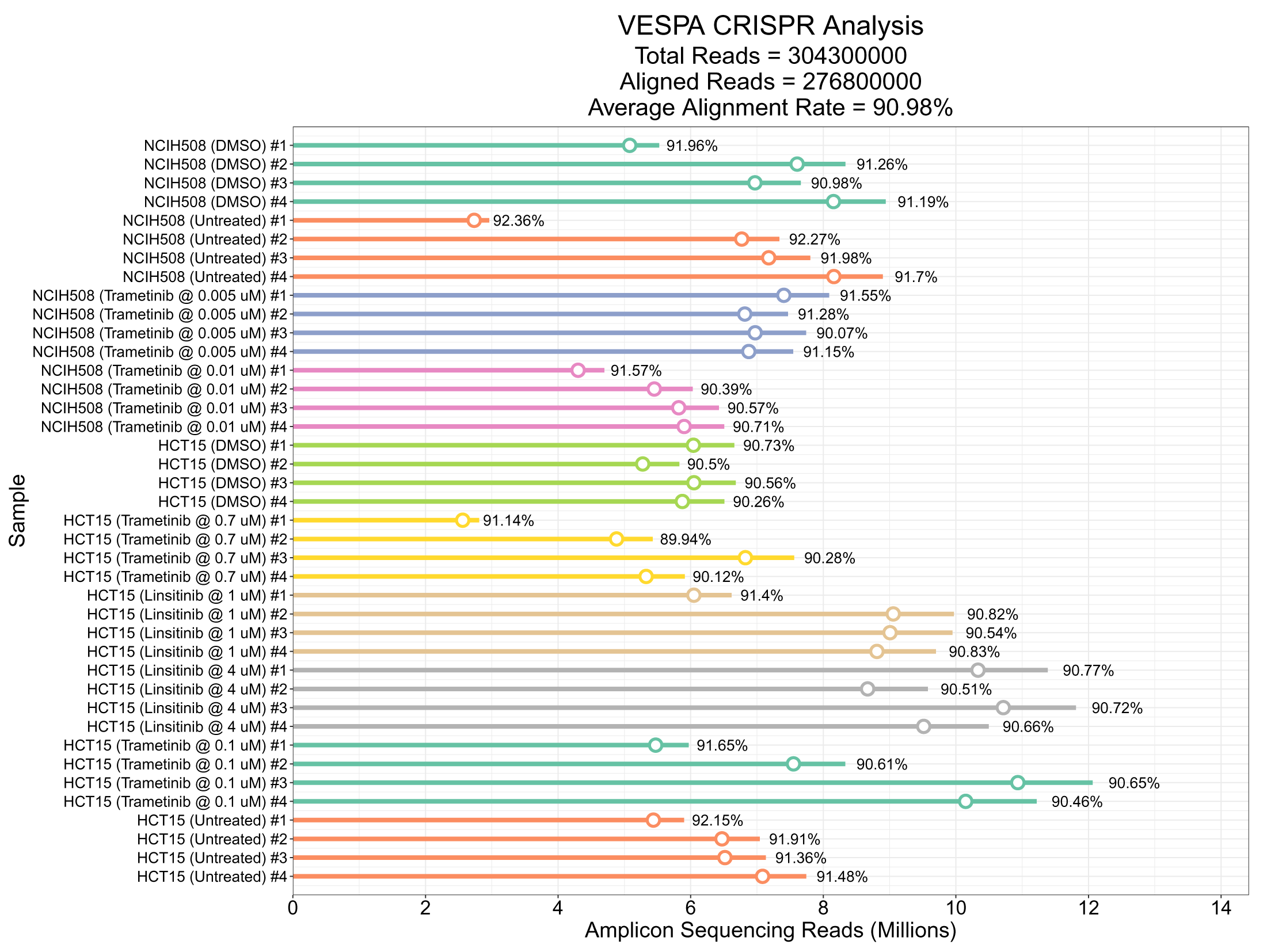
CRISPRko experiment technical assessment. The sgRNA alignment rate for each run is depicted separately.

**Supplementary Figure 17.**
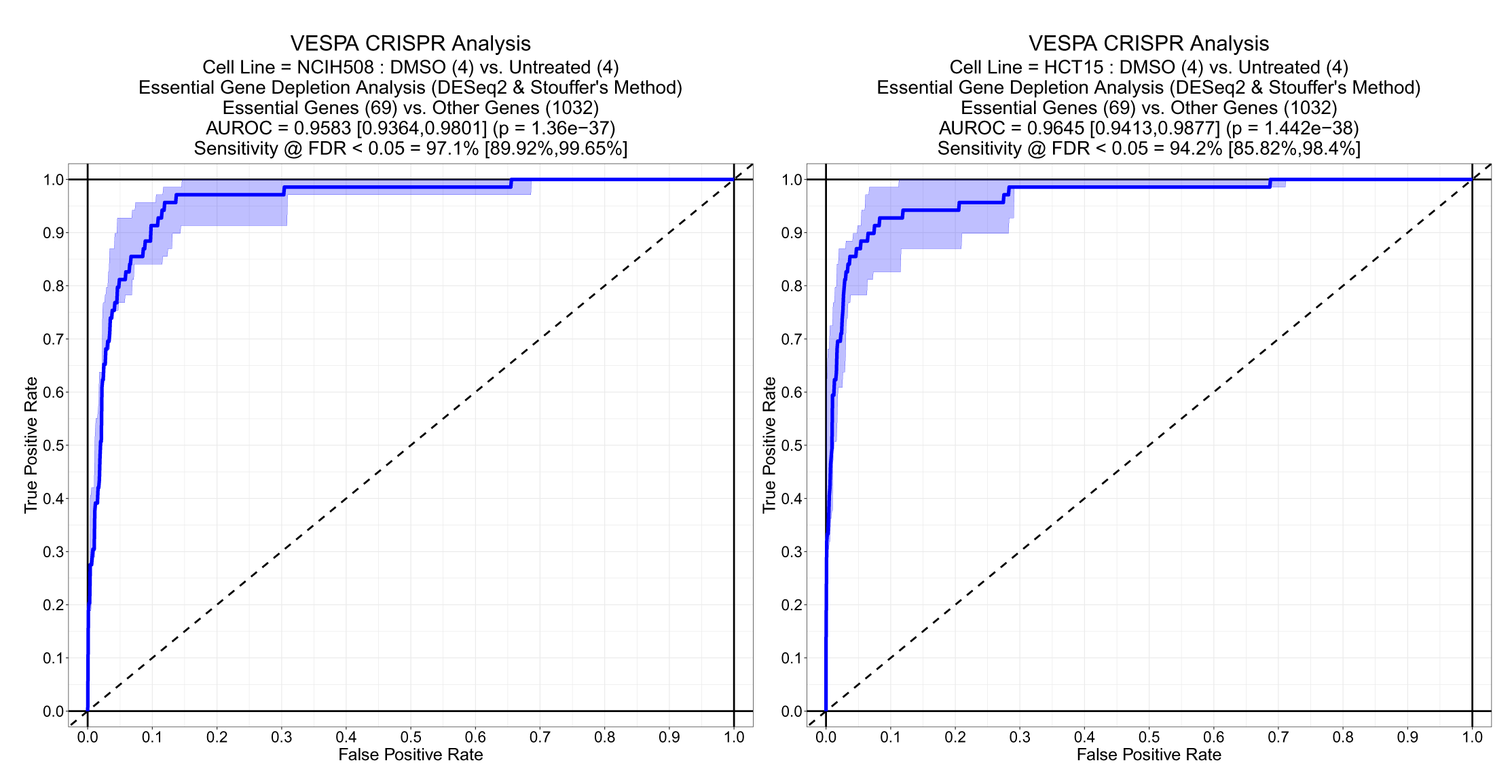
Receiver-operating-characteristics for the recovery of known essential genes in the CIRSPRko validation experiment. Comparing DMSO (last time point) *vs.* T0 (first time point) CRISPRko samples identified known essential genes in CRC with high accuracy for both NCI-H508 and HCT-15 cells (Methods).

**Supplementary Figure 18.**
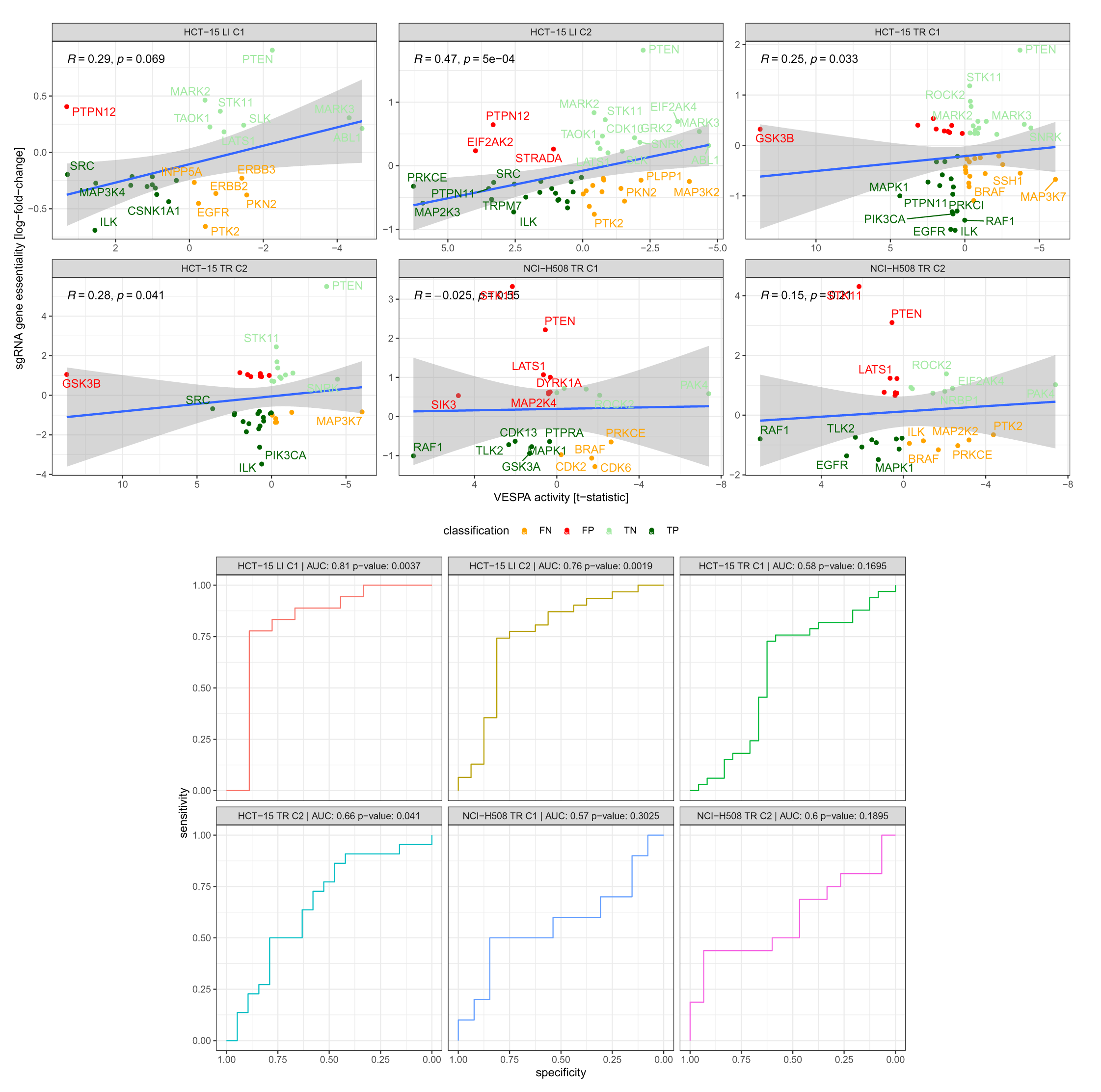
Correlation and receiver-operating-characteristics (ROC) for differential VESPA predictions of KP-enzymes involved in resistance mechanism against measured differential CRISPRko essential genes.

**Supplementary Figure 19.**
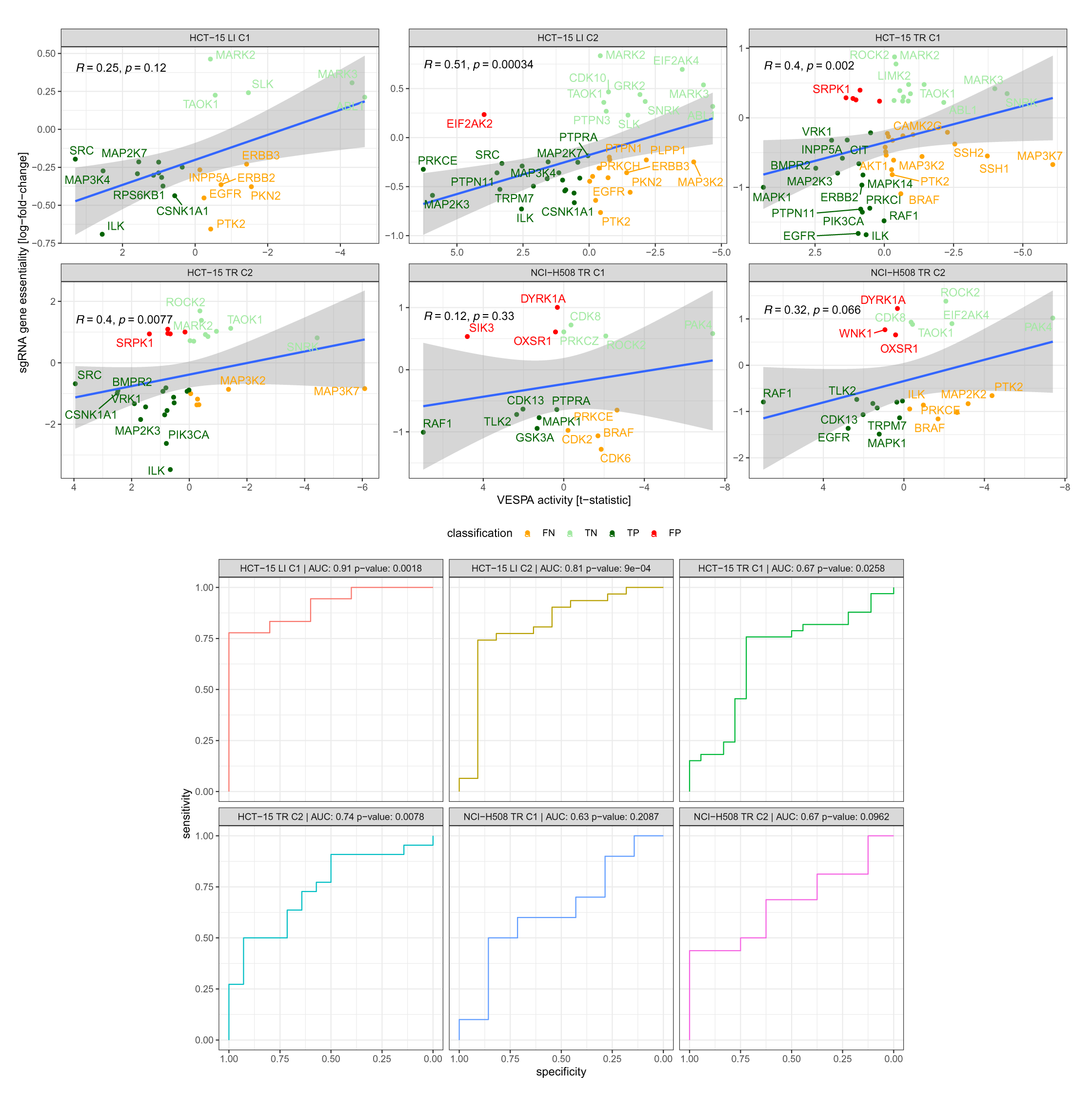
Correlation and receiver-operating-characteristics (ROC) for differential VESPA predictions of KP-enzymes involved in resistance mechanism against measured differential CRISPRko essential genes, excluding tumor suppressor genes.

## Notes

https://massive.ucsd.edu/ProteoSAFe/dataset.jsp?accession=MSV000091204

https://www.ncbi.nlm.nih.gov/geo/query/acc.cgi?acc=GSE224396

https://github.com/califano-lab/vespa

https://github.com/califano-lab/vespa.db

https://github.com/califano-lab/vespa.aracne

https://github.com/califano-lab/vespa.net

https://github.com/califano-lab/vespa.tutorial

